# Exchangeable HaloTag Ligands (xHTLs) for multi-modal super-resolution fluorescence microscopy

**DOI:** 10.1101/2022.06.20.496706

**Authors:** Julian Kompa, Jorick Bruins, Marius Glogger, Jonas Wilhelm, Michelle S. Frei, Miroslaw Tarnawski, Elisa D’Este, Mike Heilemann, Julien Hiblot, Kai Johnsson

## Abstract

We introduce exchangeable ligands for fluorescence labeling of HaloTag7 as an alternative to covalently bound probes. The exchangeable ligands open up new possibilities in imaging for a widely used labeling approach, including applications in points accumulation for imaging in nanoscale topography (PAINT), MINFLUX and live-cell, multi-frame stimulated emission depletion (STED) microscopy. We furthermore introduce orthogonal pairs of exchangeable ligands and HaloTags for dual-color PAINT and STED microscopy.

**Graphical Abstract:** 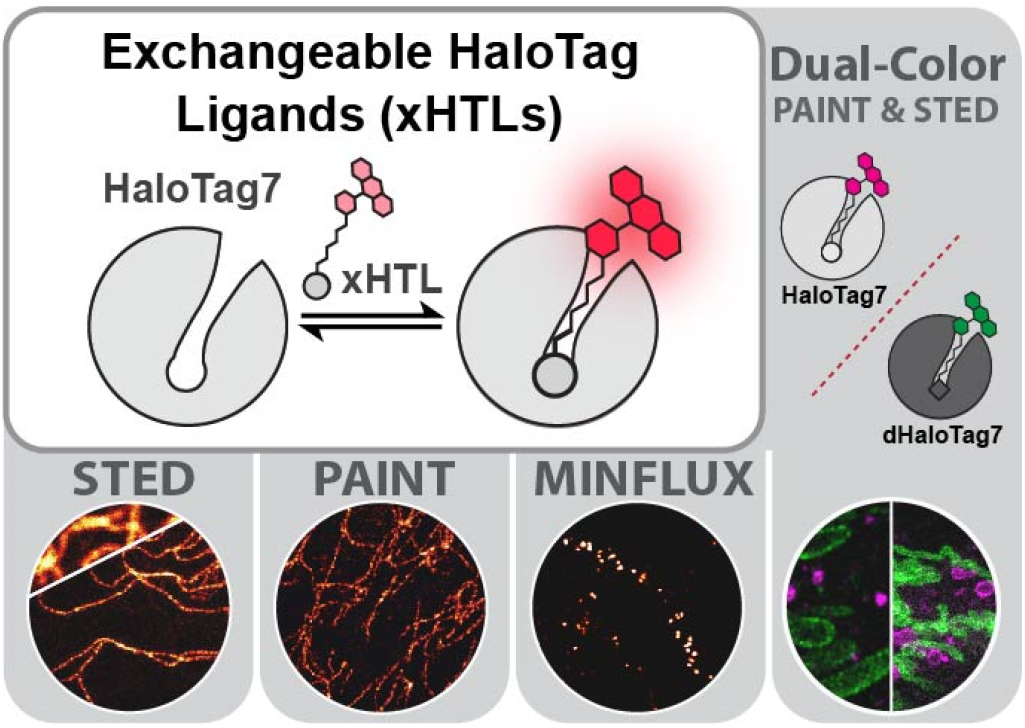

## Main

HaloTag7 is a powerful tool for live-cell imaging as it reacts covalently and specifically with various fluorescent ligands (HTLs; **Fig 1A**).^1^ The outstanding brightness and cell-permeability of the underlying synthetic fluorophores, usually rhodamines, make the approach particulary attractive for live-cell super-resolution microscopy techniques such as stimulated emission depletion (STED) microscopy.^2^ However, photobleaching limits multi-frame acquisition in STED microscopy. Exchangeable fluorophore labels that are constantly replenished from a large buffer reservoir provide an elegant way to reduce photobleaching.^3^ For example, non-covalent labeling approaches based on weak-affinity DNA hybridization show reduced photobleaching in STED microscopy due to continuous probe exchange.^4^ Transient and repetitive DNA-hybridization is also exploited to reach nanometer resolution in DNA points accumulation for imaging in nanoscale topography (DNA-PAINT) microscopy.^5^ However, exchangeable probes based on DNA are incompatible with live-cell imaging of intracellular targets.^6^ Other tools based on exchangeable probes, such as the fluorogen binder FAST^7^, are live-cell compatible but do not meet the demanding requirement of multiframe STED microscopy. Here we introduce exchangeable HaloTag7 Ligands (xHTLs, **Fig 1A**) that open up application of HaloTag7 for PAINT, MINFLUX^8^ and multi-frame, live-cell STED imaging.

**Figure 1.**
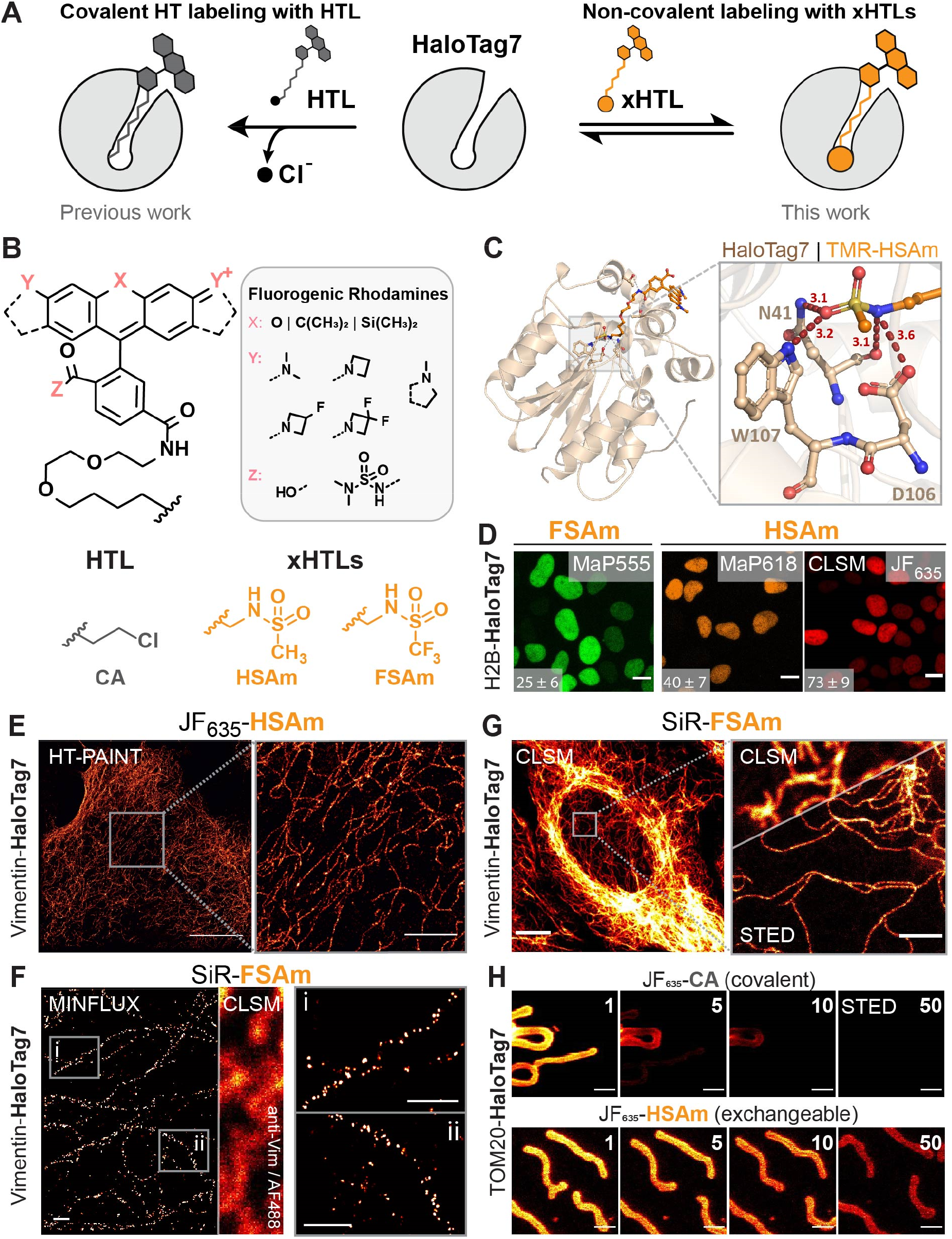
Exchangeable HaloTag Ligands (xHTLs) design, characterization and application in super-resolution fluorescence microscopy. **A.** Schematic representation of covalent HaloTag labeling with a fluorescent ligand (HTL, left) or non-covalent labeling with a fluorescent exchangeable ligand (xHTL, right). **B.** Chemical structure of the different (x)HTLs. Covalent HTLs are Dye-PEG_2_-C_6_-Cl (Chloroalkane, CA). Non-covalent xHTLs are Dye-PEG_2_-C_5_-methylsulfonamide (HSAm) and Dye-PEG_2_-C_5_-trifluoromethylsulfonamide (FSAm). Fluorogenic Rhodamines consist of tetramethyl rhodamine (TMR, X = O), carbo rhodamine (CRh, X = C(CH_3_)_2_) or silicon rhodamine (SiR, X = Si(CH_3_)_2_) cores with common modifications at the 3-carboxy (MaP-Dyes^9^, dimethylaminosulfonamide), exocylic amines (Janelia Fluor Dyes^10^, azetidine, 3,3-difluoroazetidine, 3,3-tetrafluoroazetidine) or xanthene-core (SiR700^11^, dihydroindoyl), **C.** Structural analysis of the TMR-HSAm/HaloTag7 complex. PDB-ID: 7ZJ0, 1.5 Å resolution. Protein structure is represented as cartoon, the ligands and residues are represented as sticks. Magnification on the binding pocket highlights polar interactions between HSAm and HaloTag7 residues. Distances in Å. **D.** Representative confocal laserscanning microscopy (CLSM) images of live U2OS cells expressing H2B-HaloTag7 labeled with the best dye/xHTL combinations (500 nM) over the visible spectrum. Scale bars: 10 μm. Bottom left corner, signal-over-background ratio from n ≥ 150 cells (mean ± S.E.M.). **E.** HaloTag-PAINT (HT-PAINT) image of a fixed U2OS cell endogenously expressing vimentin-HaloTag7^13^ labeled with JF_635_-HSAm (5 nM). Scale bar: 10 μm (overview) and 2 μm (magnified region). **F.** 2D-MINFLUX microscopy imaging of fixed U2OS vimentin-HaloTag7 cells with SiR-FSAm (2 nM). Vimentin immunostaining (AF488) and CLSM imaging was used as a reference. Magnification reveals vimentin-HaloTag7 molecules with a localization precision of ~4.0 nm. Intensities are represented in arbitrary units from 0 – 3 (overview) or 0 – 12 (magnified region). Scale bars: 0.2 μm. **G.** Comparison of confocal and STED images of vimentin-HaloTag7 in live U2OS cells labeled with SiR-FSAm. Same cell line as in E. Scale bars: 10 μm (overview) or 1 μm (magnified region). **H.** Multi-frame STED images of U2OS mitochondria outer membrane (TOM20-HaloTag7) labeled with JF_635_-CA or JF_635_-HSAm demonstrates improved apparent photostability of the exchangeable probes. Frame numbers indicated in the top right corner. Scale bar: 1 μm.

Taking advantage of the crystal structure of HaloTag7 labeled with tetramethylrhodamine (TMR), we used computational screening (**Fig. S1**) to identify methylsulfonamide (HSAm) and trifluoromethylsulfonamide (FSAm, **Fig. 1B**) derivatives linked to rhodamines as non-covalent ligands (xHTLs) with nanomolar dissociation constants and fast binding kinetics (k_on_ >10^6^ M^-1^ s^-1^) (**Fig. S2, Table S1**). The crystal structures of HSAm and FSAm linked to TMR (**Fig. 1C, S3, S4 & Table S2**) in complex with HaloTag7 revealed that the sulfonamides occupy the active site of HaloTag7, while TMR binds to the same site as in covalently labeled HaloTag7. FSAm and HSAm derivatives can be combined with various fluorogenic rhodamines^9–11^ in order to create cell-permeable xHTL probes spanning almost the entire visible spectrum (λ_ext_ = 525 – 670 nm). In most cases, the brightness of the xHTLs probes bound to HaloTag7 *in vitro* are comparable to those of the corresponding covalently bound probes (**Table S1**). Some of the xHTLs such as SiR-FSAm displayed comparable fluorogenicity upon binding to HaloTag7 as reported for the corresponding chloroalkanes (**Fig. S5, Table S3**).^12^ *In vitro*, FSAm derivatives bind with higher affinity to HaloTag7 than the corresponding HSAm derivatives. However, their relative performance in cellular staining depends on the attached rhodamine. FSAm worked best combined with MaP555 and JF_525_, whereas HSAm offered the highest cellular brightness with adequate signal-over-background ratios in combination with far red dyes (λ_ext_ ≥ 610 nm) such as MaP618 or JF_635_ (**Fig. 1D & S6**). Furthermore, various subcellular targets could be stained using SiR-FSAm which showed faster labeling kinetics in live U2OS cells than the covalent SiR-HTL labeling (**Fig. S7**). The exchangeable character of the xHTLs binding to HaloTag7 enabled repetitive labeling and washing of nuclear localized HaloTag7 (NLS-HaloTag7) in U2OS cells with SiR-HSAm within minutes (**Fig. S7D**).

The fast bindings kinetics and specific labeling of the xHTLs FSAm and HSAm enabled the spatio-temporal separation of binding events required for PAINT microscopy (**Table S4**).^5^ Using JF_635_-HSAm, we imaged vimentin-HaloTag7^13^ (intermediate filaments) in fixed U2OS cells by PAINT microscopy (**Fig. 1E**), further referred as HaloTag-PAINT (HT-PAINT), with a resolution comparable to that achieved by DNA-PAINT (~32 nm, **Fig S.8**).^14^ We further hypothesized that xHTL transient staining could be compatible with MINFLUX imaging which synergistically combines sparse labeling (as performed in HT-PAINT) and a patterned excitation beam featuring an intensity minimum (as in STED) to routinely reach single-digit nm resolution.^8^ Using SiR-FSAm, vimentin-HaloTag7 molecules could be imaged in 2D MINFLUX with ~ 4.0 nm localization precision by using 220 photons in the last iteration step (L = 40 nm) (**Fig. 1F & S9**).

Next, a panel of different fluorogenic xHTLs (MaP555-FSAm, MaP618-HSAm, JF_635_-HSAm, SiR-HSAm and SiR-FSAm) were characterized in live-cell STED microscopy (**Fig. 1G & S10**). For example, labeling vimentin-HaloTag7^13^ of U2OS with SiR-FSAm resulted in similar intermediate filament diameters as covalent HaloTag7 labeling. However, xHTLs staining was significantly less susceptible to photobleaching compared to covalent labeling. Multi-frame STED imaging of mitochondria in live U2OS cells (TOM20-HaloTag7, outer mitochondrial membrane) revealed that transient labeling with JF_635_-HSAm allowed to trace dynamic objects over 50 consecutive frames, while covalent labeling with JF_635_-HTL limited the imaging to 10 frames (**Fig. 1H**). Transient labeling with xHTLs increased the STED-image frame numbers with > 50% signal intensity by 3 to 5-fold compared to covalent labeling for all fluorophores tested (MaP555, MaP618 and SiR, **Fig. S10**). This increase in apparent photostability makes xHTLs attractive probes for multi-frame imaging.

Motivated by the performance of xHTLs with HaloTag7, we developed a second reversible labeling system with orthogonal specificity using an HaloTag7 mutant in which active-site residue Asp106 is mutated to Ala (dHaloTag7)^15^ (**Fig. 2A**). dHaloTag7 shows only micromolar affinities for HSAm-based probes (**Fig. S11A**), but displays nanomolar affinities and fast binding kinetics with primary alcohols derivatives with different alkane chain length (Hy4 and Hy5; **Fig. 2B**, **Fig. S11, Table 1**). Hy4 and Hy5 coupled to appropriate fluorophores also possess fluorogenicity upon dHaloTag7 binding (**Fig. S11E**). Structural analysis of the complex of TMR-Hy5 and dHaloTag7 revealed the presence of a structural chloride ion and a water molecule in the binding site (**Fig. 2C & S12**). This difference in binding modes of Hy5 and HSAm explains their orthogonality, even though the fluorophores are bound in an identical manner to the surface of both proteins. Hy5-based xHTLs showed higher binding affinity than the corresponding Hy4-based xHTLs, but their relative performance in live-cell imaging experiments also depends on the nature of the attached rhodamine. For MaP555, only coupling to Hy5 allowed selective staining of H2B-dHaloTag7, whereas for red shifted fluorophores such as SiR both Hy5- and Hy4-derivatives showed highly specific labeling (**Fig. 2D & S13A**). The orthogonal binding of spectrally distinct xHTLs-based probes to HaloTag7 and dHaloTag7 enabled to perform live cell dual-color imaging, with best results being obtained by combining SiR-HSAm and MaP555-Hy5 (**Fig. S13B**). A four-color confocal image of live U2OS cells was thus obtained using HaloTag7 labeled with SiR-HSAm, dHaloTag7 labeled with MaP555-Hy5, SNAP-tag labeled with BG-JF_585_ and SiR700-actin^16^ (**Fig. 2E**).

**Figure 2.**
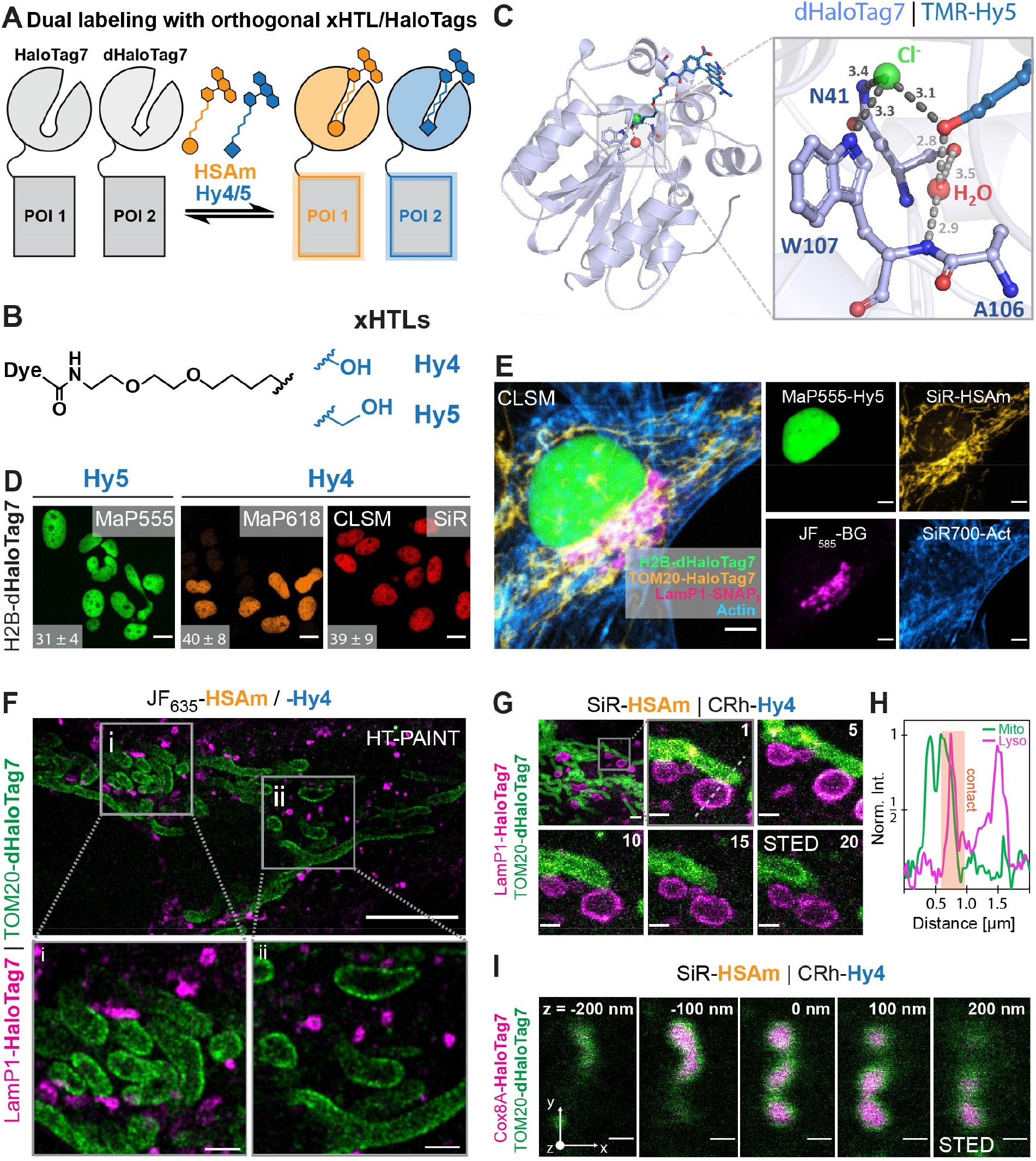
Orthogonal xHTL probes for super-resolution microscopy in dual-color. **A.** Schematic illustration of orthogonal xHTL combination with HaloTag7- and dHaloTag7-tagged proteins of interest (POIs) for dual-color staining. dHaloTag7 = HaloTag7-D106A. **B.** Chemical structure of xHTLs specific for dHaloTag7 consisting of Dye-PEG_2_-C_n_-R. R = Hydroxy, n = 4 (Hy4) or 5 (Hy5). **C.** Structural analysis of the TMR-Hy5/dHaloTag7 complex. PDB-ID: 7ZIZ, 1.5 Å resolution. Protein structure is represented as cartoon, the ligands and residues are represented as sticks. Magnification on the binding pocket highlights polar interactions between Hy5 and dHaloTag7 residues. Structural water and chloride ion are represented as spheres. Distances in Å. **D.** Representative confocal laser-scanning microscopy (CLSM) images of live U2OS cells expressing H2B-HaloTag7 labeled with the best dye/xHTL combinations over the visible spectrum. Scale bars: 10 μm. Bottom left corner, signal-over-background ratio from n ≥ 150 cells (mean ± S.E.M.). **E.** Four-color CLSM image of a live U2OS cell stained using orthogonal xHTLs (MaP555-Hy5, SiR-HSAm), BG-JF_585_ and a SiR700-actin probe^16^ to label dHaloTag7-H2B (nucleus), TOM20-HaloTag7 (mitochondria surface) and LamP1-SNAP_f_-tag (lysosome). Scale bar: 5 μm. ‘Hot’ lookup tables were applied. **F**. Dual-target exchange HT-PAINT image of mitochondria and lysosomes of fixed U2OS cells using orthogonal xHTLs. Cells co-expressing TOM20-HaloTag7 and LamP1-dHaloTag7 via T2A fusion are sequentially labeled and imaged using JF_635_-HSAm (5 nM, magenta) and JF_635_–Hy4 (3 nM, green), respectively. Scale bars: 10 μm (overview) or 1 μm (magnified region). **G.** Dual-color time-lapse STED image of mitochondria-lysosome dynamics in live U2OS cells. xHTLs were applied in 500 nM concentrations and imaged over 20 consecutive frames at an imaging speed of 2 frames/minute in a 10 μm^2^ area. Frame numbers indicated in the top left corner. SiR- and CRh-xHTLs were chosen for their higher brightness in STED imaging. Scale bars: 2 μm (overview) or 0.5 μm (magnified region). **H.** Line scan profile along lysosomal vesicle and mitochondrial revealing contact-site at high resolution. **I.** 3D-STED images of U2OS mitochondria outer membrane and inner matrix membrane labeled with xHTLs. Cells express Cox8A-HaloTag7 (matrix, mtx) and TOM20-dHaloTag7 (outer-membrane, otm) via T2A fusion and were labeled with SiR-HSAm and CRh-Hy4 (500 nM). STED imaging in a volume of 15.6 μm^3^ (x = 2.44 μm, y = 3.20 μm, 40-times z = 50 nm stacks). z-plans indicated in the top right corner. Scale bar: 0.5 μm.

In PAINT microscopy, JF_635_-Hy4 offered binding kinetics that allowed resolving vimentin-dHaloTag7 expressed in U2OS cells with similar quality as obtained with JF_635_-HSAm and HaloTag7 (**Fig. S14**). In livecell STED microscopy, MaP555-Hy5, MaP618-Hy4, JF_635_-Hy4 and SiR-Hy5 allowed imaging of mitochondria (TOM20-dHaloTag7) over significantly higher frame numbers than what was possible using covalently labeled HaloTag7 (up to 73 frames with > 50% signal intensity for SiR-Hy4, **Fig. S14**). Dual-target super-resolution imaging was demonstrated by an Exchange-PAINT approach^17^ in fixed U2OS cells by sequentially imaging mitochondria (TOM20-dHaloTag7) using JF_635_-Hy4 and then either lysosomes (LamP1-HaloTag7) or the endoplasmic reticulum (CalR-HaloTag7-KDEL) using JF_635_-HSAm (**Fig. 2F & S15A**). 2-color STED microscopy with a single depletion laser was used for multi-frame super-resolution co-imaging of lysosomes (LamP1-HaloTag7) and mitochondria (TOM20-dHaloTag7) in live U2OS cells labeled with SiR-HSAm and CRh-Hy4, respectively. This enabled the detection of dynamic mitochondria-lysosome contact sites at super-resolution over 20 frames (**Fig. 2G&H, S15**). The possibility to follow this dynamic interaction at high spatio-temporal resolution should help to clarify its role in multiple human pathologies.^18^ xHTLs also allowed the live-cell 3D STED imaging of the mitochondrial surface (TOM20-dHaloTag7) and matrix (Cox8A-HaloTag7), respectively labeled with SiR-HSAm and CRh-Hy4, in a volume of 15.6 μm^3^ (50 nm z-stacks, **Fig. 2I**).

In conclusion, we introduce fluorescent and exchangeable HaloTag ligands (xHTL) for HaloTag7 that are ideally suited for PAINT, MINFLUX and multi-frame, live-cell STED microscopy. Furthermore, both PAINT and STED microscopy can be performed in dual color using the orthogonal pairs of xHTLs together with HaloTag7 and dHaloTag7. The fluorescent xHTL probes thus open up a set of new imaging applications for a popular labeling strategy.

## Supporting information

Supplementary methods, figures and tables for Exchangeable HaloTag Ligands (xHTLs) for multi-modal super-resolution fluorescence microscopy

## Methods

Experimental procedures are described in supplementary information.

## Data availability

The X-ray crystal structures obtained for this study have been deposited to the PDB with the following deposition codes: HaloTag7/HSAm-TMR (7ZJ0), HaloTag7/TMR-FSAm (7ZIY), dHaloTag7/Hy5-TMR (7ZIZ). Key plasmids will be deposited at Addgene. Requests for data will be fulfilled by the corresponding authors.

## Acknowledgements

This work was supported by the Max Planck Society, the Ecole Polytechnique Federale de Lausanne (EPFL) and the Deutsche Forschungsgemeinschaft (DFG, German Research Foundation), TRR 186 and SFB1177; INST 161/778-1 FUGG. J.K. was supported by the Heidelberg Biosciences International Graduate School (HBIGS). The authors thank I. Schlichting for X-ray data collection. Diffraction data were collected at the Swiss Light Source, beamline X10SA, of the Paul Scherrer Institute, Villigen, Switzerland. The authors thank B. Koch, A. Bergner, V. Nasufovic, B. Réssy and D. Schmidt (all MPIMF) for providing reagents or material. We thank the mass spectrometry (S. Fabritz, T. Rudi and J. Kling) facility of the MPIMR for its support. We thank S. Jakobs (MPINat) for providing the U2OS vimentin-HaloTag7 cells and M. Lima (MPIMF) for providing a MINFLUX data analysis workflow.

## Author contributions

J.K. and J.B. designed exchangeable HaloTag Ligands. J.K. performed biochemical characterization and cellular applications of xHTLs. M.G. and M.H performed HT-PAINT. J.W. performed stopped-flow binding kinetics. E.D. provided supervision for STED and MINFLUX imaging. M.T. solved the crystal structures. J.K. and J.H. analyzed the crystal structures. M.F. generated stable cell lines. K.J. and J.H. supervised the work. J.K, J.H. and K.J. wrote the manuscript with input from all authors.

