## Supplementary methods, figures and tables for Exchangeable HaloTag Ligands (xHTLs) for multi-modal super-resolution fluorescence microscopy for "Exchangeable HaloTag Ligands (xHTLs) for multi-modal super-resolution fluorescence microscopy"

I HSAm / FSAm-xHTL 6
II Hy4 / Hy5-xHTL 9
III Deprotection 10
IV Fluorophore coupling 11

Molecular biology & Biochemistry ........ 14

Plasmids 14
Protein production and purification 14
Covalent labeling of HaloTag7 protein and in-gel fluorescence scan 15
Protein crystallization and X-ray diffraction data collection 15
Fluorophore concentrations 16
Water-Dioxane titration 17
Quantum yield 17
Extinction coefficients 17
Fluorescence increase upon protein binding assay 18
Affinity measurement by fluorescence polarisation assays 18
Stopped-Flow kinetic fluorescence polarisation assay 19

Cell biology and microscopy ........ 20

Stable cell-line establishment 20
Cell fixation and staining 20
Live-cell staining 20
Confocal Fluorescence Microscopy 21
Wide-field Fluorescence Microscopy 22
PAINT Imaging 22
PAINT-Image processing and determination of single-molecule binding kinetics 23
MINFLUX Microscopy 24
Live-cell STED Microscopy 24

**SUPPLEMENTARY FIGURES ........ 30**

Figure S. 1 | Computational screening of xHTLs candidates. 30
Figure S. 2 | *In-vitro* characterization of xHTL candidates. 31
Figure S. 3 | Structural analysis of the TMR-HSAm/HaloTag7 complex. 32
Figure S. 4 | Structural analysis of the TMR-FSAm/HaloTag7 complex. 33
Figure S. 5 | Spectral and fluorogenic properties tuning of xHTLs. 34
Figure S. 6 | Live-cell staining characterization of xHTLs. 35
Figure S. 7 | Applications of xHTLs for live-cell confocal microscopy. 36
Figure S. 8 | Comparison of HT-PAINT and DNA-PAINT. 37
Figure S. 9 | Characterization of xHTLs performance in MINFLUX microscopy. 38
Figure S. 10 | STED photobleaching resistance of xHTLs compared to covalent HTLs. 39
Figure S. 11 | Development of an orthogonal exchangeable xHTL/HaloTag7 protein pair. 41
Figure S. 12 | Structural analysis of the TMR-Hy5/dHaloTag7 complex. 42
Figure S. 13 | Dual-color confocal images using orthogonal xHTLs-(d)HaloTag7 pairs. 43
Figure S. 14 | Super-resolution fluorescence microscopy enabled by Hy4/5 and dHaloTag7. 44
Figure S. 15 | Dual-color super-resolution microscopy using xHTLs. 45

**SUPPLEMENTARY TABLES ........ 46**

Table S. 1 | Data collection and refinement statistics for the crystal structure. 47
Table S.2 | Fluorescence intensity increase of fluorogenic xHTL probes upon target binding. 48
Table S.4 | Single-molecule binding kinetics and image resolution in HT-PAINT 48

### Supplementary Methods

#### Molecular docking

The crystal structure of HaloTag7 fully reacted with TMR-CA (PDB-ID 6Y7A)[1] was prepared in the Prime module of the Schrodinger 2020-1 package. Default settings (pH value 7.4) were used including removal of crystallization ligands and water molecules. All ligands were sketched with the 2D Sketcher module and diversified with the R-group creator and enumeration tool based on the TMR-CA lead-structure (**Fig. S1B**). Resulting compound libraries were processed in the Schrodinger ligprep module to obtain their 3D-geometry and most probable ionization states (Epik module, pH 7.4 ± 2.0). A cubic 800 $nm^3$ docking grid was generated around the HaloTag7 ligand molecule (reacted TMR-CA) and subsequent docking was performed using Glide[2] in standard precision configuration yielding up to 10 docking poses per ligand. During the docking process, polar interactions with the amino acids N41, D106 and W107 were applied as constrains.

Initial docking results were filtered by their glide score (gscore), which rewards favorable interactions and penalizes steric clashes using a static protein model. Docking results were assessed by their predicted binding energy (*i.e.* $\Delta G_{bind}$, Prime MM-GBSA module) using the zwitterionic open form of TMR ($TMR_o$, **Fig. S1D**). Similarly, the predicted cell-permeability *i.e.* membrane energy[3] ($\Delta G_{memb}$, qikprop & Prime MM-GBSA module) was calculated from the closed TMR spirolactone form of tentative xHTLs ($TMR_c$, **Fig. S1D**), due to the incompatibility with charged compounds of this method. Both descriptors were evaluated in comparison to TMR-CA ($\Delta\Delta G$ = $^{TMR\text{-}CA}\Delta G$ – $^{TMR\text{-}candidate}\Delta G$) and for binding energies, a re-docking of TMR-CA was used as reference. For high accuracy, $\Delta G_{bind}$ was calculated from induced-fit docking[4] with a final subset of candidates, giving a higher degree of flexibility to residues located in the ligand area: W107, F149, F205, P206, L209. Maestro was used for evaluation of favourable protein/ligand interactions and PyMOL[5] software for presentation of protein/ligand poses.

#### Chemical Procedures

All chemical reagents and solvents for synthesis were purchased from commercial suppliers (Acros, Fluka, Merck KGaA, Roth, Sigma-Aldrich, TCI, TOCRIS) and used without further purification. Water-free solvents were stored over molecular sieves and used directly from a sealed-bottle. 6-Carboxy rhodamine dyes were either prepared according to literature procedures by V. Nasufovic, B. Réssy, D. Schmidt (MPImR, Heidelberg) or purchased from commercial sources (Sigma-Aldrich, AAT Bioquest, **Method Table 1**).

Reactions performed under air and moisture exclusion were carried out in heat-dried glassware and under inert argon atmosphere using Schlenk techniques. Evaporation *in vacuo* was achieved at 40 °C and 10 – 850 mbar at a rotary evaporator (Heidolph) or by lyophilization on a lyophilizer (Christ) equipped with a vacuum pump (Vacuubrand). Reaction progress was monitored by analytical thin-layer chromatography (TLC) or liquid chromatography coupled to mass spectrometry (LC-MS). TLC was accomplished on commercially available $SiO_2$-plates (POLYGRAM® SIL G/UV254, 0.2 mm layer pre-coated polyester sheet, 40 x 80 mm, Roth) in appropriate solvent mixtures and visualized by using 254/366 nm UV-light or staining solutions ($KMnO_4$, Ninhydrin, Vanillin) and gentle heating. LC-MS was performed on a LCMS2020 (Shimadzu) connected to a Nexera UHPLC system.

Column: $C_{18}$ 1.7 µm, 50 × 2.1 mm (ACQUITY UPLC BEH, Waters). Buffer A: 0.1% Formic acid (FA)/MiliQ® water ($ddH_2O$), buffer B: acetonitrile (MeCN). Typical gradient was from 10% to 90% B within 6 min with 0.5 mL $min^{-1}$ flow.

**Method Table 1.** Source and spectral properties according to literature of fluorophores used in this work.

| Fluorophore | reference | source | $\lambda_{abs}$ [nm] | ε [$M^{-1}\cdot cm^{-1}$] |
|---|---|---|---|---|
| $JF_{525}$ | Grimm *et al.* (2017)[6] | I | 525 | 138,000$^b$ |
| TMR | Mudd *et al.* (2015)[7] | AAT Bioquest | 555 | 89,000$^a$ |
| MaP555 | Wang *et al.* (2020)[8] | I | 558 | 142,000$^b$ |
| $JF_{585}$ | Grimm *et al.* (2017)[6] | I | 585 | 52,500$^b$ |
| CRh | Butkevich *et al.* (2016)[9] | AAT Bioquest | 616 | 152,000$^b$ |
| MaP618 | Wang *et al.* (2020)[8] | Ii | 616 | 5,500$^b$ |
| $JF_{635}$ | Grimm *et al.* (2017)[6] | Sigma-Aldrich | 635 | 17,000$^b$ |
| SiR | Lukinavičius *et al.* (2013)[10] | AAT Bioquest | 646 | 120,000$^b$ |
| $JF_{646}$ | Grimm *et al.* (2015)[11] | I | 646 | 106,000$^b$ |
| SiR700 | Lukinavičius *et al.* (2016)[12] | I | 687 | 100,000$^b$ |

**a** In PBS pH 7.4, **b** in activity buffer + 0.1% SDS, **c** in ethanol + 0.1% TFA.
**i** in-house synthesis by B. Réssy / D. Schmidt, **ii** in-house synthesis by V. Nasufovic.

Flash column purification was performed using a Biotage (Isolera™ One) flash system equipped with pre-packed $SiO_2$ columns (SiliaSep™ Flash Cartridges, 40 – 63 µm, 60 Å). Depending on the batch size 12 g, 25 g or 40 g columns with 37, 75 or 100 mL $min^{-1}$ flow rate were used. Typical gradients were 10 to 50% ethyl acetate in n-hexane or 1 to 10% methanol in dichloromethane (DCM) within 10 column volumes (CV). Small-scale preparative reversed-phase high-performance liquid chromatography (RP-HPLC) was carried out on an UltiMate 3000 system (Thermo Fisher Scientific). Column: $C_{18}$ 5 µm, 21.2 × 250 mm (Supelco). Buffer A: 0.1% TFA in MiliQ® water, buffer B: MeCN. Typical gradient was from 20% to 90% B within 45 min with 8 mL $min^{-1}$ flow. A 2998 PDA detector allows automated product collection based on the absorption wavelength of fluorescent labels (at 280, 550, 620 or 650 nm, respectively). Large-scale RP-HPLC-MS purification (> 3 mg) was carried-out with a LCMS-2020 unit (Shimadzu) coupled with a prominence LC-20AP UFLC (Shimadzu). Column: $C_{18}$ 5 µm, 30 × 250 mm (Shimadzu). Buffer A: 0.1% FA in $ddH_2O$, buffer B: MeCN. Typical gradient was from 10% to 90% B within 45 min with 20 mL $min^{-1}$ flow. UV/Vis absorption was recorded using a SPD-M20A UV-VIS photodiode array detector and the desired compounds were collected based on the calc. mass-to-charge ratio (m/z) using a DUIS-2020 dual ion source (Shimadzu).

$^1$H-, $^{13}$C- and $^{19}$F-NMR spectra were recorded in deuterated solvents on a *Bruker Avance* III HD 400 NMR spectrometer at 298 K at 400 MHz ($^1$H), 101 MHz ($^{13}$C) or 377 MHz ($^{19}$F), respectively. Chemical shifts are expressed as parts per million (ppm, δ) and referenced to the solvent signals ($^1$H / $^{13}$C) as internal standards: $CDCl_3$ (7.26 / 77.16 ppm), $CD_3OD$ (3.31 / 49.00 ppm), DMSO-$d_6$ (2.05 / 39.52) and MeCN-$d_3$ (1.94 / 118.26 ppm) while $^{19}$F-signals were not referenced. Coupling constants J are reported in Hz. Signal descriptions include: s = singlet, d = doublet, t = triplet, q = quartet, p =pentet, s – sextet, h – heptet and m = multiplet. $^1$H-qNMR was performed using dry 1,4-dioxane (Acros, 3.57 ppm in DMSO-$d_6$, 8H) as a reference standard to quantify fluorophore concentrations. HRMS validation of synthesized chemical compounds was performed on a Bruker maXis II™ ETD spectrometer with electron spray ionization (ESI) by the *Mass Spectrometry* facility (MPImR, Heidelberg).

**I HSAm / FSAm-xHTL**

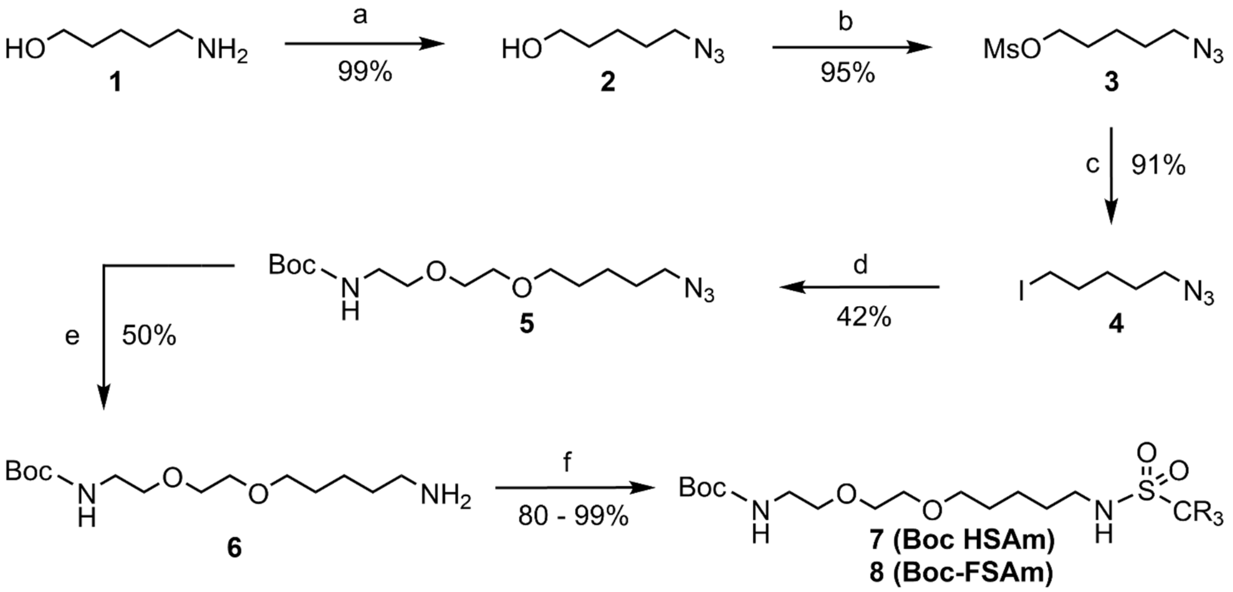

**Method Figure 1. Synthetic route to xHTL precursor Boc-HSAm (7) and Boc-FSAm (8).**
**a.** Imidazole-1-sulfonyl azide · HCl, $K_2CO_3$, Cu(II)$SO_4$ · 5 $H_2O$, MeOH, 12 h, 0 °C to room temperature (rt). **b.** MsCl, $NEt_3$, DCM, 3 h, 0 °C to rt. **c.** NaI, acetone, o/n, rt. **d.** BocNH-PEG$_2$-OH, NaH, THF/DMF, 3 h, 0 °C to rt. **e.** $PPh_3$, THF/H2O, 48 h, rt **f.** MsCl or TfCl, $NEt_3$, DCM, 3 h, 0 °C to rt. R: $CH_3$ (HSAm), $CF_3$(FSAm). Boc: -COO-C($CH_3$)$_3$.

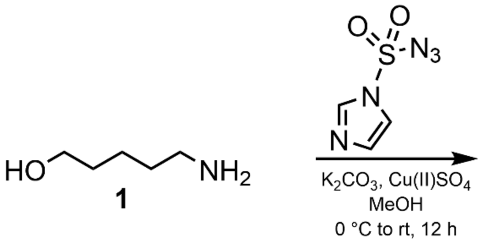

**5-Azido-1-pentanol (2):** The diazo-transfer reagent imidazole-1-sulfonyl azide hydrochloride was obtained in 74% yield according to the protocol of Goddard-Borger *et al.* 2007.[13] **$^1$H NMR** (400 MHz, $CDCl_3$): δ 7.47 (s, 1H), 5.97 (s, 1H), 5.64 (s, 1H).

A suspension of 5-amino-1-pentanol (**1**, 1.3 mL, 13 mmol, 1 eq.), $K_2CO_3$ (4.1 g, 29.1 mmol, 2.25 eq.) and Cu(II)$SO_4$ · 5 $H_2O$ (32 mg, 0.13 mmol, 1 mol%) was prepared in a heat-dried Schlenk-flask in 5 mL dry methanol (MeOH) under a stream of argon. The solution was cooled to 0 °C and imidazole-1-sulfonyl azide · HCl (3.3 g, 15.5 mmol, 1.2 eq.) was added portion-wise. The mixture was left stirring for 12 h at rt, concentrated under reduced pressure and acidified with conc. HCl. The solid was taken up in 200 mL ethyl acetate (EtOAc), washed with 100 mL $H_2O$ and brine, dried over $MgSO_4$, filtered and evaporated to afford the crude product as a pale-yellow oil. Flash column chromatography (5 to 70% EtOAc in n-hexane for 10 CV) afforded compound **2** (1.7 g, 13 mmol, 99%) as a pale-yellow oil. **$^1$H NMR** (400 MHz, $CDCl_3$): δ 3.63 (t, J = 6.4 Hz, 2H), 3.26 (t, J = 6.9 Hz, 2H), 1.59 (m, 4H), 1.44 (m, 2H). **$^{13}$C NMR** (101 MHz, $CDCl_3$): δ 62.24, 51.38, 32.10, 28.62, 22.98. **HRMS** (m/z): $[M + H]^+$ calcd. for $C_5H_{12}N_3O^+$, 130.0975; found, 130.0974.

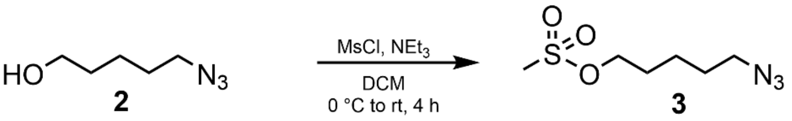

**5-Azidopentyl-1-methansulfonate (3)**: A heat-dried round-bottom flask was charged with 65 mL dry DCM and triethylamine ($NEt_3$, 2.7 mL,19.5 mmol, 1.5 eq.). **2** (1.7 g, 13 mmol, 1 eq.) was added and the mixture was cooled to 0 °C. Methanesulfonylchloride (1.26 mL, 16.3 mmol, 1.25 eq.) was added dropwise under continued stirring and the solution was left stirring for 1 h at 0 °C, was warmed to rt and stirred for 3 h. 10 mL 10% $NH_4Cl$ solution was added and the aq. layer was extracted three times with 100 mL DCM. The combined organic phases were washed with 50 mL brine, dried over $MgSO_4$, filtered and the solvent was removed. Flash column chromatography (0 to 3% MeOH in DCM for 9 CV) afforded compound **3** (2.5 g, 12.4 mmol, 95%) as a pale-yellow oil. **$^1$H NMR** (400 MHz, $CDCl_3$): δ 4.22 (t, $J$ = 6.4 Hz, 2H), 3.29 (t, $J$ = 6.7 Hz, 2H), 3.00 (s, 3H), 1.82 – 1.73 (m, 2H), 1.68 – 1.57 (m, 2H), 1.54 – 1.44 (m, 2H). **$^{13}$C NMR** (101 MHz, $CDCl_3$): δ 69.70, 51.20, 37.44, 28.77, 28.37, 22.84. **HRMS** (m/z): $[M + H]^+$ calcd. for $C_6H_{16}N_3O_3S^+$, 208.0750; found, 208.0753.

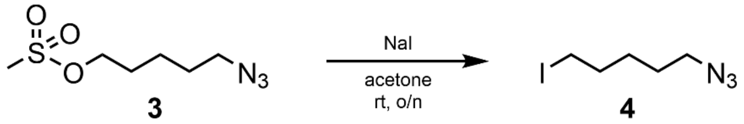

**5-azido-1-iodopentane** (**4**): **3** (2.5 g,12.4 mmol, 1 eq.) and NaI (18.6 g, 124 mmol, 10 eq.) were mixed in 18 mL acetone in a round-bottom flask. The suspension was stirred at rt overnight. Residual solvent was removed *in vacuo* and the remaining solid was dissolved in 100 mL DCM and $H_2O$ each. The aq. layer was extracted twice with 100 mL DCM. The organic layers were combined, washed with 100 mL sat. $Na_2S_2O_3$ and brine, dried over $MgSO_4$, filtered and the solvent was removed. Flash column chromatography (5 to 30% EtOAc in n-hexane in 6 CV) afforded compound **4** (2.6 g, 11.3 mmol, 91%) as a colorless liquid. **$^1$H NMR** (400 MHz, $CDCl_3$): δ 3.29 (t, J = 6.8 Hz, 2H), 3.19 (t, J = 6.9 Hz, 2H), 1.85 (p, J = 7.0 Hz, 2H), 1.68 – 1.43 (m, 4H). **$^{13}$C NMR** (101 MHz, $CDCl_3$): δ 51.31, 33.02, 27.97, 27.79, 6.51.

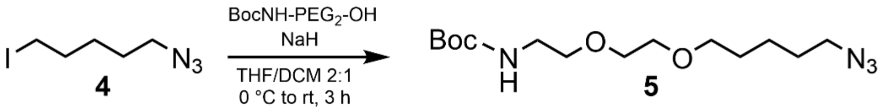

***tert*-Butyl N-[2-[2-(5-azidopentyloxy)ethoxy]ethyl]carbamate (5):** A heat-dried reaction tube was charged with 12 mL of a 2/1 ratio (v/v) of dry tetrahydrofuran (THF) and N,N-Dimethylformamid (DMF) under Schlenk conditions. *tert*-Butyl(2-(2-hydroxyethoxy)ethyl)carbamate (BocNH-$PEG_2$-OH, 1.2 g, 5.86 mmol, 1 eq.) was added and dissolved under vigorous stirring. The mixture was cooled to 0 °C and NaH (260 mg, 60% immersion on mineral oil, 6.5 mmol, 1.1 eq.) was added portion-wise. The evolving gas was released carefully and the mixture was left stirring at 0 °C for 30 min under an inert gas atmosphere. **4** (1.85 g, 8.2 mmol, 1.4 eq.) was added directly into the suspension at 0 °C, warmed to rt and left stirring for 3 h. 10 mL 10% $NH_4Cl$ and EtOAc were added and the aq. layer was extracted three times with 100 mL EtOAc. The org. layers were combined and washed with brine once and aq. 10% LiCl trice, dried over $MgSO_4$, filtered and the solvent was removed. Flash column chromatography ($SiO_2$. 20 to 50% EtOAc in n-hexane in 8 CV) afforded compound **5** (650 mg, 2.1 mmol, 42%) as a colorless oil.

**$^1$H NMR** (400 MHz, $CDCl_3$): δ 5.01 (s, 1H), 3.58 – 3.53 (m, 6H), 3.45 (t, J = 6.5 Hz, 2H), 3.34 – 3.22 (m, 4H), 1.67 – 1.55 (m, 4H), 1.43 (s, 9H), 1.46 – 1.37 (m, 2H). **$^{13}$C NMR** (101 MHz, $CDCl_3$): δ 156.10, 79.26, 71.20, 70.33, 70.16, 51.45, 40.40, 29.21, 28.77, 28.51 (3C), 21.17, 14.30. **HRMS** (m/z): $[M + Na]^+$ calcd. for $C_{14}H_{28}N_4O_4Na^+$, 339.2003; found, 339.2004.

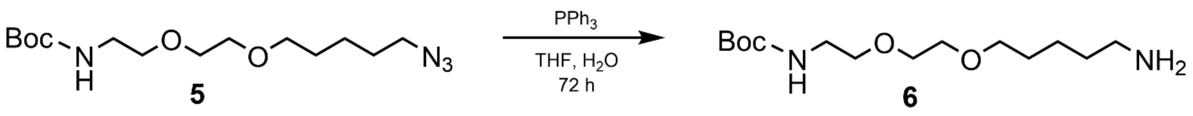

***tert*-Butyl N-[2-[2-(5-aminopentyloxy)ethoxy]ethyl]carbamate (6): 5** (550 mg, 1.56 mmol, 1 eq.) was dissolved in 2 dry THF in a dry round-bottom flask under an argon atmosphere. $PPh_3$ (550 mg, 2.3 mmol, 1.5 eq.) was added and the mixture was left stirring for 24 h air-excluded. Afterwards, $H_2O$ (440 µL, 23 mmol, 15 eq.) were added and the mixture was left stirring for 48 h. The solution was concentrated *in vacuo* and flash column chromatography ($SiO_2$. 10% MeOH, 0.5% $NEt_3$ in DCM for 12 CV) afforded **6** (500 mg, 0.79 mmol, 50%) as a colorless liquid. **$^1$H NMR** (400 MHz, MeOD): δ 3.59 (q, *J* = 1.5 Hz, 4H), 3.50 (td, *J* = 6.6, 6.1, 3.0 Hz, 4H), 3.22 (t, *J* = 5.6 Hz, 2H), 2.67 (t, *J* = 7.1 Hz, 2H), 1.67 – 1.57 (m, 2H), 1.55 – 1.47 (m, 4H), 1.44 (s, 9H). Spectrum 1**.** **$^{13}$C NMR** (101 MHz, MeOD): δ 157.60, 79.20, 71.39, 70.35, 70.18, 41.53, 40.39, 32.51, 29.63, 27.91 (3C), 23.64. **HRMS** (m/z): $[M + H]^+$ calcd. for $C_{14}H_{31}N_2O_4^+$, 291.2278; found, 291.2275.

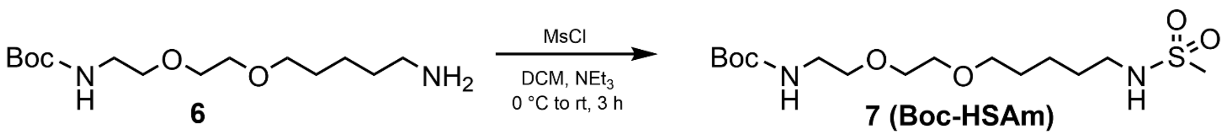

***tert*-Butyl (2-(2-((5-(methylsulfonamino)pentyl)-oxy)ethoxy)ethylcarbamate** (**7, Boc-HSAm**)*.* **6** (100 mg, 0.34 mmol, 1 eq.) was dissolved in 2.5 mL dry DCM in a dry round-bottom flask. $NEt_3$ (70 µl, 0.51mmol, 1.5 eq.) was added and the mixture was cooled to 0 °C and stirred vigorously under an argon atmosphere. Methanesulfonyl chloride (32 µl, 0.42 mmol, 1.25 eq.) was added slowly at 0 °C. The mixture was stirred for 1 h at 0 °C, warmed to rt and left stirring overnight at rt. Afterwards, 5 mL DCM and 2.5 mL 1 N HCl was added and the aq. phase was extracted twice with 10 mL DCM. The combined organic layers were washed with 10 mL brine, dried over $MgSO_4$, filtered and the solvent was removed. Flash column chromatography ($SiO_2$. 20 to 80% EtOAc in n-hexane in 8 CV) afforded **Boc-HSAm** (**7,** 100 mg, 0.28 mmol, 64%) as a pale-yellow oil. **$^1$H NMR** (400 MHz, $CDCl_3$): δ 3.62 – 3.48 (m, 6H), 3.44 (t, *J* = 6.3 Hz, 2H), 3.29 (q, *J* = 6.6, 5.7 Hz, 2H), 3.10 (q, *J* = 6.8 Hz, 2H), 2.92 (s, 3H), 1.65 – 1.52 (m, 4H), 1,46 – 1.35 (m, 2H), 1.44 (s, 9H), Spectrum 2. **$^{13}$C NMR** (101 MHz, $CDCl_3$): δ 156.06, 79.25, 77.26, 70.98, 70.22, 70.06, 43.15, 40.28, 29.83, 28.91, 28.44 (3C), 23.19, 21.3. **HRMS** (m/z): $[M + Na]^+$ calcd. for $C_{16}H_{32}N_2O_6SNa^+$, 369.2051; found, 369.2051.

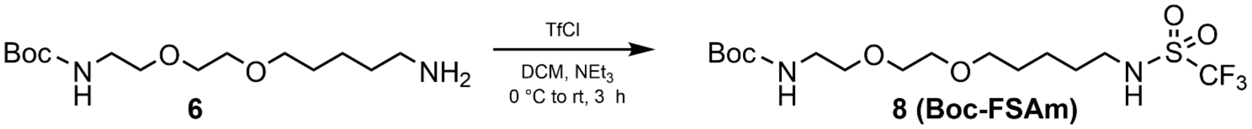

***tert*-Butyl (2-(2-((5-(trifluoromethylsulfonamino)-pentyl)oxy)ethoxy)ethylcarbamate** (**8**) was prepared analogously to **7** from **6** (100 mg, 0.34 mmol, 1 eq.) and trifluormethanesulfonyl chloride. Flash column chromatography ($SiO_2$. 40 to 80% EtOAc in n-hexane in 8 CV) afforded **Boc-FSAm** (**8,** 150 mg, 0.34 mmol, 99%) as a pale-yellow oil. **$^1$H NMR** (400 MHz, $CDCl_3$): δ 3.65 – 3.51 (m, 6H), 3.49 (t, J = 5.8 Hz, 2H), 3.30 (q, J = 6.3 Hz, 4H), 1.69 – 1.59 (m, 4H), 1,50 (td, J = 9.6, 7.0, 3.5 Hz, 2H), 1.44 (s, 9H), Spectrum 3. **$^{13}$C NMR** (101 MHz, $CDCl_3$): δ 156.34, 124.50 (q, $CF_3$), 77.36, 77.48, 70.99, 70.27 (3C), 60.58, 44.30, 29.83, 28.50 (4C), 23.2. **HRMS** (m/z): [M + Na]$^+$ calcd. for $C_{16}H_{29}N_2O_6SF_3Na^+$, 445.1591; found, 445.1587.

#### II Hy4 / Hy5-xHTL

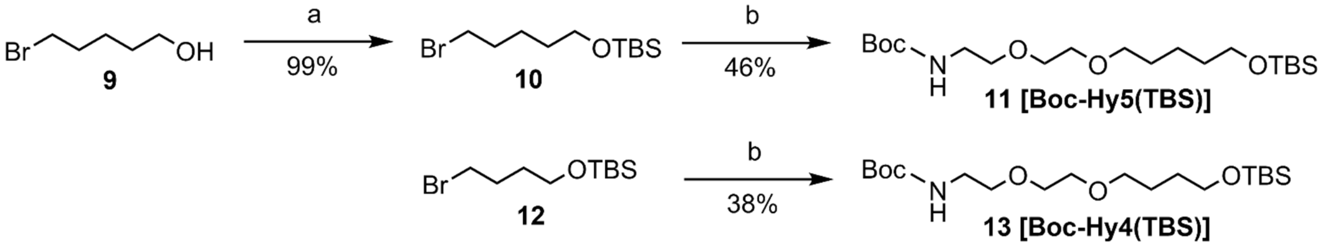

**Method Figure 2 Synthetic route to xHTL precursor Boc-Hy5(TBS) (11) and Boc-Hy4(TBS) (13).**
**a**. TBS-Cl, DCM, imidazole, 1 h, 0 °C to rt. **b.** BocNH-PEG$_2$-OH, NaH, THF/DMF, o/n, 0 °C to rt.

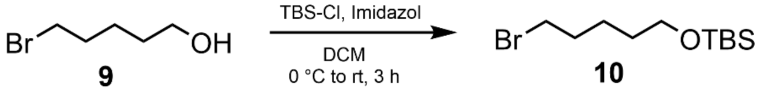

***tert*-Butyl (5-bromopentoxy)-dimethylsilane (10)**. 1-Bromopentan-5-ol (**9**, 1.5 g, 9 mmol, 1 eq.) was dissolved in 20 mL dry DCM and imidazole (930 mg, 13.5 mmol, 1.5 eq.) was added. The mixture was cooled to 0 °C under vigorous stirring and *tert*-Butyldimethylsilyl chloride (TBS-Cl, 1.5 g, 10 mmol, 1.1 eq.) was added portion-wise. After 1 h stirring at rt 10 mL 10% $NH_4Cl$, the aq. layer was extracted with 50 mL DCM. The org. phases were combined and washed with 100 mL brine, dried over $MgSO_4$, filtered and the solvent was removed. Flash column purification ($SiO_2$. 0 to 30% EtOAc/n-hexane in 8 CV) afforded **10** (2.5 g, 9 mmol, 99%) as a colorless liquid. **$^1$H NMR** (400 MHz, $CDCl_3$): δ 3.61 (t, J = 6.1 Hz, 2H), 3.41 (t, J = 6.8 Hz, 2H), 1.88 (m, 2H), 1.52 (m, 4H), 0.89 (s, 9H), 0.05 (s, 6H). **$^{13}$C NMR** (101 MHz, $CDCl_3$): δ 62.99, 33.96, 32.76, 32.05, 26.01 (3C), 24.73, - 5.15 (2C). **HRMS** (m/z): [M + H]$^+$ calcd. for $C_{11}H_{25}BrOSi^+$, 281.0931, 283.0911; found, 281.0933, 283.0913.

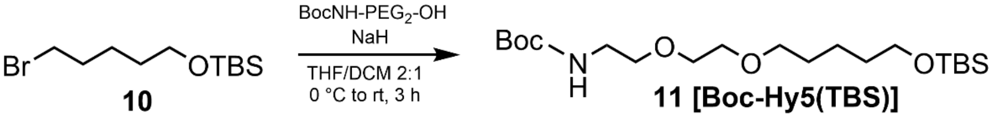

***tert*-Butyl (2,2,3,3-tetramethyl-4,8,11-trioxa-3-silapentadecan-15-yl)carbamate** (**11**, Boc-Hy5-TBS). The title compound was synthesized similarly as described for (**5**) from **10** (2 g, 7.1 mmol, 1 eq.) with the exception that the reaction required 16 h and afforded **Boc-Hy5-TBS** (1.3 g, 3.3 mmol, 46%) as a colorless liquid. **$^1$H NMR** (400 MHz, $CDCl_3$): δ 3.62 – 3.49 (m, 8H), 3.44 (t, *J* = 6.8 Hz, 2H), 3.29 (q, *J* = 5.3 Hz, 2H), 1.69 – 1.55 (m, 2H), 1.55 – 1.46 (m, 2H), 1.42 (s, 9H), 1.39 – 1.28 (m, 2H), 0.87 (s, 9H), 0.02 (s, 6H), Spectrum 4. **$^{13}$C NMR** (101 MHz, $CDCl_3$): δ 156.11, 71.57, 50.39, 70.32, 70. 12, 40.46, 37.75, 29.49, 28.58 (3C), 26.08 (3C), 22.45, 18.46, -5.17 (2C). **HRMS** (m/z): $[M + H]^+$ calcd. for $C_{20}H_{43}NO_5Si^+$, 406.2983; found, 406.2983.

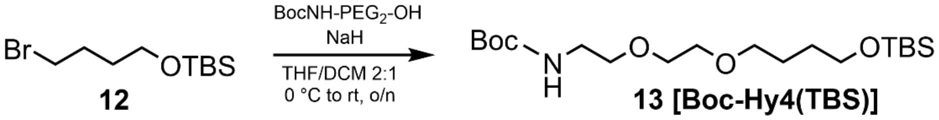

***tert*-Butyl (2,2,3,3-tetramethyl-4,8,11-trioxa-3-silatetradecan-14-yl)carbamate** (**13**, Boc-Hy4-TBS). The title compound was synthesized similarly as described for (**5**) from commercially available *tert*-butyl (3-bromobutoxy)dimethylsilane (**12**, 2 mL, 7.2 mmol, 1 eq.) with the exception that the reaction required 16 h and afforded **Boc-Hy4-TBS** (820 mg, 2.7 mmol, 38%). **$^1$H NMR** (400 MHz, $CDCl_3$): δ 3.66 – 3.51 (m, 8H), 3.48 (t, *J* = 6.5 Hz, 2H), 3.31 (s, 2H), 1.69 – 1.61 (m, 2H), 1.60 – 1.53 (m, 2H), 1.44 (s, 9H), 0.89 (s, 9H), 0.04 (s, 6H), Spectrum 5. **HRMS** (m/z): $[M + Na]^+$ (ΔTBST) calcd. for $C_{13}H_{27}NO_5Na$, 300,1718; found 300,1718.

#### III Deprotection

For Boc and TBS protecting groups removal, compound **7**, **8**, **11** or **13** were dissolved in 3 mL/mmol of a 1/1 (v/v) mixture of trifluoroacetic acid (TFA) in dry DCM. It was stirred at rt for at least 3 h and the solvent was removed under a stream of argon gas. Residual TFA was removed by co-evaporation with DCM. Resulting xHTL were obtained in quantitative yields and used for fluorophore coupling without further purification.

**N-(5-(2-(2-aminoethoxy)ethoxy)pentyl)methanesulfonamide** (**14, HSAm**). **$^1$H NMR** (400 MHz, MeOD): δ 3.65 – 3.50 (m, 6H), 3.43 (t, J = 6.4 Hz, 2H), 3.04 (t, J = 5.0 Hz, 2H), 2.97 (t, J = 6.8 Hz, 2H), 1.58 – 1.43 (m, 4H), 1.41 – 1.31 (m, 2H). **$^{13}$C NMR** (101 MHz, MeOD): δ 71.53, 70.68, 70.50, 67.22, 43.26, 40.04, 38.91, 30.26, 29.47, 23.56. **HRMS** (m/z): $[M + H]^+$ calcd. for $C_{10}H_{24}N_2O_4S$, 269.1530 found, 269.1528.

**N-(5-(2-(2-aminoethoxy)ethoxy)pentyl)trifluoromethanesulfonamide** (**15, FSAm**). **$^1$H NMR** (400 MHz, $CDCl_3$): δ 3.70 – 3.58 (m, 6H), 3.51 (t, J = 6.2 Hz, 2H), 3.21 (t, J = 5.2, 2H), 3.12 (t, 6.4 Hz, 2H), 1.62 – 1.54 (m, 4H), 1.48 – 1.42 (m, 2H). **$^{13}$C NMR** (101 MHz, $CDCl_3$): δ 121.50 (q, $CF_3$),70.69, 69.92, 69.77, 68.47, 43.48, 39.28, 29.66, 28.61, 22.52. **$^{19}$F NMR** (377 MHz, $CDCl_3$) δ 79.52. **HRMS** (m/z): $[M + H]^+$ calcd. for $C_{10}H_{21}F_3N_2O_4S$, 323.1243 found, 323.1247.

**5-(2-(2-aminoethoxy)ethoxy)pentan-1-ol (16, Hy5).** **$^1$H NMR** (400 MHz, MeOD): δ 4.28 (t, J = 6.6 Hz, 2H), 3.70 – 3.61 (m, 2H), 3.57 (dd, J = 5.8, 2.9 Hz, 2H), 3.50 (dd, J = 5.7, 3.0 Hz, 2H), 3.40 (t, J = 6.6 Hz, 2H), 3.09 (s, 2H), 1.69 (dt, J = 14.9, 6.8 Hz, 2H), 1.54 (q, J = 7.1 Hz, 2H), 1.40 – 1.28 (m, 2H). **$^{13}$C NMR** (101 MHz, MeOD): δ 71.01, 70.31, 69.90, 68.16, 39.70, 28.89, 27.85, 22.18. **HRMS** (m/z): $[M + H]^+$ calcd. for $C_9H_{21}NO_3$, 192.1594 found, 192.1593.

**5-(2-(2-aminoethoxy)ethoxy)butan-1-ol (17, Hy4).** **$^1$H NMR** (400 MHz, MeOD): δ 3.72 – 3.61 (m, 6H), 3.57 (t, J = 6.2 Hz, 2H), 3.53 (t, J = 6.2 Hz, 2H), 3.15 – 3.11 (m, 2H), 1.72 – 1.54 (m, 4H). **HRMS** (m/z): $[M + H]^+$ calcd. for $C_8H_{19}NO_3$, 178.1438 found, 178.1439.

#### IV Fluorophore coupling

For fluorophore coupling, 6-carboxy rhodamine (1-5 mg, 1 eq.) was dissolved in ~100 µL/µmol dry DMSO-$d_6$. The mixture was added on top of N,N,N′,N′-Tetramethyl-O-(N-succinimidyl)uroniumtetrafluorborat (TSTU, 1.2 eq.) and diisopropylethylamine (DIPEA, 10 eq.) were spiked into the solution. It was left stirring for at least 10 min, the xHTL **14** - **17** were dissolved in the same amount of dry DMSO-$d_6$ with 10 eq. DIPEA and both solutions were mixed and left stirring for 2 h at 40 °C. The desired products were obtained by RP-HPLC $H_2O$/MeCN linear gradient from 20 to 90% containing 0.1% TFA.

**TMR-HSAm** – **$^1$H NMR** (400 MHz, $CD_3CN$): δ 8.11 (d, J = 8.1 Hz, 1H), 8.05 (dd, J = 8.1, 1.5 Hz, 1H), 7.58 (d, J = 1.5 Hz, 1H), 7.32 (t, J = 5.6 Hz, 1H), 6.78 (d, J = 8.8 Hz, 2H), 6.65 – 6.60 (m, 4H), 3.57 – 3.49 (m, 4H), 3.48 – 3.43 (m, 4H), 3.32 (t, J = 6.4 Hz, 2H), 3.06 (s, 12H), 2.95 (q, J = 6.7 Hz, 2H), 2.82 (s, 3H), 1.44 (h, J = 6.8, 6.4 Hz, 4H), 1.33 – 1.22 (m, 2H). Spectrum 6.

**TMR-FSAm** – **$^1$H NMR** (400 MHz, $CD_3CN$): δ 8.28 (d, J = 8.1 Hz, 1H), 8.08 (dd, J = 8.2, 1.7 Hz, 1H), 7.69 (d, J = 1.7 Hz, 1H), 7.40 (t, J = 5.6 Hz, 1H), 7.04 (d, J = 9.4 Hz, 2H), 6.86 (dd, J = 9.4, 2.5 Hz, 2H), 6.79 (d, J = 2.5 Hz, 2H), 3.62 – 3.44 (m, 8H), 3.35 (t, J = 6.4 Hz, 2H), 3.20 (s, 12H), 1.57 – 1.40 (m, 4H), 1.36 – 1.24 (m, 4H). Spectrum 7.

**TMR-Hy5** – **$^1$H NMR** (400 MHz, $CD_3CN$): δ 8.24 (d, *J* = 8.2 Hz, 1H), 8.09 (dd, *J* = 8.1, 1.7 Hz, 1H), 7.67 (d, *J* = 1.7 Hz, 1H), 7.36 (d, *J* = 5.7 Hz, 1H), 6.97 (d, *J* = 9.3 Hz, 2H), 6.81 (dd, *J* = 9.3, 2.5 Hz, 2H), 6.75 (d, *J* = 2.5 Hz, 2H), 3.60 – 3.44 (m, 7H), 3.42 (t, *J* = 6.4 Hz, 2H), 3.35 (t, *J* = 6.5 Hz, 2H), 3.17 (s, 12H), 1.43 (tt, *J* = 14.2, 6.4 Hz, 4H), 1.35 – 1.18 (m, 4H). Spectrum 8.

TMR-Hy4 – **$^1$H NMR** (400 MHz, $CD_3CN$): δ 8.29 (d, J = 8.2 Hz, 1H), 8.11 (dd, J = 8.2, 1.8 Hz, 1H), 7.71 (d, J = 1.8 Hz, 1H), 7.55 (t, J = 5.7 Hz, 1H), 7.05 (d, J = 9.4 Hz, 2H), 6.88 (dd, J = 9.4, 2.5 Hz, 2H), 6.81 (d, J = 2.4 Hz, 2H), 3.63 – 3.45 (m, 8H), 3.34 (t, J = 6.7 Hz, 2H), 3.21 (s, 12H), 1.47 – 1.38 (m, 2H), 1.30 – 1.20 (m, 2H). Spectrum 9.

**Method Table 2.** HRMS data of fluorophore derivatized xHTLs.

| xHTL | Dye | Chem. Form. | $[M]_{calcd.}$ | $[M]_{found}$ | xHTL | Dye | Chem. Form. | $[M]_{calcd.}$ | $[M]_{found}$ |
|---|---|---|---|---|---|---|---|---|---|
| HSAm | $JF_{525}$ | $C_{37}H_{41}N_4O_8F_4S$ | 777.2575 | 777.2575 | Hy5 | $JF_{525}$ | $C_{36}H_{38}N_3O_7F_4$ | 700.2640 | 700.2640 |
| | TMR | $C_{35}H_{45}N_4O_8S$ | 681.2953 | 681.2952 | | TMR | $C_{34}H_{42}N_3O_7$ | 604.3017 | 604.3016 |
| | MaP555 | $C_{37}H_{51}N_{46}O_9S_2$ | 787.3154 | 787.3152 | | MaP555 | $C_{36}H_{47}N_5O_8S$ | 710.2118 | 710.2120 |
| | $JF_{585}$ | $C_{40}H_{47}N_4O_7F_4S$ | 803.3096 | 803.3097 | | CRh | $C_{37}H_{47}N_3O_6$ | 630.3538 | 630.3538 |
| | CRh | $C_{38}H_{51}N_4O_7S$ | 707.3473 | 707.3473 | | MaP618 | $C_{39}H_{53}N_5O_7S$ | 736.3738 | 736.3735 |
| | MaP618 | $C_{40}H_{51}N_6O_8S_2$ | 813.3674 | 813.3652 | | SiR | $C_{36}H_{48}N_3O_6Si$ | 646.3307 | 646.3306 |
| | $JF_{635}$ | $C_{39}H_{49}N_4O_7F_2SSi$ | 783.3054 | 783.3048 | | SiR700 | $C_{38}H_{48}N_3O_6Si$ | 670.3307 | 670.3296 |
| | $JF_{646}$ | $C_{39}H_{51}N_4O_7SSi$ | 747.3242 | 747.3239 | | $JF_{525}$ | $C_{36}H_{38}N_3O_7F_4$ | 700.2640 | 700.2640 |
| | SiR | $C_{37}H_{51}N_4O_7SSi$ | 723.3242 | 723.3240 | Hy4 | TMR | $C_{33}H_{40}N_3O_7Na^+$ | 444.0431 | 444.0428 |
| | SiR700 | $C_{39}H_{51}N_4O_7SSi$ | 747.3242 | 747.3247 | | $JF_{585}$ | $C_{38}H_{42}N_3O_6F_4$ | 712.3004 | 712.3005 |
| FSAm | $JF_{525}$ | $C_{37}H_{38}N_4O_8F_7S$ | 831.2290 | 831.2293 | | CRh | $C_{36}H_{46}N_3O_6$ | 616.3381 | 616.3378 |
| | TMR | $C_{35}H_{42}N_4O_8F_3S$ | 735.2665 | 735.2670 | | MaP618 | $C_{38}H_{51}N_5O_7S$ | 722.3582 | 722.3579 |
| | MaP555 | $C_{37}H_{48}N_6O_9F_3S_2$ | 841.2871 | 841.2870 | | $JF_{635}$ | $C_{37}H_{44}N_3O_6F_2Si$ | 692.2962 | 692.2945 |
| | $JF_{585}$ | $C_{40}H_{44}N_4O_7F_7S$ | 857.2813 | 857.2813 | | $JF_{646}$ | $C_{37}H_{46}N_3O_6Si$ | 659.3158 | 659.3150 |
| | CRh | $C_{38}H_{48}N_4O_7SF_3$ | 761.3190 | 761.3193 | | SiR | $C_{35}H_{46}N_3O_6Si$ | 632.3150 | 632.3153 |
| | SiR | $C_{37}H_{48}N_4O_7SF_3Si$ | 777.2966 | 777.2960 | | SiR700 | $C_{37}H_{46}N_3O_6Si$ | 656.3150 | 656.3148 |

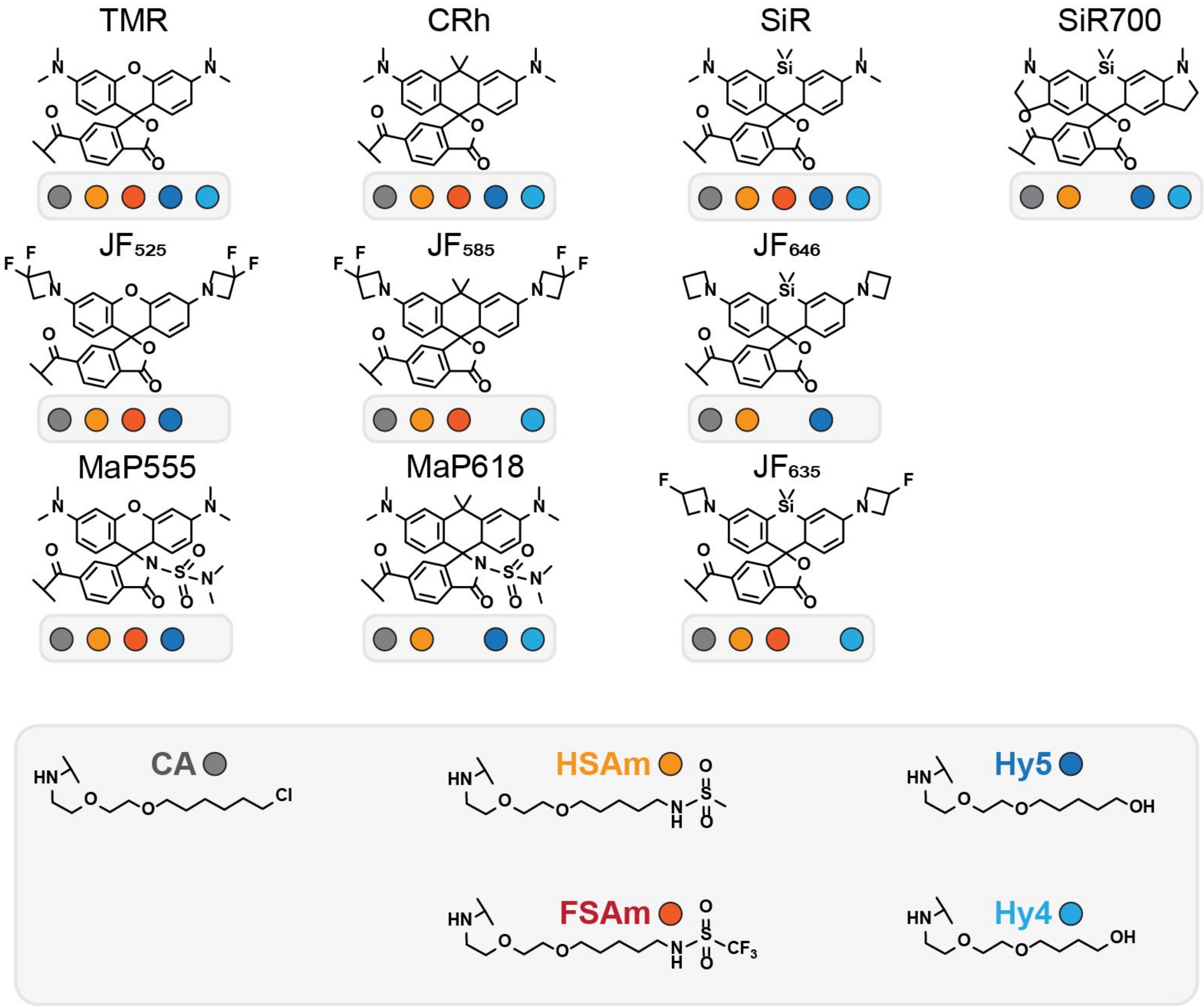

**Method Figure 3. Chemical structures of fluorophore-derivatives of different HaloTag ligands (x)HTLs.** Coloured dots indicate which fluorophore derivatives were characterized in this study.

#### Molecular biology & Biochemistry

##### Plasmids

HaloTag7 and its dead variant (HaloTag7-D106A = dHaloTag7)[14] proteins were produced in *Escherichia coli* using a pET51b(+) expression vector (Novagen) carrying a N-terminal $His_{x10}$-tag and a tobacco etch virus (TEV)[15] protease cleavage site, as previously reported.[1] Mammalian protein expression was performed using the pcDNA5/FRT/TO expression vector (ThermoFisher Scientific). HaloTag variants were fused to different sub-cellular markers (H2B, CEP41, TOM20, COX8, CalR/KDEL, LAMP1, Ig-κ-/PDGFR or LifeAct) whose origins are reported in **Method Table 8.**[1,16,17] Proteins were co-expressed by translational fusion using the self-cleaving peptide sequence T2A.[18,19] Molecular cloning was performed by Gibson assembly.[20] dHaloTag variant was obtained by introducing a point-mutation in HaloTag7 with the Q5®-site-directed mutagenesis kit (NEB) according to the manufacturer protocol. Plasmid sequences were verified by Sanger sequencing (Eurofins) and plasmid DNA was stored at -20 °C.

##### Protein production and purification

pET51b(+) plasmids were transformed in *E. coli* strain BL21(DE3)-pLysS (Novagen) and cultured at 37 °C in $LB^{Amp}$ (**Method Table 3**) until an optical density at 600 nm ($OD_{600}$) of 0.8 was reached. Then, the temperature was reduced to 18 °C and transgene expression was induced with 0.5 mM isopropyl β-thiogalacopyranoside (IPTG). After overnight expression, cells were harvested by centrifugation (4 000 g, 15 min, 4 °C), resuspended in ice-cold extraction buffer (**Method Table 3**) including 1 mM phenylmethylsulphonyl fluoride (PMFS) and 0.25 mg/mL lysozyme. Cells were lyzed by sonication (SONOPLUS, 7 min, 50% on/off cycles, 70% amplitude) on wet ice. The lysate was cleared from the cell debris by centrifugation (20 min, 50 000 g, 4 °C). Proteins were purified via immobilized metal ion affinity chromatography (IMAC) using a HisTrap FF crude column (Cytiva, Marlborough, MA) and an ÄktaPure FPLC instrument (Cytiva) with IMAC wash and elution buffers (**Method Table 3**). Buffer was exchanged to activity buffer on a HiTrap® 26/10 Desalting Column (Cytiva) (**Method Table 3**) using the ÄktaPure FPLC.

**Method Table 3.** Buffer and media composition.

| Buffer | Composition |
|---|---|
| $LB^{Amp}$ | 5 g/L yeast extract, 10 g/L peptone, 0.1 g/L ampicillin |
| Activity buffer | 50 mM HEPES, 50 mM NaCl, pH 7.3 |
| Extraction buffer | 50 mM $KH_2PO_4$, 300 mM NaCl, 5 mM imidazole, pH 8.0 |
| IMAC wash buffer | 50 mM $KH_2PO_4$, 300 mM NaCl, 10 mM imidazole, pH 7.5 |
| IMAC elution buffer | 50 mM $KH_2PO_4$, 300 mM NaCl, 500 mM imidazole, pH 7.5 |
| TEV-cleavage buffer | 25 mM $Na_2HPO4$, 200 mM NaCl, pH 8.0 |
| Laemmli Buffer | 31.5 mM Tris-HCl, pH 6.8, 10% glycerol, 1% SDS, 0.005% Bromophenol Blue |

Buffers/media were prepared in Mili-Q® water.

The protein solution was concentrated with Ultra 15 mL Centrifugal Filters (Amicon®, MWKO: 10 kDa). Proteins were quantified by UV-absorption at 280 nm (NanoDrop™ 2000c, ThermoFisher) using the molar extinction coefficient ε of (d)HaloTag7 (60 055 $M^{-1}$ $cm^{-1}$) and concentration was adjusted to 500 µM with activity buffer. Finally, proteins were aliquoted, flash frozen in liquid nitrogen and stored at -80 °C. Furthermore, the correct molecular weight and purity of proteins were assessed by sodium dodecyl sulfate–polyacrylamide gel electrophoresis (SDS–PAGE). ~50 µg of isolated protein was denatured in Laemmli sample buffer (BioRad with 10 mM dithiothreitol (DTT)) at 95 °C for 10 min and loaded onto a stain-free gel (mini-PROTEAN® TGX™ Stain-Free™, 4 - 20%, Bio-Rad) besides a *PrecisionPlus Protein™ All Blue* (Bio-Rad) pre-stained marker. Proteins were revealed by 2 min of UV light exposure and imaged by applying a UV-trans illumination from a ChemiDoc XRS+ imager (Bio-Rad, **Method Table 4**). Additionally, the correct mass of purified proteins was confirmed by electrospray-ionization mass-spectrometry (ESI-MS) performed by the MS-facility (MPImR, Heidelberg) on a maXis II ETD-instrument. Protein amino acid sequences are listed at the protein sequences section.

##### Covalent labeling of HaloTag7 protein and in-gel fluorescence scan

Purified HaloTag7 protein (5 µM) was incubated in presence of 4x excess of the (x)HTLs (20 µM). Reaction was carried out in activity buffer for 30 min at 37 °C. 5 µg protein were separated by SDS-PAGE as previously explained.[1] Covalent labeling was revealed by in-gel fluorescence imaging performed on a ChemiDoc XRS+ imager (Bio-Rad) using either the green or red epi illumination (**Method Table 4**). These gels were further Coomassie stained (Bio-Rad) and imaged by white light converted UV trans illumination (**Method Table 4**) using the same imager.

**Method Table 4.** Imaging settings in-gel fluorescence

| Channel | $\lambda_{ext}$ [nm] | $\lambda_{em}$ [nm] | Detection |
|---|---|---|---|
| Epi-red | 520 - 545 | 605 ± 50 | TMR |
| Epi-green | 625 - 650 | 695 ± 5 | SiR |
| Epi-white | / | 590 ± 110 | Coomassie |
| Trans-UV | 302 | 590 ± 110 | Unstained proteins |

##### Protein crystallization and X-ray diffraction data collection

For protein crystallization, HaloTag7 and dHaloTag7 (D106A variant) proteins were produced and purified as previously described (carried out by A. Bergner)[1,17]. In short, after IMAC purification the buffer was exchanged to TEV-cleavage buffer (**Method Table 3**). TEV protease[15] (weight ratio 30:1) were added to protein sample and cleavage was performed at 30 °C overnight. The solution was then filtered (0.22 µm) and the cleaved protein was harvested by reverse IMAC purification on a HisTrap FF crude column (Cytiva) on an ÄktaPure FPLC (Cytiva) as previously explained but collecting the flow through. The protein was then concentrated and further purified by size exclusion chromatography on a HiLoad 26/600 Superdex 75 pg column (Cytiva) exchanging the buffer to activity buffer. The protein was concentrated again to a final concentration of 5 µM. 10 µM TMR-xHTL was added to the protein for at least 4 h at room temperature (rt). Afterwards, the protein was concentrated to ~250 µL reaching a final concentration between 10 – 15 mg/mL of HaloTag protein (quantified using the absorbance at 280 nm and the extinction coefficient of the protein corrected for TMR absorbance at 280 nm, ${}^{TMR}CF_{280nm}$ = 0.16, 13 920 $M^{-1}$ $cm^{-1}$).

Crystallization was performed at 20 °C using the vapor-diffusion method. The HaloTag7 or dHaloTag7 in complex with tetramethylrhodamine (TMR) ligands, were concentrated to 10 – 15 mg/mL in 50 mM HEPES pH 7.3, 50 mM sodium chloride. Crystals of HaloTag7 with HSAm-TMR ligands and dHaloTag7 (D106A variant) with Hy5-TMR ligands were grown by mixing equal volumes of protein solution and a reservoir solution containing 0.1 M MES pH 6.0, 1.0 M lithium chloride and 19-20% (m/v) PEG 6000. Crystals of HaloTag7 with FSAm-TMR ligand were obtained by mixing equal volumes of protein solution and precipitant solution composed of 0.1 M MES pH 6.0, 0.2 M calcium acetate, 21% (m/v) PEG 8000. In both cases, the crystals were briefly washed in cryoprotectant solution consisting of the reservoir solution with glycerol added to a final concentration of 20% (v/v), prior to flash-cooling in liquid nitrogen.

Single crystal X-ray diffraction data were collected at 100 K on the X10SA beamline at the SLS (PSI, Villigen, Switzerland). All data were processed with XDS[21]. The structures of HaloTag7 in complex with TMR ligands were determined by molecular replacement (MR) using Phaser[22] and HaloTag7-TMR coordinates (PDB-ID: 6Y7A) as a search model. Geometrical restraints for TMR ligands were generated using the Grade server[23]. The final models were optimized in iterative cycles of manual rebuilding using Coot[24] and refinement using Refmac5[25] and phenix.refine[26]. Data collection and refinement statistics are summarized in **Table S2**, model quality was validated with MolProbity[27] as implemented in PHENIX. Atomic coordinates and structure factors have been deposited in the Protein Data Bank under accession codes: 7ZJ0 (HaloTag7 HSAm-TMR), 7ZIY (HaloTag7 FSAm-TMR) and 7ZIZ (dHaloTag7 Hy5-TMR).

#### Fluorophore concentrations

The concentrations of fluorophore-ligands were determined by UV-Vis absorption at their maximum absorption wavelength (**Method Table 1**) using the Lambert-Beer's law (**eq. 1**):

$$\mathrm{Abs} = \mathrm{c} \cdot \mathrm{d} \cdot \varepsilon \qquad (\mathbf{1})$$

Abs: absorption at $\lambda_{max}$ [A.U.], c: concentration [M], d: pathway length [cm], ε extinction coefficient [1/(M cm)]

Absorbance was measured with a Nanodrop 2000c™ spectrophotometer (ThermoFisher) using 2 µL of solution (d = 0.1 cm) or 500 µL in a polystyrene cuvette (Sarstedt, 10 x 4 x 45 mm, d = 1 cm). Samples were prepared in PBS pH 7.4 (Gibco), 0.1% w/v SDS in activity buffer or 0.1% v/v TFA in ethanol, depending of the fluorophore properties and according to literature precedents (**Method Table 1**).[6,28] In-house characterized extinction coefficients ε (for detailed description see below) were used to calculate the concentration of neo-synthesized fluorophore ligands (see **Method Table 1, Method Figure 3**). Fluorescent molecules were prepared as stock solutions in dry DMSO (> 1 mM) and diluted such that the DMSO concentration never exceed 1% (v/v) in experiments.

#### Water-Dioxane titration

Solutions of SiR-(x)HTLs (5 µM, 100 µL) were prepared in 20/80 to 90/10 (v/v) water-dioxane mixtures (dry, Acros) in transparent flat-bottom, chimney well polypropylene 96-well plates (Greiner Bio-One) in technical triplicates. Absorbance spectra were recorded between 400 and 750 nm in a microplate reader (Spark20M, Tecan) with 2 nm step size. The maximal absorbance of SiR at 646 nm ($\lambda_{max}$) was blank corrected in each condition. The data was normalized to the max. absorbance measured (SiR-HSAm) and presented as the ratio of absorbance variations between 0 and 1. The data from three technical replicates was averaged and plotted against the dielectric constants $\varepsilon_R$ of the water-dioxane mixtures.[29] A sigmoidal function (**eq. 2**) was fitted to the data yielding $D_{50}$-values as the point of inflection. Mean values and standard deviations from three independent experiments are presented.

$$Abs = \frac{1}{(1+10^{(D50-\varepsilon_R)})} \qquad (\mathbf{2})$$

Abs: Norm. absorbance, $\varepsilon_R$: dielectric constant, $D_{50}$: $\varepsilon_R$ at half-maximal absorbance.

#### Quantum yield

The fluorophore-xHTLs (500 nM) were incubated with or without 100 µM (d)HaloTag7 protein in activity buffer in 800 µL Glass Crimp Neck Vial (8 mm, Fisherbrand™, used for TMR, MaP555, CRh, MaP618) or in 2 mL Quarz cuvettes (used for $JF_{635}$, SiR) for 2 h at rt. Quantum yields were measured in technical triplicates by exciting the fluorophore at its maximum absorbance ($\lambda^{abs}$) (**Method Table 1**) in an absolute quantum yield spectrometer (Hamatsu, Quantaurs-QY, model C11347). Mean values and standard deviations from three individual measurements (n = 3) are given.

#### Extinction coefficients

Prior to extinction coefficient measurements the fluorophore-xHTL ligands were quantified by qNMR using a dioxane standard (5 µM). A dilution series of 0.5, 1, 1.5, 2 and 3 µM of fluorophore-xHTL was prepared in activity buffer, 0.1% SDS in activity buffer or in presence of 100 µM (d)HaloTag7 protein in a total volume of 200 µL (clear bottom non-binding 96-well plates, Greiner). Absorbance spectra were recorded on a plate reader (Spark20M, Tecan) from 400 to 700 nm. The data was baseline corrected and the maximum absorbance values were plotted against the concentration. A linear function (**eq. 3**) was fitted to the data and extinction coefficients were calculated from the slope b which was corrected for the path length using the Lambert-Beer's law (**eq 1.**). The path length in 96-well plates was obtained measuring the absorbance of SiR-CA in 0.1% SDS for which the extinction coefficient was previously reported.[10]

$$Abs = a + bx \qquad (\mathbf{3})$$

Abs: absorbance [A.U.], x: fluorophore concentration [M].

#### Fluorescence increase upon protein binding assay

The fluorophore-(x)HTLs (50 nM) were incubated in presence and absence of (d)HaloTag7 protein (100 µM) for 30 min at 37 °C in activity buffer containing 1% (w/v) BSA in a black flat bottom 384 well plate (Greiner, 20 µL). Fluorescence emission scans were recorded by exciting the fluorophore and measuring the fluorescence emission intensity over a spectral range covering the maximum emission (2 nm step, 10 nm bandwidth) on a microplate reader (Spark20M, Tecan) using the settings summarized in **Method Table 5**. For representation, the fluorescence emission was normalized to the max. intensity measured for each dye (*e.g.* SiR-HSAm). The data from three technical replicates were averaged and the fluorescence intensity increase represented by their ratio in presence and absence of the respective HaloTag protein ($F/F_0$) at the fluorescence emission maxima was obtained (**Method Table 5**). The uncertainty was calculated through propagation of uncertainty from the replicate's standard deviations.

**Method Table 5.** Spectral settings used for fluorescence turn-on assay.

| Fluorophore | $\lambda_{ext}$ [nm] | $\lambda_{em}$ [nm] | $\lambda_{em,\,max}$ [nm] |
|---|---|---|---|
| $JF_{525}$ | 500 ± 10 | 540 - 750 | 558 |
| TMR | 530 ± 10 | 570 - 750 | 580 |
| MaP555 | 530 ± 10 | 570 - 750 | 580 |
| $JF_{585}$ | 560 ± 10 | 602 - 750 | 612 |
| CRh | 580 ± 10 | 620 - 800 | 634 |
| MaP618 | 580 ± 10 | 620 - 800 | 638 |
| $JF_{635}$ | 600 ± 10 | 638 - 800 | 660 |
| SiR | 605 ± 10 | 652 - 800 | 670 |
| $JF_{656}$ | 610 ± 10 | 650 - 800 | 670 |

#### Affinity measurement by fluorescence polarisation assays

The dissociation constants $K_d$ of fluorescent xHTL (10 nM) and (d)HaloTag proteins [0 – 100 µM] were determined in activity buffer supplemented with 0.5% (w/v) Bovine Serum Albumin (BSA, Fraktion V, Roth) as previously described.[1] All measurement were performed in black flat bottom low-volume 384 well plates (Greiner, 20 µL) at 37 °C. The fluorescence polarization (FP, **eq. 4**) was measured in technical triplicates on a microplate reader (Spark20M, Tecan). Excitation and emission settings used for various dyes are listed in **RKd: ratio** of Kd values observed for xHTLs with HaloTag7 and dHaloTag7. $K_d^{dHT7}$: $K_d^{dHT7}$:

**Method Table 6** and the gain and G factor were optimized for each dye individually and kept constant over the replicates. Maximum FP values of each dye were determined using covalently labeled HaloTag7 (100 µM) labeled with the respective fluorophore-HTL substrate (20 nM).

$$\mathrm{FP} = \frac{I_{II} - I_{\perp}G}{I_{II} + I_{\perp}G} \qquad (\mathbf{4})$$

FP: fluorescence polarization, $I_{II}$: fluorescence intensity parallel to the excitation light polarisation, $I_{\perp}$: fluorescence intensity perpendicular to the excitation light polarisation, G: grating factor $I_{II}/I_{\perp}$.

Data was normalized between 0 - 1 *i.e.* the minimum FP value provided by the free fluorophore and the maximum FP value provided by the fluorophore fully bound (or corresponding covalent labeling). Technical triplicates were averaged obtaining mean and standard deviations (S.D.) and a single-site binding model (**eq. 5**) was fitted to the data to estimate $K_d$ values. Confidence intervals and standard deviations of fitted parameters were estimated with the Monte Carlo[30] method with N = 1000. $K_d$ values are represented with 95% confidence interval (CI 95%). All experiments were performed in at least three individual replicates and all data was used for the fit ($x_n \geq 3$).

$$Y = \frac{1}{1+\frac{K_d}{[HT7]}} \quad (\mathbf{5})$$

Y: Normalized fluorescence polarization, A: min. FP, B: max. FP, [HT7]: (d)HaloTag7 protein concentration [M], $K_d$: dissociation constant [M].

The ratio of $K_d$ values observed for xHTLs with HaloTag7 and dHaloTag7 ($R_{Kd}$ or $R_{Kd}^{-1}$) was calculated according to **eq. 6**. Uncertainties were estimated with the Monte Carlo[30] method (N = 1000).

$$R_{\mathrm{Kd}} = \frac{K_d^{dHaloTag7}}{K_d^{HaloTag7}} \quad (\mathbf{6})$$

$R_{Kd}$: ratio of $K_d$ values observed for xHTLs with HaloTag7 and dHaloTag7. $K_d^{dHT7}$: $K_d^{dHT7}$:

**Method Table 6.** Spectral settings used for fluorescence polarization assays.

| Fluorophore | $\lambda_{ext}$ [nm] | $\lambda_{em}$ [nm] |
|---|---|---|
| TMR | 535 ± 12.5 | 595 ± 17.5 |
| MaP555 | 535 ± 12.5 | 595 ± 17.5 |
| CRh | 600 ± 10 | 660 ± 10 |
| SiR/$JF_{635}$ | 635 ± 10 | 655 ± 10 |

#### Stopped-Flow kinetic fluorescence polarisation assay

Binding rate constants were determined by measuring fluorescence anisotropy changes over time with a BioLogic SFM-400 stopped flow instrument (BioLogic Science Instruments). 4 µM of HaloTag7 protein was mixed in a 1:1 (v/v) ratio (single-mix sequence) with 1 µM fluorophore-xHTL in activity buffer at 37 °C. Fluorescence intensity and fluorescence anisotropy were measured over 300 s in 0.1 ms intervals. Monochromator wavelengths for excitation and longpass (LP) emission filters are summarized in **Method Table 7**.

**Method Table 7.** Filter-settings used for stopped-flow kinetics fluorescence polarization experiments.

| Fluorophore | $\lambda_{ext}$ [nm] | Emission filter LP [nm] |
|---|---|---|
| TMR/MaP555 | 553 | 570 |
| $JF_{635}$ | 635 | 655 |
| SiR | 650 | 665 |

The dead time estimated by the instrument software (3.7 ms) was subtracted from time points. Background anisotropy corresponding to the free fluorophore was subtracted from the data and at least 8 technical replicates were averaged. Time-dependent anisotropy data was normalized to the highest polarization values (B) and a second order integrated rate equation (**eq. 7**) was fitted to the data yielding $k_1$ as the binding rate constant (on-rate).

$$\mathrm{A} = 1 + \frac{\frac{-1}{[D]_0} \cdot \left([D]_0 \cdot ([D]_0 - [HT7]_0)\right) \cdot e^{([D]_0 - [HT7]_0) \cdot k_1 \cdot t}}{[D]_0 \cdot e^{([D]_0 - [HT7]_0) \cdot k_1 \cdot t} - [HT7]_0} \qquad (\mathbf{7})$$

A: fluorescence anisotropy, $[D]_0$: initial dye concentration, $[HT7]_0$: initial (d)HaloTag7 protein concentration, t: time [s], $k_1$: on-rate [$M^{-1}$ $s^{-1}$].

#### Cell biology and microscopy

All U2OS cell lines (Flp-IN T-REx™ cells (ThermoFisher Scientific), corresponding stable cell lines[17] and CRISPR Vimetin-Halo[31]) were maintained in T-25 flasks (Greiner) in high-glucose Dulbecco's Modified Eagle Medium (DMEM GlutaMAX™, phenol-red, Gibco). Growth medium was supplemented with 10% (v/v) fetal calf serum (FCS) and cells were stored in a humidified tissue culture incubator at 37 °C and 5% $CO_2$. Cells were passaged using phosphate buffered saline (PBS, pH 7.4, Gibco) and TrypLE™ Select Enzyme (1x, phenol-red free, Gibco) every 2-3 days and regularly tested for mycoplasma contamination. Cell titters were determined using a fluidlab R-300 handheld cell counter.

##### Stable cell-line establishment

Stable cell lines were generated using the Flp-IN T-REx™ system[32] in U2OS cells. Cells were grown in T-25 flasks to 80% confluence, pcDNA5/FRT/TO plasmid encoding the gene of interest (GOI) and pOG44 plasmid were co-transfected (1:10). The transfection medium was replaced 16-18 h after transfection with fresh growth medium and stable cell lines were selected using 50 µg $ml^{-1}$ hygromycin B (ThermoFisher Scientific) 24 h post-transfection for 2 days. Cells with genomic transgene integration were eventually selected for high-expression level using fluorescence activated cell sorting (FACS) on a FACSMelody™ Cell sorter (BD Biosciences) using xHTL staining (500 nM). The different cell lines used in this study are summarized in **Method Table 8**.

##### Cell fixation and staining

Cells were fixed after PBS wash using prewarmed 4% (v/v) cell-culture grade paraformaldehyde solution (PFA, Electron Microscopy Sciences) and 0.1% (v/v) glutaraldehyde (GA, Electron Microscopy Sciences) in PBS for 30 min at 37 °C. Excess fixative was quenched using 50 mM ammonium chloride and the cells were subsequently washed with PBS three times. Cells were stored in PBS up to 1 week at 4 °C.

##### Live-cell staining

For live-cell microscopy imaging, 1.0 to 1.5 x $10^5$ cells per well were seeded into tissue culture treated CellCarrier 96 wells black plates (PerkinElmer) or CELLview slide 10 wells (Greiner Bio-One) with an optically clear glass-bottom. The medium was changed 14 h after seeding to regular growth medium, supplemented with 0.1 mg/mL doxycycline to induce protein expression in case of U2OS Flp-IN T-REx™ stable cell lines. Transient transfection was performed using Lipofectamine 3000® reagent (ThermoFisher Scientific) according to the manufacturer's protocol. Live-cell staining was performed 22 h after seeding or transfection. Fluorophore-xHTL were applied in imaging medium (DMEM GlutaMAX™, 10% FCS, phenol-red free, Gibco) at 500 nM concentrations for 2 h at 37 °C prior to imaging, keeping the DMSO concentration below 1%.

#### Confocal Fluorescence Microscopy

Live-cell confocal fluorescence images were acquired on a Leica DMi8 microscope (Leica Microsystems) equipped with a Leica TCS SP8 X scanhead and a SuperK white light laser and a 405 nm diode laser (Hoechst channel). A HC PL APO 40x/1.10 W motCORR CS2 water objective (A) or a HC PL APO 63x/1.40 Oil CS2 oil immersion objective (B) were used in combination with hybrid detectors (HyD). Multi-color images were acquired by sequential imaging to avoid spectral bleed-through. All confocal images were recorded in 1024x1024 pixel resolution (12 bit) with a pixel dwell time of 862 ns and a pinhole size of 1 Airy Unit unless otherwise stated. Imaging settings are summarized in **Method Table 9**. Live cells were maintained in a $CO_2$ (5%) and temperature-controlled (37 °C) chamber (Life Imaging Services).

Pulse-chase experiments were carried out in µ-Slide VI 0.5 Glass Bottom slides (Ibidi) connected to a custom-build perfusion system by addition or replacement of the staining solution (SiR-xHTLs, 500 nM in HBSS, 1x, no $Ca^{2+}$/$Mg^{2+}$, phenol red free, Corning) directly under the microscope and recording of 75 consecutive z-stacks (2 channels, each 10 slices of 2 µm step size) with an imaging speed of 2 stacks per minute. The experiment was repeated three times and the fluorescence intensity of individual nuclei was extracted using the Fiji[33] software and normalized to MaP555-BG/SNAP counter-staining.

Image processing and analysis was performed using the Fiji[33] software where brightness and contrast were adjusted. For image representation, sum or maximum projections were created as indicated. 'Hot' lookup tables (LUT) were applied as indicated. From circular ROIs, the cellular brightness ($R_{FI}$) was measured in H2B-(d)HaloTag7-T2A-mEGFP[17] U2OS cells (n≥150 cells) from 4 fields of view (FOVs) (**eq. 8**) from

- the fluorescence intensities of the dye in the nucleus ($FI_{nuc}^{Dye}$) and
- the fluorescence intensities in the cytosolic mEGFP channel ($FI_{cyto}^{mEGFP}$).

$$R_{FI} = \frac{FI_{nuc}^{Dye}}{FI_{cyto}^{mEGFP}} \qquad (\mathbf{8})$$

$R_{FI}$: Cellular brightness, $FI_{nuc}^{Dye}$: Nuclear dye fluorescence intensity on H2B-(d)HaloTag7, $FI_{cyto}^{mEGFP}$: Cytosolic mEGFP fluorescence intensity.

In H2B-HaloTag7 expressing cells co-seeded with cells not expressing any HaloTag fusion, the signal specificity, *i.e.* signal-over-background ratio (S/B), was calculated from (n≥50 cells, 3 FOVs, **eq. 9**)

- the fluorescence intensities of the dye in the nucleus of (d)HaloTag7 expressing cells ($FI_{HaloTag7}^{Dye,\ nuc}$) and
- the fluorescence intensities of the dye in the nucleus of non-expressing cells ($FI_{emtpy\ cell}^{Dye,\ nuc}$).

$$\text{S/B} = \frac{FI_{HaloTag7}^{Dye,\ nuc}}{FI_{empty\ cell}^{Dye,\ nuc}} \quad (\mathbf{9})$$

S/B: Signal-over-background, $FI_{HaloTag7}^{Dye,\ nuc}$: nuclear dye fluorescence intensity of cells expressing H2B-HaloTag7, $FI_{empty\ cell}^{Dye,\ nuc}$: nuclear dye fluorescence intensity of cells expressing no HaloTag7 fusion.

#### Wide-field Fluorescence Microscopy

Widefield fluorescence imaging was performed on a DMi8 widefield microscope (Leica) equipped with a HC PL APO 20x/0.8 (dry) objective. SiR fluorescence was excited at 635/18 nm and detected using a 700/75 nm filter. The eGFP intensity was recorded using a 474/27 nm excitation and 525/50 nm emission filter (2x2 pixel binning). Live-cells were maintained in a $CO_2$ (5%) and temperature-controlled (37 °C) chamber (PeCon). Imaging settings are summarized in **Method Table 9**.

Live-cell labeling kinetics were measured in µ-Slide VI 0.5 Glass Bottom slides (Ibidi). SiR-xHTLs (25 nM in HBSS, 1x, no $Ca^{2+}$/$Mg^{2+}$, phenol red free, Corning) were flushed in the imaging chambers directly under the microscope while recording Z-stacks (2 channels, 20 stacks, 2 stacks/minute). Signal brightness ($R_{FI}$) was extracted as explained before (**eq. 9**) from n≥15 cells. The data was corrected by the background fluorescence (t = 0) and the maximal final intensity (= 1) for each individual dye and plotted against the imaging time. An exponential association function (**eq. 10**) was fitted to the data to determine the rate-constants k and extract the half-labeling time $\tau_{1/2}^{kin}$ (**eq. 11**).

$$y = A\cdot(1 - e^{-kx}) \quad (\mathbf{10})$$

y: normalized fluorescence intensity, x: time [min], A: amplitude, k: rate-constant [$min^{-1}$].

$$\tau_{1/2}^{kin} = \frac{ln(2)}{k} \quad (\mathbf{11})$$

$\tau_{1/2}^{kin}$: half-labeling time [min], k: rate-constant [$min^{-1}$].

#### PAINT Imaging

PAINT experiments were performed using fixed U2OS cells on µ-Slide VI 0.5 Glass Bottom (Ibidi) at a confluence of 80%. For DNA-PAINT experiments cells were permeabilized using 3% IgG-free BSA and 0.1 % saponin in PBS for 30 min at rt. Afterwards, primary vimentin antibodies (recombinant monoclonal anti-Vimentin, EPR3776, ab92547, abcam) were diluted 1:200 in PBS and added to the chamber. After incubation for 2 h at rt, excess primary antibodies were removed by three washing steps with PBS followed by incubation for 2 h at rt with custom DNA-docking strand labeled secondary antibodies diluted 1:100 in PBS (AffiniPure donkey-anti rabbit IgG (H+L, 711-005-152), 5'-TTCATTACTTCT -3', 1,3 mg $mL^{-1}$). After removing excess secondary antibodies with three washing steps using PBS, cells were post-fixed with 4% PFA for 10 min at rt and finally washed thrice with PBS. Prior to DNA-PAINT measurements, P1-ATTO655 imager strands (P1 5'-AGAAGTAATG-ATTO655-3') were diluted in imaging buffer (500 mM NaCl in PBS, pH 8.3) to a concentration of 1 nM and added to the chambers.

For sample drift correction and image registration fiducial gold markers (125 nm gold beads, Nanopartz, USA) were used. They were diluted 1:30 in PBS, sonicated for 10 min and 100 µL of the solutions were added to the microscopy chambers. After settlement of the gold beads (5 min), the samples were washed thrice with PBS.

For single-colour HT-PAINT and the determination of the single-molecule binding kinetics, fluorescent xHTLs were diluted in PBS to a final concentration of 1 nM and added to the samples. For exchange 2-color PAINT[34] the orthogonal xHTLs were diluted to 3-5 nM in PBS and added to the sample using a microfluidic system (Bruker) at a flow-rate of 600 µL/min, sequentially including a PBS wash step at a flow-rate of 600 µL/min in between. For DNA-PAINT measurements, P1-ATTO655 imager strands were diluted in DNA-PAINT imaging buffer and added to the sample at a concentration of 1 nM.

Single-molecule fluorescence microscopy was performed at the commercial N-STORM super-resolution microsystem (Nikon, Japan). The optical setup was equipped with an oil immersion objective (objective Apo, 100x, NA 1.49) and an EMCCD camera (DU-897U-CS0-#BV, Andor Technology, Ireland). SiR and $JF_{635}$-xHTLs or DNA-PAINT imager strands were excited with a collimated 647 nm laser beam at an intensity of 1.1 kW $cm^{-2}$ (measured at the objective) at highly inclined and laminated optical sheet mode (HILO). 30 000 – 40 000 consecutive frames were acquired. Following parameters were used for image acquisition: camera integration time of 100-150 ms in active frame transfer mode with an EMCCD camera gain of 200, a pre amp gain of 1 and at an effective pixel size of 158 nm. Software tools NIS Elements (Nikon, Japan), LCControl (Agilent, USA), and Micro-Manager 1.4.22[35] were used for optical setup control and image acquisition. For 2-color single-molecule microscopy, a microfluidic system (Bruker, USA) was used for automated exchange of fluorescent ligands.

#### PAINT-Image processing and determination of single-molecule binding kinetics

Localization of single molecules and the reconstruction of super-resolution images were performed with the modular software package Picasso[36]. Single-molecule spots were identified in individual frames using the integrated Gaussian maximum likelihood estimation using the following parameter: Min. net. gradient of 10 000-15 000, a baseline of 205, a sensitivity of 4.78 and a quantum efficiency of 0.95. The images were subsequently corrected for lateral drift using the position of fiducial marker via an integrated redundant cross-correlation algorithm. Single-molecule localizations were then filtered based on SiR, $JF_{635}$ and ATTO655 single-molecule footprints (PSF symmetry $0.6 < FWHM(x)/FWHM(y) < 1.3$) and intensity thresholds. Binding events from the same origin that were spanning over multiple consecutive frames were spatiotemporally linked within a radius of five times the nearest-neighbor based analysis (NeNa[35]) localization precision and allowing a maximum dark time of 5 consecutive frames.

The average binding times (bright time, $\tau_b$) and off-rate constant ($k_{off}$) of the fluorophore-xHTLs were determined from reconstructed super-resolution images. In brief, spatiotemporally linked single binding events (n > 300) were picked manually from reconstructed super-resolution images using the Picasso software. The relative frequency of $\tau_b$ was then plotted in Origin software 2019[37] and fitted with a gaussian distribution to determine the average binding time. $k_{off}$ was then calculated as $1/\tau_b$. For PAINT imaging, 'Hot' LUT were applied for data representation.

#### MINFLUX Microscopy

For MINFLUX experiment, U2OS vimentin-HaloTag7[31] cells were seeded on glass coverslips (precision cover glasses, thickness No. 1.5H, tol. ± 5 µm, 18 mm Ø, Martinsried), grown until a confluence of 60% was reached and chemically fixed as described above. Cells were permeabilized using 0.1% Triton-X100 for 10 min at rt followed by 3x PBS-wash and afterwards incubated with a primary mouse anti-vimentin antibody (Sigma Aldrich, cat. V6389,1:500, 1 h, rt) and a secondary goat anti-mouse, AlexaFluor488-labeled antibody Thermo Fisher, cat. A32723,1:1000, 1 h, rt) with 5 washing steps with PBS w. 1% (w/v) BSA in between and after. Next, the cells were incubated with fiducial markers (5 min, rt, NanoParz, cat. A12-40-980-CTAB-DIH-1-25) washed three times with PBS w. 1% (w/v) BSA and stained with 2 nM SiR-FSAm in PBS w. 1% (w/v) BSA.

MINFLUX imaging was performed on an Abberior 3D MINFLUX (Abberior Instruments GmbH, Göttingen, Germany) built on a motorized inverted microscope IX83 (Olympus, Tokyo, Japan) and equipped with 640 nm and 561 nm MINFLUX laser lines, 488 nm and 405 nm confocal lines. Detection was performed with APDs in the spectral windows 720-685 nm and 650-685 nm. Images were acquired using the default 2D imaging sequence, with an L in the last iteration step of 40 nm, and photon limit of 200 photons. Pinhole was set to 0.67 A.U. and excitation power at the periscope for the first iteration was ~ 35 µW.

Five filters were applied in the post-processing step to avoid false single molecule emission events. To exclude detections originated from transient background, we set the maximum thresholds of 1.2 for the center frequency ratio (CFR) test and 100 kHz for the detected fluorescence rate.[38] Localizations from the same emission trace, *i.e.* with same TiD, further than three standard deviations with respect to the mean trace position were considered outliers and excluded from the trace. Only the traces containing at least 3 localizations within the first 180 minutes of the measurement were considered to calculate the localization precision. The experimental localization precision was estimated by co-aligning the mean values of all localizations obtained from individual emission traces fitted with a Gaussian function to estimate the overall standard deviation.[39] For MINFLUX imaging, 'Hot' LUT were applied for data representation.

#### Live-cell STED Microscopy

Live cells STED nanoscopy was performed using an Abberior STED Expert Line 595/775/RESOLFT QUAD scanning microscope (excitation lines: 355 nm, 405 nm, 485 nm, 561 nm, 640 nm; STED lines: 595 nm and 775 nm; RESOLFT lines: 405 nm, 488 nm) equipped with a UPlanSApo 100x/1.4 oil immersion objective lens (Abberior Instruments, A), or an Abberior STED Infinite line 660/775 QUAD scanning microscope (excitation lines: 520 nm, 561 nm, 640 nm, multiphoton; STED lines: 655 nm and 775 nm) equipped with a 60x/1.42 UPLXAPO60XO oil immersion objective lens (Abberior Instruments, B, both Abberior Instruments GmbH). Detection was performed with avalanche photodiodes (APD) and spectral detection. Fluorophores, microscope and imaging settings are summarized in **Method Table 10**. For STED imaging, 'Hot' LUT were applied for data representation.

STED performance was assessed by measuring the Full Width at Half Maximum (FWHM) of single intermediate filament fibers revealed by Vimentin-HaloTag7. STED images were acquired in an 8x8 µm window with a pixel dwell time of 10 µs and a pixel size of 20 nm with 2 average line scans. Image analysis was performed with *ImageJ*[33] by extracting fluorescence intensity profiles perpendicular to vimentin filaments. Mean filament diameters were calculated from at least ten individual fibrils and from at least three individual images ($n \geq 3$) by fitting a gaussian function (**eq. 12**), yielding FWHM from **eq. 13**.

$$y = y_0 + \frac{A}{\omega\sqrt{\frac{\pi}{2}}} e^{-2\frac{(x-x_C)^2}{\omega^2}} \qquad (\mathbf{12})$$

y: normalized fluorescence intensity, x: x-coordinate [µm], $y_0$: offset, A: area, ω: width, $x_c$: center.

$$FWHM = \omega\sqrt{2\ln(2)} \qquad (\mathbf{13})$$

FWHM: Full Width at Half Maximum, ω: Gauss width.

Photobleaching by multi-frame STED acquisition was recorded in an 8x8 µm window (detailed settings are summarized in **Method Table 10**) and exemplary magnification of single mitochondrial tubules are shown. Bleaching curves were generated by extracting the mean pixel values over the imaging series using *ImageJ.* The image borders were excluded from the evaluation to avoid imaging artefacts. The data was background corrected and normalized to the first frame intensity. Mean intensities from at least three individual experiments ($n \geq 3$) were plotted against the frame number and individually fitted to a mono-exponential decay equation (**eq. 14**) to derive the frame number before reaching half the initial intensity (frame number at half-intensity $\tau_{1/2}$). Mean values and standard deviation of $\tau_{1/2}$ are presented.

$$I = I_0 \cdot e^{\frac{-x}{\tau_{1/2}}} \qquad (\mathbf{14})$$

I: normalized fluorescence intensity, $I_0$: Initial intensity, x: frame number, $\tau_{1/2}$: frame number at half-intensity.
3D-STED images were generated by recording consecutive x-y-frames with 100% axial STED (z-direction).

#### Statistical analysis and reproducibility.

Propagation of uncertainty for variables from products and quotients was calculated using **eq. 15**.

$$\sigma = |C|\sqrt{\left(\frac{\sigma_A}{A}\right)^2 + \left(\frac{\sigma_B}{B}\right)^2} \qquad (\mathbf{15})$$

σ: standard deviation, A, B: variables, $\sigma_A, \sigma_B$: standard deviation of variables A and B, $C = \bar{A} * \bar{B}\ or\ C = \frac{\bar{A}}{\bar{B}}$.

Statistical significance of a sample set over a reference set ($n \geq 3$) was probed by performing two-tailed t-test (spectroscopic characterization of Φ, ε, and $F/F_0$ and confocal as well as STED bleaching $\tau_{1/2}$). A Welch correction for larger sample sizes was used (for cellular $R_{Fl}$ and S/B, $n \geq 50$). Statistical analysis were performed with the *OriginLab*[37] software. p-values > 0.05 were considered as non-significant (n.s.) and $p \leq 0.0025$ as significant as indicated in the respective figure captions.

**Method Table 8.** Plasmids and stable cell lines derived thereof.

| Name | Addgene# | Plasmid | Gene | SCL |
|---|---|---|---|---|
| pET51b(+)_HaloTag7 [a] | 167266 | pET51b(+) | HaloTag7 | - |
| pET51b(+)_dHaloTag7 [a] | 167267 | pET51b(+) | dHaloTag7 = HaloTag7-D106A | - |
| pCDNA5/FRT/TO_ NLS-HaloTag7-SNAP-tag | - | pCDNA5/FRT/TO | NLS-HaloTag7-SNAP-tag | Yes |
| pCDNA5/FRT/TO_H2B-HaloTag7_T2A_mEGFP [c] | 187070 | pCDNA5/FRT/TO | H2B-HaloTag7 and mEGFP | Yes |
| pCDNA5/FRT/TO_H2B-dHaloTag7_T2A_mEGFP | 187071 | pCDNA5/FRT/TO | H2B-dHaloTag7 and mEGFP | Yes |
| pCDNA5/FRT/TO_TOM20-HaloTag7_T2A_mEGFP [b] | 187072 | pCDNA5/FRT/TO | TOM20-HaloTag7 and mEGFP | Yes |
| pCDNA5/FRT/TO_TOM20-dHaloTag7_T2A_mEGFP | 187073 | pCDNA5/FRT/TO | TOM20-dHaloTag7 and mEGFP | Yes |
| pCDNA5/FRT/TO_LifeAct-HaloTag7_T2A_mEGFP | - | pCDNA5/FRT/TO | LifeAct-HaloTag7 and mEGFP | - |
| pCDNA5/FRT/TO_Vimentin-HaloTag7_T2A_mEGFP [c] | 187074 | pCDNA5/FRT/TO | Vimentin-HaloTag7 and mEGFP | Yes |
| pCDNA5/FRT/TO_Vimentin-dHaloTag7_T2A_mEGFP | 187075 | pCDNA5/FRT/TO | Vimentin-dHaloTag7 and mEGFP | Yes |
| pCDNA5/FRT/TO_H2B-HaloTag7 [c] | 169329 | pCDNA5/FRT/TO | H2B-HaloTag7 | Yes |
| pCDNA5/FRT/TO_H2B-dHaloTag7 | - | pCDNA5/FRT/TO | H2B-dHaloTag7 | Yes |
| pCDNA5/FRT/TO_TOM20-HaloTag7 [c] | 169330 | pCDNA5/FRT/TO | TOM20-HaloTag7 | Yes |
| pCDNA5/FRT/TO_TOM20-dHaloTag7 | - | pCDNA5/FRT/TO | TOM20-dHaloTag7 | Yes |
| pCDNA5/FRT/TO_NES-HaloTag7 [c] | - | pCDNA5/FRT/TO | NES-HaloTag7 | - |
| pCDNA5/FRT/TO_Ig-κ-HaloTag7-PDGFR [c] | - | pCDNA5/FRT/TO | Ig-κ-HaloTag7-PDGFR | - |
| pCDNA5/FRT/TO_Ig-κ-dHaloTag7-PDGFR | - | pCDNA5/FRT/TO | Ig-κ-dHaloTag7-PDGFR | - |
| pCDNA5/FRT/TO_CalR-HaloTag7-KDEL [c] | - | pCDNA5/FRT/TO | CalR-HaloTag7-KDEL | - |
| pCDNA5/FRT/TO_CalR-dHaloTag7-KDEL | - | pCDNA5/FRT/TO | CalR-dHaloTag7-KDEL | Yes |
| pCDNA5/FRT/TO_LamP1-HaloTag7 [c] | - | pCDNA5/FRT/TO | LamP1-HaloTag7 | - |
| pCDNA5/FRT/TO_SKL-HaloTag7 [c] | - | pCDNA5/FRT/TO | SKL-HaloTag7 | - |
| pCDNA5/FRT/TO_CEP41-HaloTag7 [b] | - | pCDNA5/FRT/TO | CEP41-HaloTag7 | Yes |
| pCDNA5/FRT/TO_Lifeact-HaloTag7 [c] | - | pCDNA5/FRT/TO | Lifeact-HaloTag7 | - |
| pCDNA5/FRT/TO_Lifeact-dHaloTag7 | - | pCDNA5/FRT/TO | Lifeact-dHaloTag7 | Yes |
| pCDNA5/FRT/TO_TOM20-dHaloTag7_T2A_Vimentin-HaloTag7 | - | pCDNA5/FRT/TO | TOM20-dHaloTag7 and Vimentin-HaloTag7 | Yes |
| pCDNA5/FRT/TO_TOM20-dHaloTag7_T2A_Lifeact-HaloTag7 | 187076 | pCDNA5/FRT/TO | TOM20-dHaloTag7 and Lifeact-HaloTag7 | - |
| pCDNA5/FRT/TO_TOM20-dHaloTag7_T2A_Cox8-HaloTag7 | 187077 | pCDNA5/FRT/TO | TOM20-dHaloTag7 and Cox8-HaloTag7 | Yes |
| pCDNA5/FRT/TO_TOM20-dHaloTag7_T2A_LamP1-HaloTag7 | 187078 | pCDNA5/FRT/TO | TOM20-dHaloTag7 and LamP1-HaloTag7 | Yes |
| pCDNA5/FRT/TO_TOM20-dHaloTag7_T2A_CalR-HaloTag7-KDEL | 187079 | pCDNA5/FRT/TO | TOM20-dHaloTag7 and CalR-HaloTag7-KDEL | Yes |
| CRISPR Vimentin-HaloTag [d] | - | - | Vimentin-HaloTag7 | / |

Plasmid/cell-line published in [a] Wilhelm, Kühn *et al.* (2021)[1], [b] Frei *et al.* (2019)[16], [c] Frei *et al.* (2021)[17] and [d] Butkevich *et al.* (2018)[40].
SCL = Stable cell lines U2OS Flp-In TREx, SNAP$^f$-tag = SNAP-tag with fast mutation E30R.

**Method Table 9**. Conventional fluorescence microscopy data acquisition parameters.

| Figure | Label | Ligand | Set-up | Obj. | Excitation [nm] (%) | Emissinon [nm] | Comment |
|---|---|---|---|---|---|---|---|
| 1D | H2B-HaloTag7 | MaP555-FSAm | A | A | 550 (16) | 560 - 610 | Sum projections |
| | | MaP618-HSAm | | | 620 (10) | 630 - 790 | |
| | | $JF_{635}$-HSAm | | | 635 (3) | 645 - 780 | |
| 2D | H2B-dHaloTag7 | MaP555-Hy5 | A | A | 550 (16) | 560 - 610 | Sum projections |
| | | MaP618-Hy4 | | | 620 (10) | 630 - 790 | |
| | | SiR-Hy4 | | | 635 (3) | 645 - 780 | |
| 2E | H2B-dHaloTag7 | MaP555-Hy5 | A | A | 550 (3) | 560 - 600 | Sum projections<br>Seq. imaging |
| | TOM20-HaloTag7 | SiR-HSAm | | | 590 (12) | 600 - 635 | |
| | LamP1-$SNAP^{f}$-tag | $JF_{585}$-BG | | | 635 (13) | 645 - 680 | |
| | Actin | SiR700 | | | 690 (12) | 720 - 795 | |
| S6A | H2B-HaloTag7 | $JF_{525}$-FSAm | A | A | 525 (5) | 535 - 600 | Sum projections |
| | | MaP555-FSAm | | | 550 (16) | 560 - 610 | |
| | | $JF_{585}$-HSAm | | | 585 (3) | 595 - 685 | |
| | | MaP618-HSAm | | | 620 (10) | 630 - 790 | |
| | | $JF_{635}$-HSAm | | | 635 (3) | 645 - 780 | |
| | | SiR700-HSAm | | | 670 (10) | 680 - 795 | |
| S6B | H2B-HaloTag7-T2A-mEGFP | SiR-HSAm | A | A | 633 (3) | 650 - 790 | Ratiometric and sum projection |
| | | mEGFP | | | 488 (50) | 500 - 540 | |
| S7A | NES-HaloTag7 | SiR-HSAm | A | A | 633 (2) | 650 - 790 | Sum projections<br>2 Lines average |
| | IgK-HaloTag7-PDGFR | | | B | 633 (10) | | |
| | CalR-HaloTag7-KDEL | | | A | 633 (3) | | |
| | LamP1-HaloTag7 | | | A | 633 (10) | | |
| | TOM20-HaloTag7 | | | B | 633 (3) | | |
| | SKL-HaloTag7 | | | B | 633 (15) | | |
| | Vimentin-HaloTag7 | | | A | 633 (3) | | |
| | CEP41-HaloTag7 | | | B | 633 (20) | | |
| | Hoechst | | | A/B | 405 (2) | 432 - 500 | |
| S7B | H2B-HaloTag7 | SiR-HSAm | A | A | 633 (3) | 650 - 790 | Confocal plane |
| | | | B | | 635 (20) | 663 - 738 | z-stack |
| S7D | NLS-HaloTag7-SNAP-tag | SiR-HSAm | A | A | 640 (4) | 650 - 690 | Ratiometric and sum projection |
| | | BG-MaP555 | | | 550 (4) | 560 - 620 | |
| S13A | H2B-dHaloTag7 | $JF_{525}$-Hy5 | A | A | 530 (4) | 540 - 600 | Sum projections |
| | | MaP555-Hy5 | | | 550 (4) | 560 - 600 | |
| | | $JF_{585}$-Hy4 | | | 585 (3) | 595 - 685 | |
| | | MaP618-Hy4 | | | 620 (10) | 630 - 790 | |
| | | $JF_{635}$-Hy4 | | | 635 (1) | 645 - 780 | |
| | | SiR700-Hy4 | | | 670 (10) | 680 - 795 | |

| | | | | | | | |
|---|---|---|---|---|---|---|---|
| | IgK-dHaloTag7-PDGFR | TMR-Hy5 | | | 550 (2) | 560 -600 | |
| | TOM20-HaloTag7 | SiR700-HSAm | | | 680 (4) | 680 - 795 | |
| S13B | H2B-dHaloTag7 | CRh-Hy4 | A | A | 600 (3) | 610 - 650 | Sum projections |
| | LifeAct-HaloTag7 | SiR-HSAm | | | 633 (2) | 650 - 795 | |
| | TOM20-dHaloTag7 | MaP555-Hy5 | | | 550 (1) | 560 - 620 | |

Microscopes: **A** SP8-FALCON (confocal), **B** DMi8 (widefield).

**Method Table 10.** STED microscopy data acquisition parameters.

| Fig. | Label | Ligand | Set-up | Excitation [nm] (%) | STED [nm] (%) | Pixel dwell time [µs] | Pixel size [nm] | Size [µm] | Emission [nm] | Comment |
|---|---|---|---|---|---|---|---|---|---|---|
| 1F | Vim-HaloTag7 | SiR-FSAm | A | 640 (2) | 775 (15) | 10 | 20 | 8 x 8 | 650 - 757 | 2 Lines accu., STED & Conf. |
| 1G | TOM20-HaloTag7 | $JF_{635}$-HSAm<br>$JF_{635}$-CA | A | 640 (4) | 775 (20) | 10 | 30 | 10 x 10 | 650 - 757 | 2 Lines accu, 4 frame/min |
| | LamP1-HaloTag7 | SiR-HSAm | | 640 (2) | 775 (20) | | | | 650 - 757 | |
| 2G | TOM20-dHaloTag7 | CRh-Hy4 | B | 561 (40) | 660 (20) | 10 | 30 | 10 x 10 | 571 - 630 | 2 Lines accu. 2 frames/min |
| | LamP1-HaloTag7 | SiR-HSAm | | 640 (2) | 775 (20) | | | | 650 - 757 | |
| 2I | TOM20-dHaloTag7 | CRh-Hy4 | A | 561 (40) | 775 (20) | 10 | 20 | 2.4 x 3.2 | 575 - 630 | 40 x 50 nm z-stacks, 3 Lines accu. 100% 3D-STED |
| | LamP1-HaloTag7 | SiR-HSAm | | 640 (4) | 775 (10) | | | | 670 - 757 | |
| S10A | Vim-HaloTag7 | MaP555-FSAm | B | 561 (10) | 660 (10) | 15 | 30 | 10 x 10 | 571 -650 | 2 Lines accu., STED & Conf. |
| | | MaP618-HSAm | | 640 (6) | | | | | | |
| | | $JF_{635}$-HSAm | A | 640 (6) | 775 (15) | 15 | 30 | 10 x 10 | 571 -693 | |
| | | SiR-HSAm | | 640 (2) | | | | | | |
| S10B | Vim-HaloTag7 | SiR-FSAm<br>SiR-CA | A | 640 (2) | 775 (15) | 10 | 20 | 8 x 8 | 650 - 757 | 2 Lines accu., STED & Conf. |
| S10E | TOM20-HaloTag7 | MaP555-FSAm<br>MaP555-CA<br>$JF_{585}$-CA | B | 561 (2) | 660 (30) | 15 | 30 | 15 x 15 | 650 | 3 Lines accu., 0.2 frames/min |
| S10F | TOM20-HaloTag7 | MaP618-HSAm<br>MaP618-CA | A | 640 (6) | 775 (20) | 15 | 30 | 10 x 10 | 650 - 757 | 2 Lines accu, 1 frame/min |
| S10G | TOM20-HaloTag7 | SiR-HSAm<br>SiR-CA | A | 640 (4) | 775 (20) | 10 | 30 | 10 x 10 | 650 - 757 | 2 Lines accu, 4 frames/min |
| S14C | Vim-dHaloTag7 | MaP555-Hy5 | B | 561 (2) | 660 (30) | 15 | 30 | 15 x 15 | 650 | 3 Lines accu.,0.2 frames/min |
| | | MaP618-Hy4 | | 640 (6) | 775 (20) | 15 | 30 | 10 x 10 | 650 - 757 | 2 Lines accu, 1 frames/min |
| | | $JF_{635}$-Hy4 | A | 640 (4) | 775 (20) | 10 | 30 | 10 x 10 | 650 - 757 | 2 Lines accu., 4 frames/min |
| | | SiR-Hy4 | | 640 (4) | 775 (20) | 10 | 30 | 10 x 10 | 650 - 757 | 2 Lines accu, 4 frames/min |
| S15B | TOM20-dHaloTag7 | MaP555-Hy5 | B | 561 (4) | 775 (15) | 10 | 100/40 | 80 x 80 | 571 - 630 | 3 Lines accu., Conf (overview) & STED (zoom) |
| | LamP1-HaloTag7 | SiR-HSAm | | 640 (2) | 775 (20) | 10 | | 10 x 10 | 650 – 757 | |
| S15C | Sees Fig. 2G | | | | | | | | | |
| S15D | TOM20-dHaloTag7 | MaP618-Hy4 | A | 561 (40) | 775 (20) | 10 | 20 | 2.4 x 3.2 | 575 - 630 | 40 x 50 nm z-stacks, 3 Line accu. 100% 3D-STED |
| | LamP1-HaloTag7 | SiR-HSAm | | 640 (4) | 775 (10) | | | | 670 – 757 | |

Microscopes: **A** STED Expert line with 660/775 depletion lasers, **B** STED Expert Line with 595/775 depletion lasers.

### Supplementary Figures

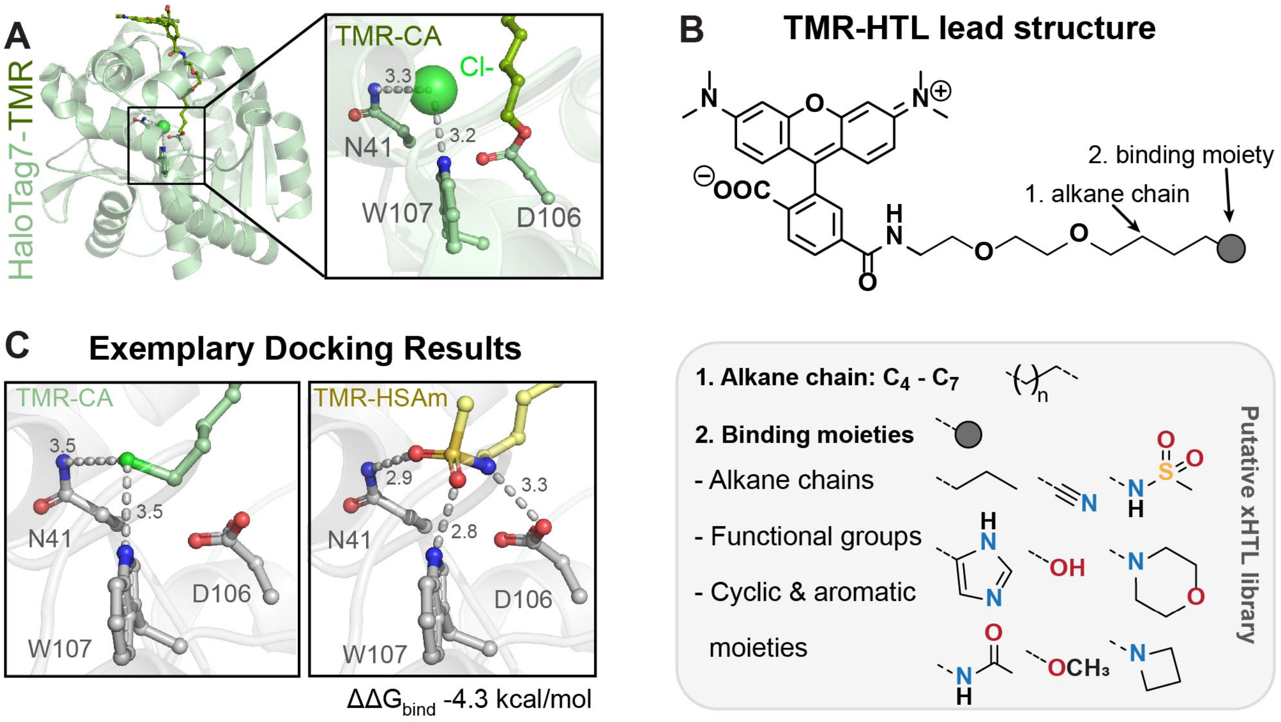

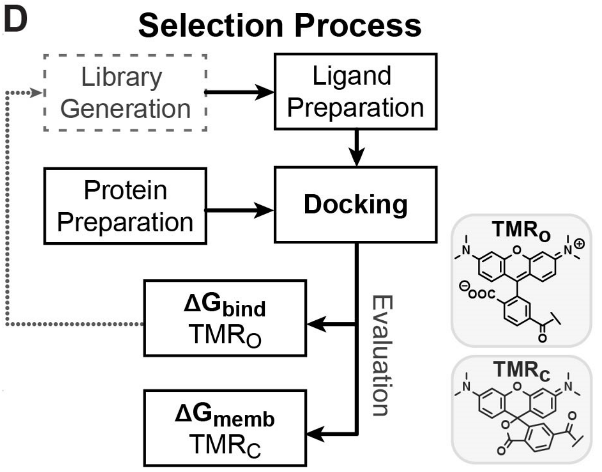

**E**

| TMR-PEG$_2$- | $\Delta\Delta G_{bind}$ TMR$_o$ [kcal/mol] | $\Delta\Delta G_{memb}$ TMR$_c$ [kcal/mol] |
|---|---|---|
| $C_5$-NH-$SO_2$-$CH_3$ | -4.3 | 4.6 |
| $C_5$-$NH_2$ | -3.3 | 8.5 |
| $C_4$-NH-$SO_2$-$CH_3$ | -2.3 | 6.6 |
| $C_6$-Cl (HTL) | 0 | 0 |
| $C_5$-OH | 4.2 | 3.9 |
| $C_4$-OH | 5.5 | 5.9 |
| $C_4$-$NH_2$ | 6.4 | 8.9 |
| $C_4$-CN | 7.1 | 2.5 |
| $C_6$ | 27.4 | -0.3 |
| $C_5$-$CF_3$ | 35.4 | -1.3 |

**Figure S. 1 | Computational screening of xHTLs candidates.**

**A.** Crystal structure of HaloTag7-TMR (PDB: 6Y7A)[1] with zoom onto the active site residues. HaloTag7 is represented as cartoon while the alkane-PEG-TMR and important residues are represented as sticks. Distances are indicated in Å. HaloTag7-TMR was used as a target for docking tentative xHTLs. **B.** TMR ligand chemical structure, which was used as lead structure for dockings. Structural elements that were altered for ligand design are represented in the lower insert, namely the terminal alkane chain length (1) and binding moiety (2) generating 2000 xHTLs candidates. **C.** Exemplary result from Glide docking in standard precision configuration. Docking poses comparison at the active site of TMR-HTL and TMR-HSAm. Relative improvement of the binding energy $\Delta\Delta G_{bind}$ is given below (see D for definition). Distances are indicated in Å. Molecular docking into HaloTag7 yielded poses comparable to TMR-CA redocking. The similarity of the poses allows to compare their relative predicted free Gibbs binding energies ($\Delta G_{bind}$) from Molecular Mechanics/Generalized Born Surface Area (MM-

GBSA) calculations used as a proxy for anticipated relative binding affinities. **D.** Computational screening process. $\Delta G_{bind}$ predicted from docking was calculated for TMR ligands in the open form ($TMR_o$). Membrane permeability $\Delta G_{memb}$[3] was predicted for closed TMR variants ($TMR_c$). All descriptors were evaluated in comparison to redocking of TMR-HTL such that $\Delta\Delta G = {}^{TMR\text{-}HTL}\Delta G - {}^{candidate}\Delta G$. **E.** Table summarizing the molecular descriptors for 9 exemplary xHTL candidates which were ranked by $\Delta\Delta G_{bind}$ considering $\Delta\Delta G_{memb}$. $\Delta\Delta G_{bind}$ from induced-fit docking ranged from 35.4 (worst binding) to -4.3 kcal/mol (best binding). $\Delta\Delta G_{memb}$ ranged from 8.5 (poor permeability) to -1.3 kcal/mol (great permeability). Best $\Delta\Delta G_{bind}$ were obtained for $C_4$- or $C_5$-aliphatic linkers terminating with hydrogen bond donor moieties such as amines or sulfonamides ($\Delta\Delta G_{bind}$ -4.3 to -2.3 kcal/mol). While primary amines were predicted as good binders ($\Delta\Delta G_{bind}$ = -3.3 kcal/mol), they were discarded from further study due to their predicted low cell permeability ($\Delta\Delta G_{memb}$ +8.5 kcal/mol) and literature-reported metabolic conversion of primary amine drugs.[41] Fluorinated ligands such as trifluorosulfonamides, not initially present in virtual screening, were later considered as they are reported to improve cell permeability.[42]

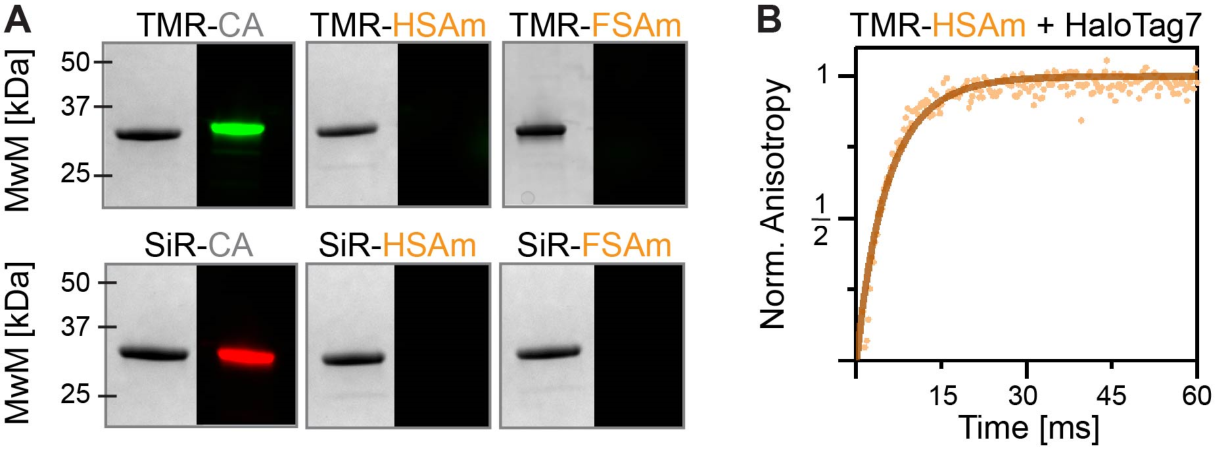

**Figure S. 2 | *In-vitro* characterization of xHTL candidates.**
**A.** HaloTag7 covalent labeling experiments. SDS-PAGE followed by in-gel fluorescence scan and Coomassie-staining. Fluorophore derivatives of CA/HSAm/FSAm ligand were incubated with HaloTag7. xHTLs show non-covalent binding to HaloTag7 and therefore migrated through the gel in contrast to HTL covalent labeling (Dye-CA). MwM – Molecular weight marker. **B.** Binding kinetics of TMR-HSAm to HaloTag7. Kinetics were measured by stopped-flow fluorescence anisotropy. Non-linear fit of a second order binding model to averaged and normalized anisotropy traces (n≥10) yielded the binding rate constant ($k_{on}$) of $6.4 \pm 0.1 \cdot 10^6\ M^{-1}\ s^{-1}$ (mean ± standard fit error).

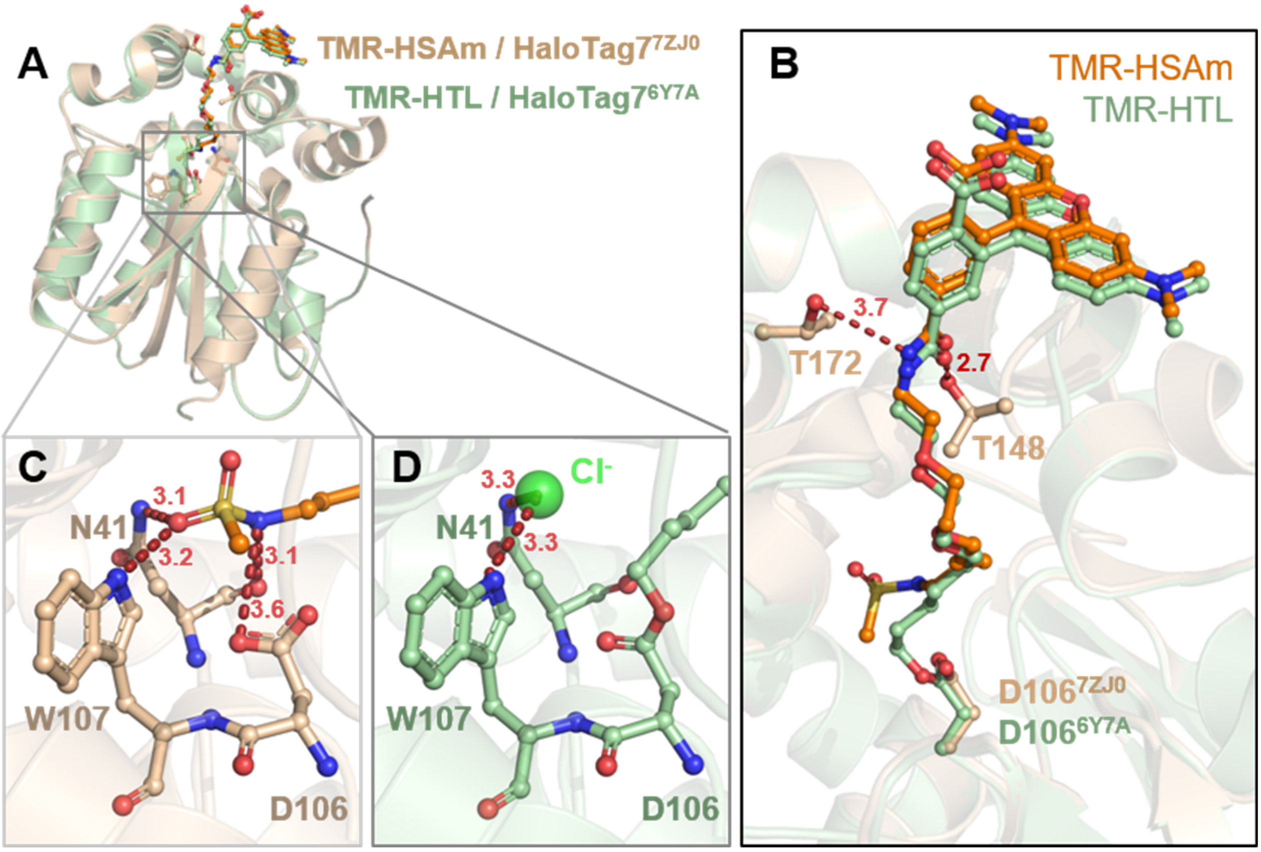

**Figure S. 3 | Structural analysis of the TMR-HSAm/HaloTag7 complex.**

**A.** Structural comparison between the TMR-HSAm/HaloTag7 complex (PDB-ID: 7ZJ0, 1.5 Å resolution) and TMR-HTL/HaloTag7 covalent complex (PDB-ID: 6Y7A, 1.4 Å resolution)[1]. Tertiary structures are represented as cartoons. The ligands and relevant residues are represented as sticks. **B.** Magnification on the TMR-ligands binding sites. Distances in Å. The TMR moieties are located at the protein's surface while the alkane chains are buried in the protein hydrophobic tunnel as previously described.[1] **C.** Magnification on the HSAm binding site. Polar interactions between HSAm and HaloTag7 residues: TMR-HSAm amide moiety interacts with D106 while the sulfonamide group occupies the chloride binding pocket between N41 and W107. **D.** Magnification on the covalent bond between TMR-HTL and the HaloTag7 protein. The TMR-labeled HaloTag7 features a chloride ion (green sphere) inside the active center where the sulfonamide is located in the HSAm complexed structure.

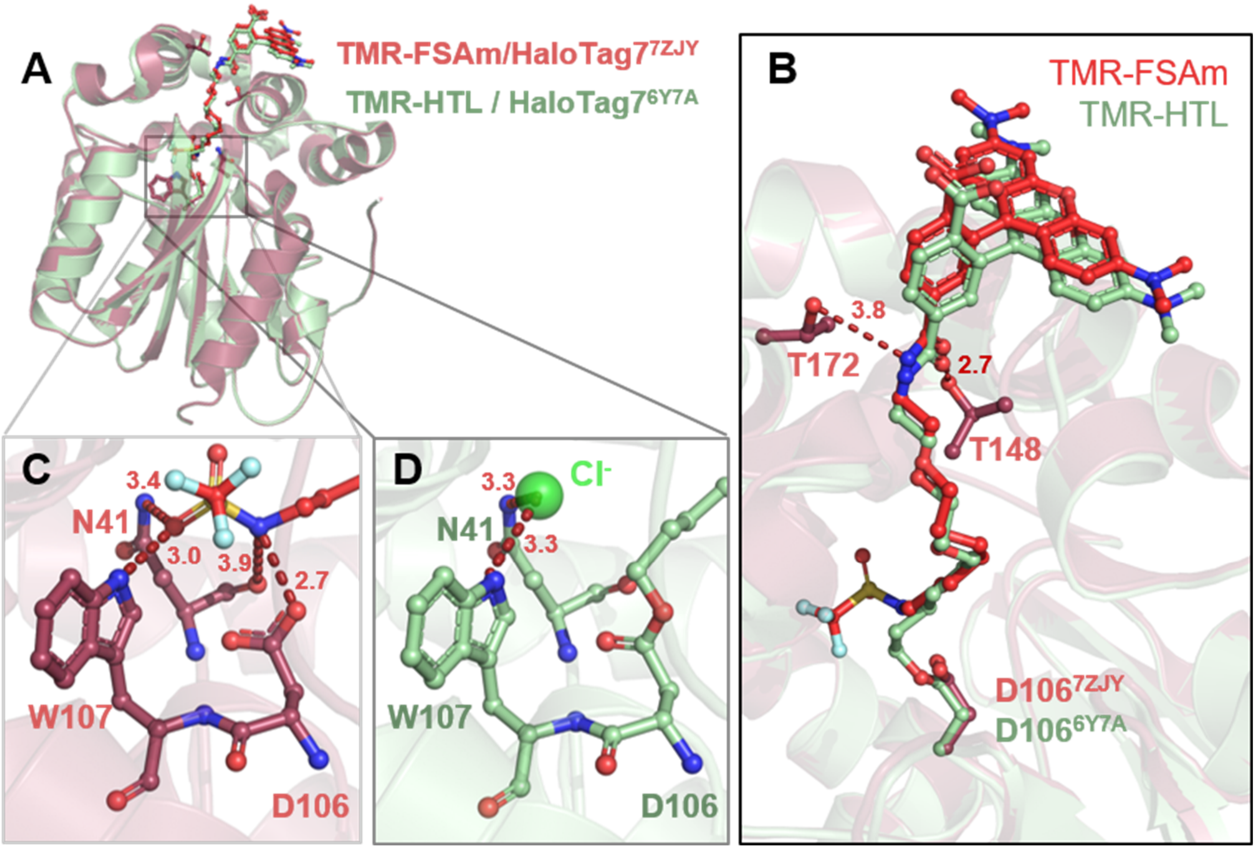

**Figure S. 4 | Structural analysis of the TMR-FSAm/HaloTag7 complex.**
**A.** Structural comparison between the TMR-FSAm/HaloTag7 complex (PDB-ID: 7ZJ0, 1.5 Å resolution) and TMR-HTL/HaloTag7 covalent complex (PDB-ID: 6Y7A, 1.4 Å resolution)[1]. Tertiary structures are represented as cartoons. The ligands and relevant residues are represented as sticks. **B.** Magnification on the TMR-ligands binding sites. Distances in Å. The TMR moieties are located at the protein's surface while the alkane chains are buried in the protein hydrophobic tunnel as previously described.[1] **C.** Magnification on the FSAm binding site. Polar interactions between FSAm and HaloTag7 residues: TMR-FSAm amide moiety interacts with D106 while the sulfonamide group occupies the chloride binding pocket between N41 and W107. **D.** Magnification on the covalent bond between TMR-HTL and the HaloTag7 protein. The TMR-labeled HaloTag7 features a chloride ion (green sphere) inside the active center where the sulfonamide is located in the FSAm complexed structure.

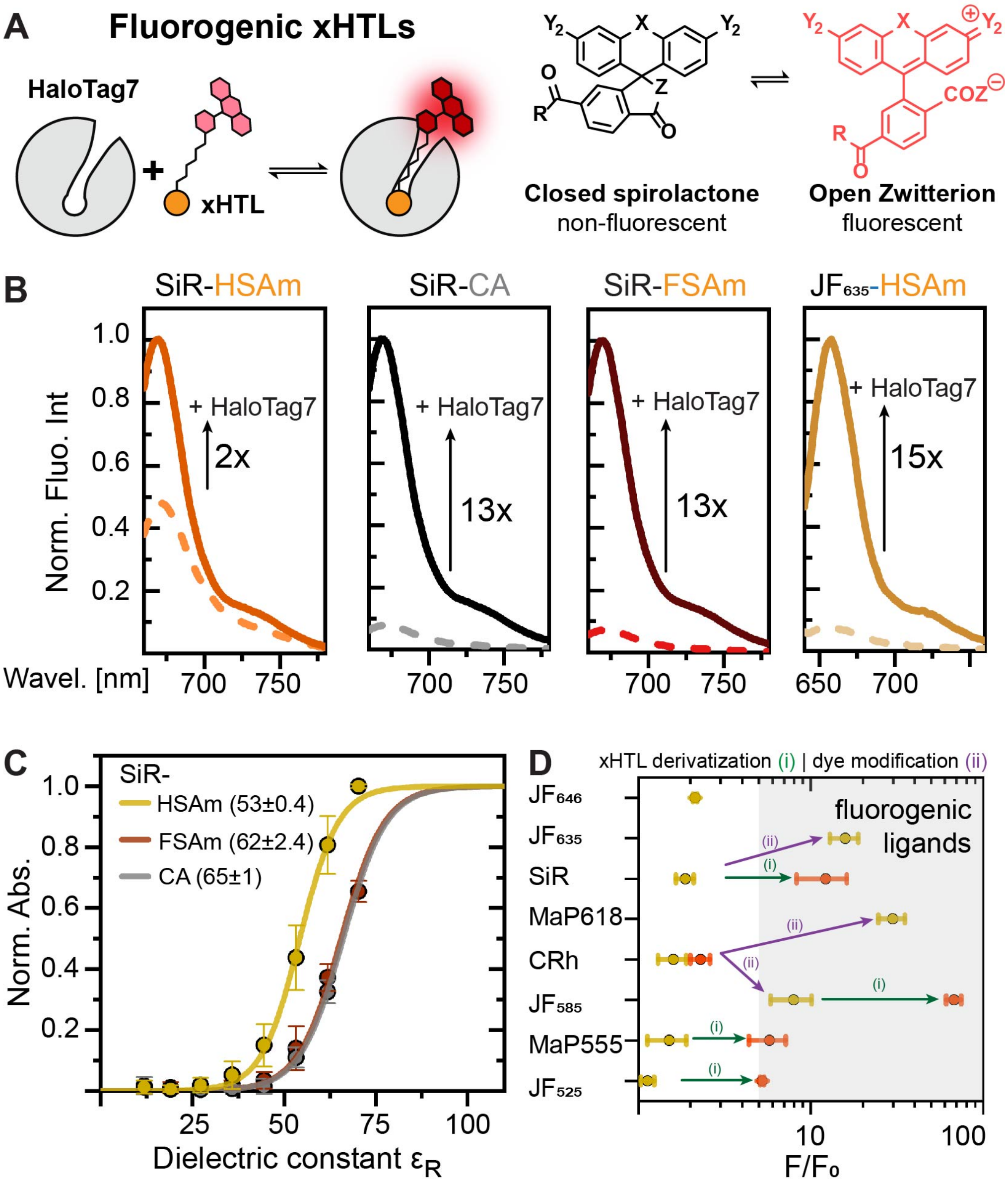

**Figure S. 5 | Spectral and fluorogenic properties tuning of xHTLs.**

**A.** Scheme of fluorogenic xHTL fluorescence intensity increase upon HaloTag7 binding. Dynamic equilibrium and general chemical structure of fluorogenic rhodamine dyes used in this study. X: O, $C(CH_3)_2$, $Si(CH_3)_2$. Y: dimethylamine, azetidine, 3,3-difluoroazetidine, 3,3-tetrafluoroazetidine. Z: -OH, $NH\text{-}SO_2\text{-}N(CH_3)_2$. **B.** Fluorescence emission spectra comparison between free (dashed lines) and HaloTag7-bound (plain line) SiR/$JF_{635}$-(x)HTLs. Arrows indicate fluorescence intensity increase ($F/F_0$) upon protein binding. SiR-HSAm exhibits a low fluorescence increase upon protein binding which was restored by introducing SiR-FSAm derivative or using the more fluorogenic dyes with similar spectra properties *i.e.* $JF_{635}$. **C.** SiR-(x)HTL water-dioxane titration curves. Sigmoidal fit to averaged (n=3) and normalized (to max. absorbance from SiR-HSAm) data yielded the $D_{50}$-values (in brackets ± standard deviation), demonstrating that SiR-HSAm (53.0 ± 0.4) was less fluorogenic than SiR-FSAm (62.3 ± 2.4). **D.** xHTL fluorescence intensity increase ($F/F_0$) upon HaloTag7 binding of Dye-HSAm (yellow) and -FSAm (red). Colored arrows indicate which strategy was used to improve the xHTL fluorogenicity by using either FSAm derivatization (i – green) or dye modification (ii – purple). Mean values (n>3) and standard deviation presented. xHTLs with a $F/F_0 > 5$ were considered as fluorogenic.

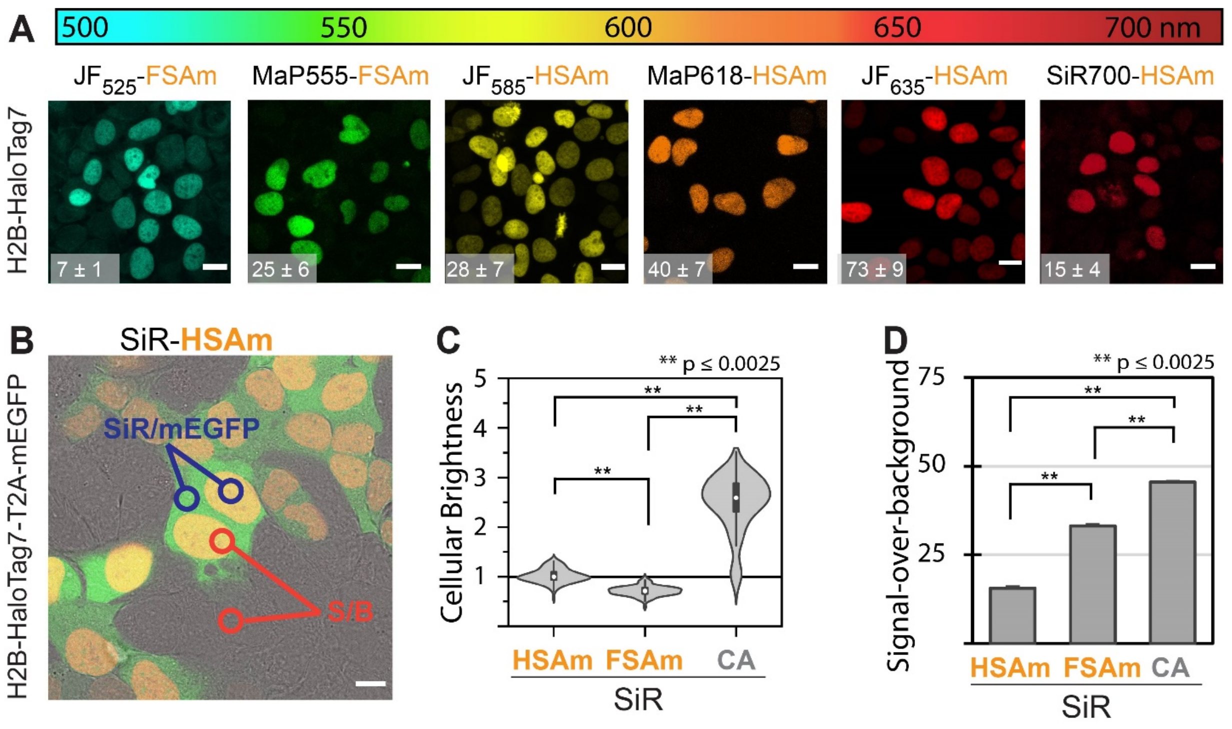

**Figure S. 6 | Live-cell staining characterization of xHTLs.**
**A.** Live-cell confocal images of different fluorescent xHTLs probes covering the visible spectrum. H2B-HaloTag7 expressing U2OS cells stained with 500 nM xHTLs and imaged via live-cell confocal microscopy. Sum projections. Scale bars: 10 µm. Signal-over-background ratios (S/B) were calculated by dividing nuclear fluorescence intensities from cells expressing HaloTag fusions by non-expressing cells ($FI^{SiR,\ nuc}_{HT7}/FI^{SiR,\ nuc}_{empty\ cell}$, n ≥ 50 cells, from at least two individual experiments, mean values ± standard error of the mean). Values are given in bottom-left corners. **B.** Confocal image illustrating the calculation of parameters employed to evaluate the SiR-xHTL staining quality. Live-cell confocal image of H2B-HaloTag7-T2A-mEGFP expressing U2OS cells stained with 500 nM SiR-HSAm. Overlay of SiR (red), eGFP (green) and bright-field (gray) channels, sum projections. The fluorescence intensity from single nuclei was normalized by the cytosolic mEGFP fluorescence intensity of the same cell (expression control) to assess the cellular brightness. The signal-to-background ratio (S/B) was evaluated as explained in A. Scale bar: 10 µm. **C.** Comparison of cellular brightness obtained for different SiR-(x)HTLs. Staining using 500 nM, no-wash. Live U2OS cells, n ≥ 150 cells, 4 images from at least two individual experiments. SiR-HSAm shows superior cellular brightness than SiR-FSAm (31 ± 3% on HaloTag7) but the covalent staining remains generally brighter by a factor of 2 – 3 compared to any xHTL staining. Violin plot = light grey, whisker plot = dark grey, box = 25%–75% percentile and whiskers = 5%–95% percentile, circle = mean. Populations were normalized to the mean brightness of SiR-HSAm (horizontal reference line). Significance was calculated using two-sided t-tests including the Welch correction, not significant (n.s.): p > 0.05 (*), significant: p ≤ 0.0025 (**). **D.** Comparison of live cell S/B of SiR-(x)HTL stains under confocal imaging conditions (n ≥ 150 cells, 4 images from at least two individual experiments, mean values ± S.E.M.). SiR-FSAm shows improved S/B by 112 ± 2% over SiR-HSAm but remains nevertheless 20 ± 1% lower than HaloTag7 covalent labeling.

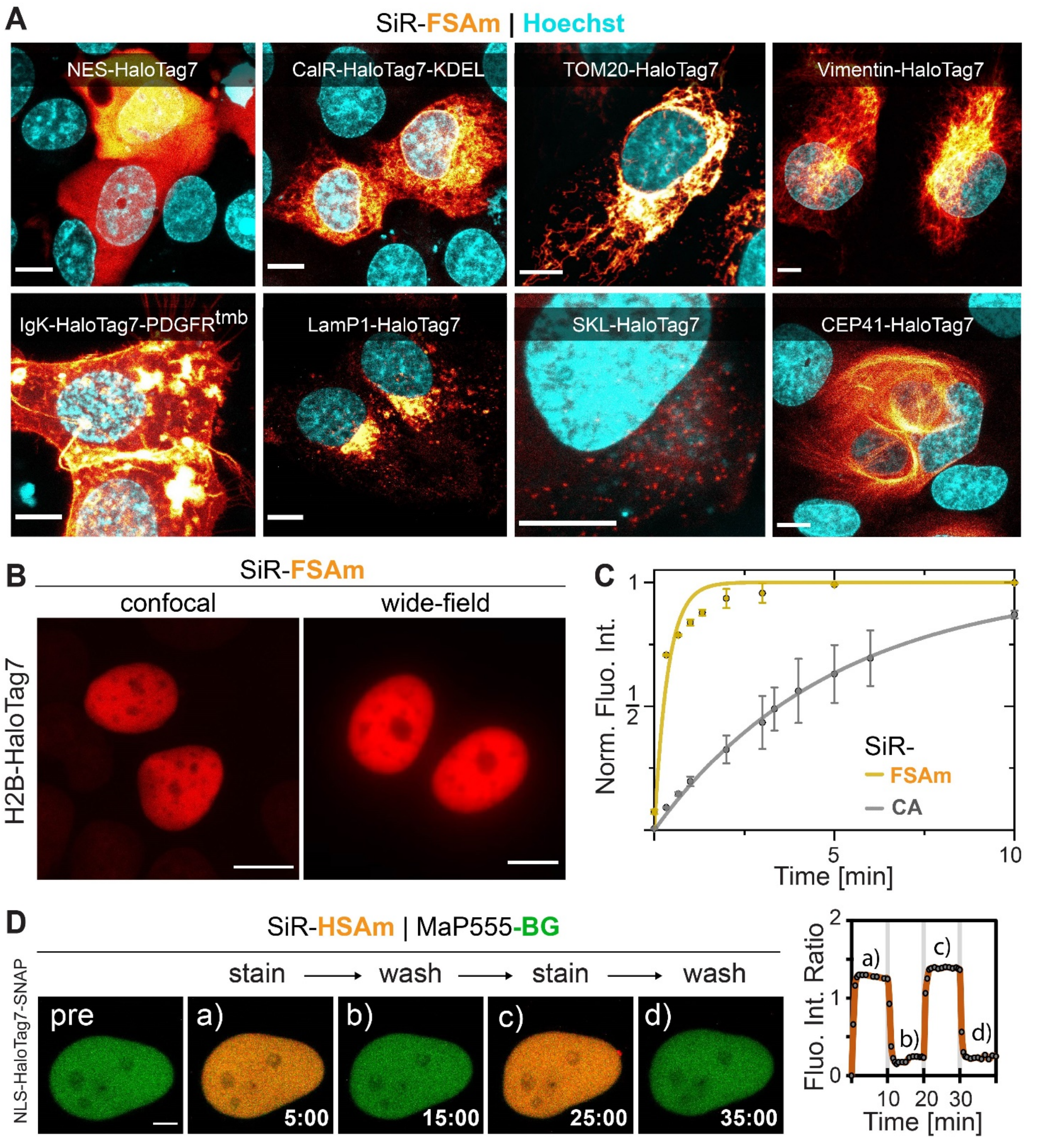

**Figure S. 7 | Applications of xHTLs for live-cell confocal microscopy.**
**A.** Images of sub-cellular structures in U2OS cells. HaloTag7-fusion protein overexpressed, stained with SiR-HSAm (500 nM) and Hoechst (1 µg/mL) and live imaged by confocal microscopy (max. projection). Localizations (fusion protein): Cytoplasm (NES), outer-plasma membrane (IgK/PDGFRtmb), endoplasmic reticulum (CalR/KDEL), lysosomes (LamP1), mitochondria surface (TOM20), peroxisomes (SKL), intermediate filaments (vimentin) and tubulin (CEP41). Scale bars: 10 µm. 'Hot' LUT were applied. **B.** SiR-FSAm staining under confocal (500 nM) and wide-field (100 nM) microscopy conditions. Live U2OS cells expressing H2B-HaloTag7. Scale bars: 5 µm. SiR-FSAm enables to image cells in wide-field microscopy despite the impossibility to conduct washing steps. **C.** Live-cell staining kinetics comparing covalent HTL and non-covalent xHTLs. U2OS express

H2B-HaloTag7 stained with 25 nM SiR-(x)HTLs and imaged on a wide-field microscopy. Signal was normalized to maximum final signal obtained after 20 min. Average data from n ≥ 15 cells from one experiment. Data fitted with a mono-exponential association function yielding the half-labeling times $\tau_{1/2}^{kin}$ (mean ± standard fitting error). SiR-FSAm: 0.2 ± 0.1 min, SiR-CA: 2.9 ± 0.6 min. **D.** Reversible cellular staining using xHTLs. Live-cell confocal images of U2OS expressing NLS-HaloTag7-SNAP-tag covalently labeled with MaP555-BG and iteratively stained with 500 nM SiR-HSAm or washed with imaging medium (10 min cycles). Scale bar: 2 µm. Images recorded every 30 s. Intensity-time-trace given on the right panel. Fluorescence intensity ratio: $FI_{SiR}$ / $FI_{MaP555}$.

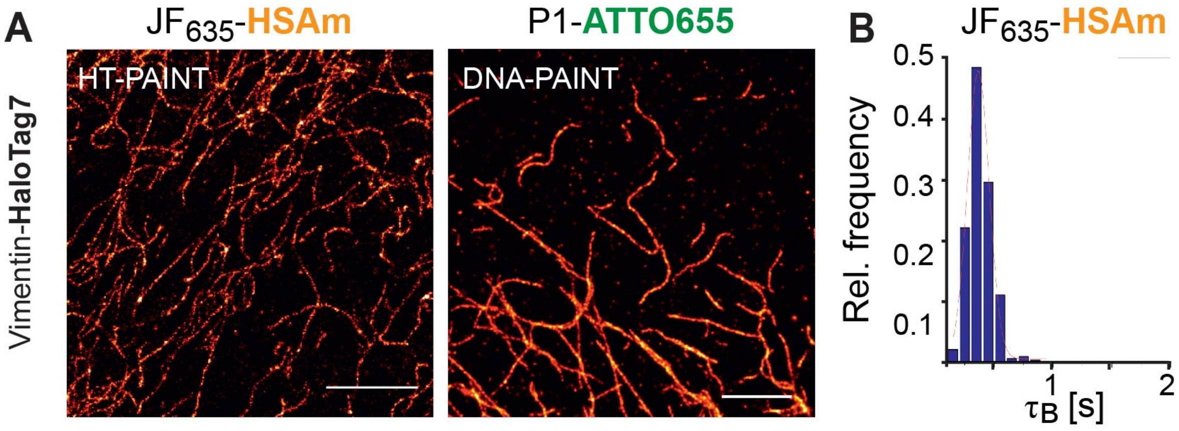

**Figure S. 8 | Comparison of HT-PAINT and DNA-PAINT.**
**A.** Comparison of HT- and DNA-PAINT imaging. For HT-PAINT, fixed U2OS cell endogenously expressing vimentin-HaloTag7[31] were stained with $JF_{635}$-HSAm (1 nM). For DNA-PAINT, anti-vimentin immunostaining was used to label vimentin from the same cell-line with a DNA-oligonucleotide (docking strand). Complementary imager strand (P1-Atto655, 1 nM) was used for staining. Reconstructed super-resolution images with 31 nm (HT-PAINT) or 32 nm (DNA-PAINT) resolution. Scale bars: 1 µm. **B.** Exemplary relative frequency distribution recorded for $JF_{635}$-HSAm. A Gaussian function was fitted to the data to determine the mean bright times ($\tau_B$) in experiment performed as described in A. Analogously, the $\tau_B$ [ms] was obtained for additional xHTLs (mean ± standard deviation): SiR-HSAm = 715 ± 15, $JF_{635}$-HSAm = 365 ± 5, SiR-FSAm = 1125 ± 15. The inverse to $\tau_B$ yields the kinetic unbinding constant $k_{off}$ [$s^{-1}$]: $JF_{635}$-HSAm: 2.74, SiR-HSAm: 1.68, SiR-FSAm: 1.50.

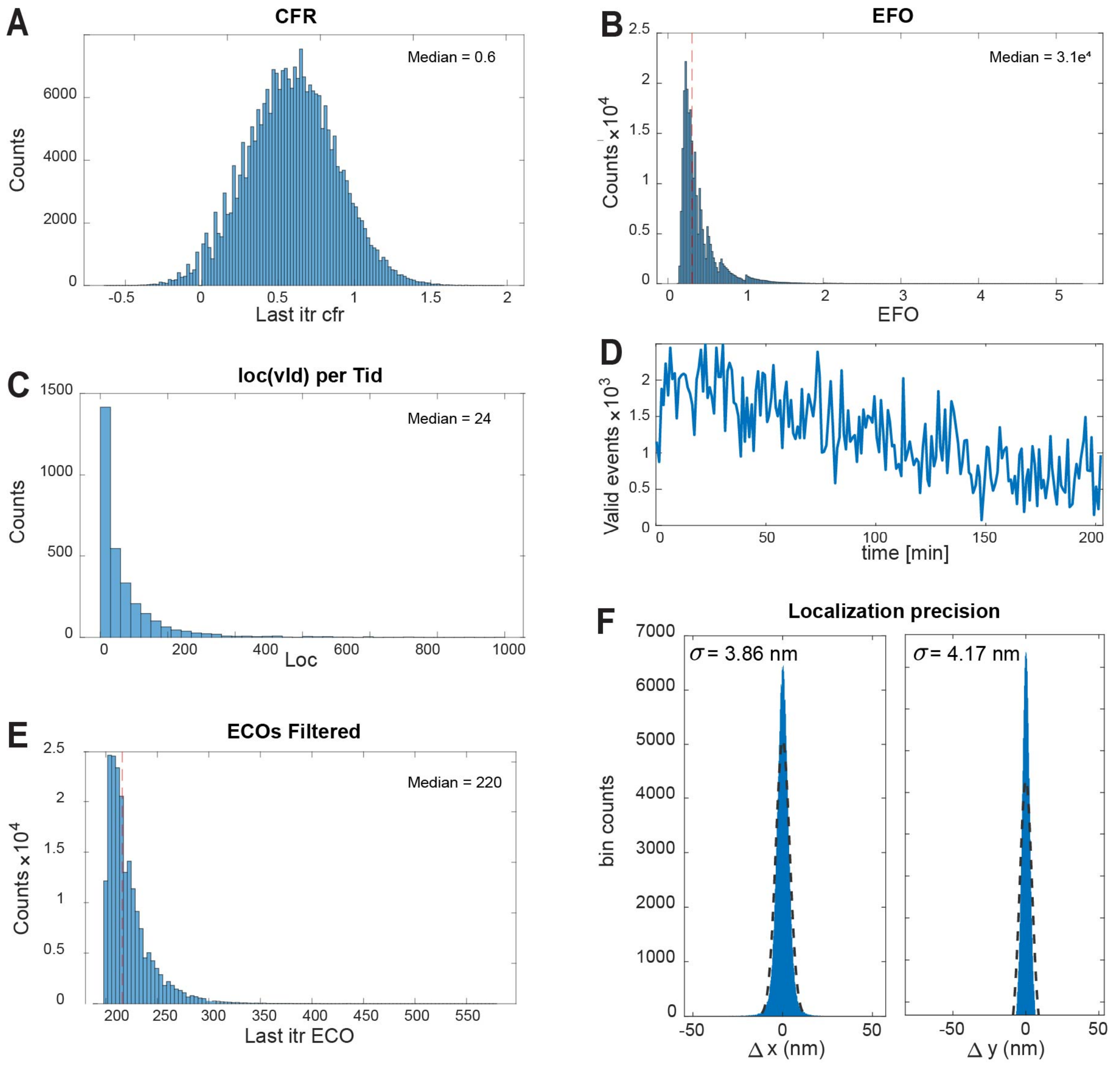

**Figure S. 9 | Characterization of xHTLs performance in MINFLUX microscopy.**

**A–D.** Raw data used for filtering of MINFLUX measurements shown in Fig 1F. Center frequency ratio (CFR) from last iteration (itr), median CFR = 0.6 (**A**). Effective frequency at offset (EFO), median EFO = 30 kHz (**B**). Number of valid (val) localized events (loc) per trace-iD (Tid), median loc/TiD = 34 (**C**). Valid event count over time (**D**). **E.** Effective counts at offset (ECO) from after filtering. 220 photons (median) were used in the last iteration step for localization. **F.** The localisation precision of MINFLUX measurements shown in Fig 1F was calculated to be 3.86 nm (x) and 4.17 nm (y), respectively, after filtering the data.

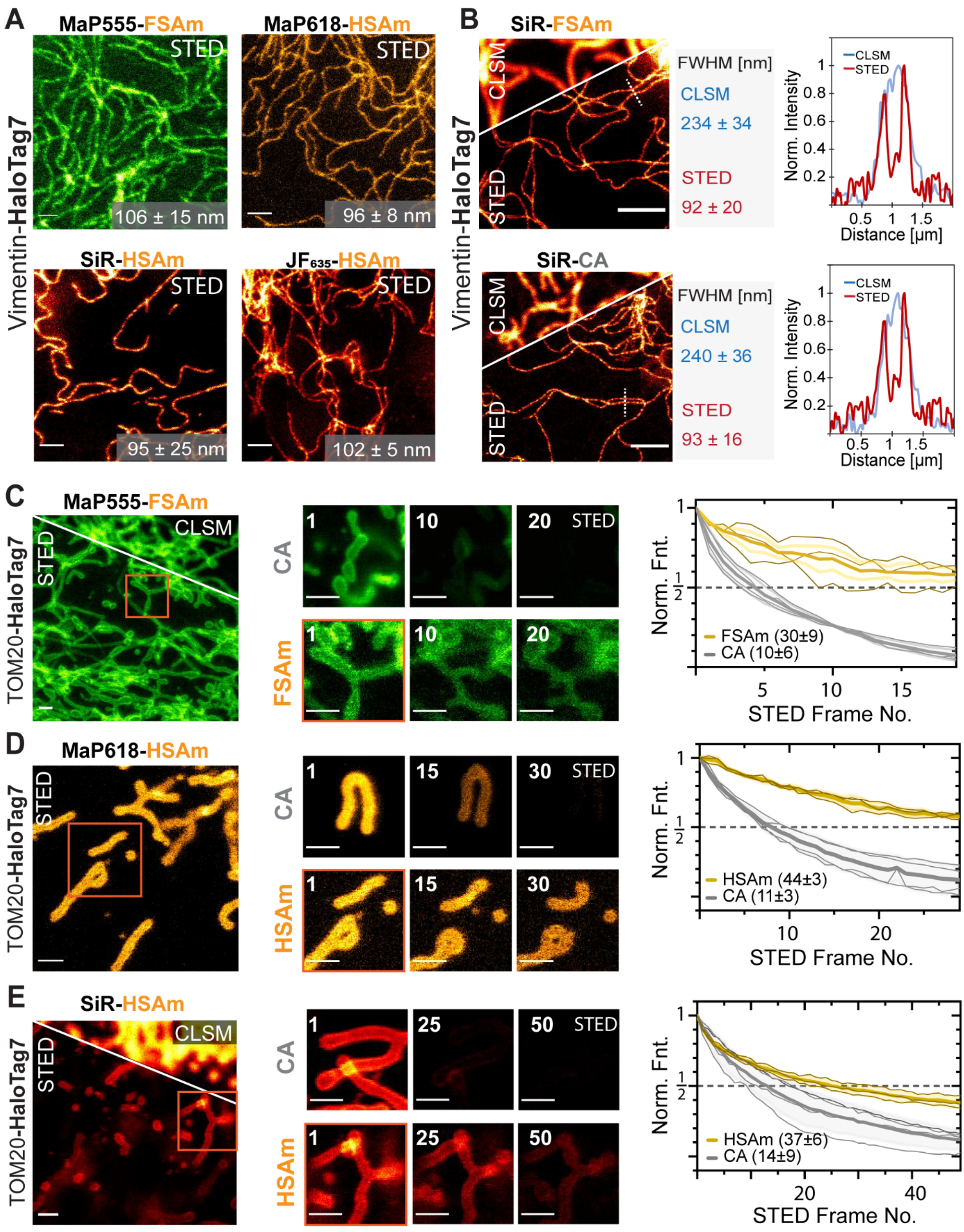

**Figure S. 10 | STED photobleaching resistance of xHTLs compared to covalent HTLs.**

**A.** Live-cell STED-images of U2OS cells endogenously expressing vimentin-HaloTag7[31] and stained using different xHTLs. Scale bar: 1 µm. The mean filament diameter was characterized by calculating the full with at half maximum (FWHM) from fluorescence intensity profiles perpendicular to vimentin filaments under STED microscopy conditions ($n > 15$ filaments from $\geq 2$ samples) as indicated in the bottom-right corner. **B.** Comparison of confocal laser scanning microscopy (CLSM) and STED images of live U2OS intermediate filaments stained with SiR-FSAm and SiR-CA. Same cell line as in A. Left side: confocal and STED images comparison. Middle: Comparison of the FWHM achieved under confocal and STED microscopy conditions ($n > 15$ filaments from $\geq 2$ samples). Right side: representative intensity profile along two adjunct intermediate filaments. **C-E.** Multi-frame STED imaging of U2OS mitochondria (TOM20-HaloTag7) stained with various xHTLs and HTLs. Left side: Overview STED image (8x8 µm) of cells expressing TOM20-HaloTag7 stained with different xHTLs (500 nM) and used for signal quantification during multi-frame STED imaging. Scale bar: 1 µm. Middle: Exemplary zoom on dynamic mitochondria during multi-frame STED-imaging comparing exchangeable xHTL and covalent HTL staining. Frame numbers indicated in the top left corner. Scale: 1 µm. Right side: Bleaching curves (thick lines: mean value and standard deviation, thin lines: individual experiments) and corresponding mean intensity half-life τ given in brackets ($n \geq 3$, from at least 2 individual experiments).

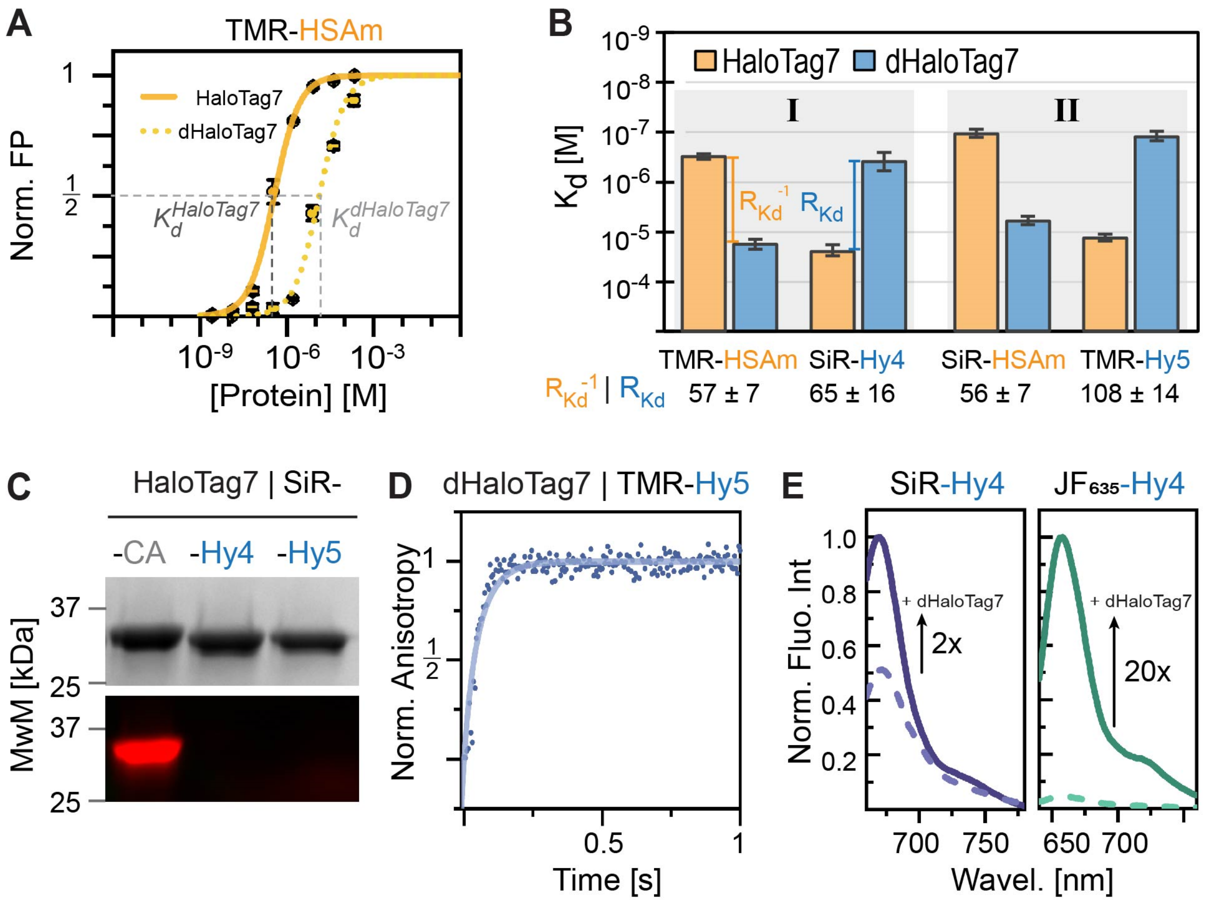

**Figure S. 11 | Development of an orthogonal exchangeable xHTL/HaloTag7 protein pair.**
**A.** Affinity titration of TMR-HSAm with HaloTag7 and dHaloTag7 measured by fluorescence polarization. **B.** Affinity comparison of spectrally distinct fluorescent xHTLs probes with HaloTag7 and dHaloTag7. $R_{Kd}^{-1}$ ($K_d^{HaloTag7}/K_d^{dHaloTag7}$) and $R_{Kd}(K_d^{dHaloTag7}/K_d^{HaloTag7})$ (see **Table S1**) were calculated as a measure for the binding specificity (mean values ± error calculated by Monte Carlo analysis). Using SiR and TMR fluorophores, xHTLs offer two spectrally distinct and orthogonal combinations: (I) TMR-HSAm/SiR-Hy4 and (II) SiR-HSAm/TMR-Hy5. **C.** HaloTag7 covalent labeling experiments. SDS-PAGE followed by in-gel fluorescence scan and Coomassie-staining. SiR derivatives of CA/Hy4/Hy5 ligands were incubated with HaloTag7. xHTLs show non-covalent binding to HaloTag7 in contrast to HTL covalent labeling (SiR-CA). MwM – Molecular weight marker. **D.** Binding kinetics of TMR-Hy5 to dHaloTag7 measured by stopped-flow fluorescence anisotropy experiments. Non-linear fitting from 10 kinetic traces yielded kinetic on-rate ($k_{on}$) **E.** Fluorescence emission spectra of free SiR/$JF_{635}$-Hy4 (dashed lines) or SiR/$JF_{635}$-Hy4 bound to dHaloTag7 (plain line). Arrows indicate fluorescence intensity increase ($F/F_0$) upon protein binding. SiR-Hy4 exhibits a low fluorescence increase upon protein binding which was restored by using the more fluorogenic dyes $JF_{635}$.

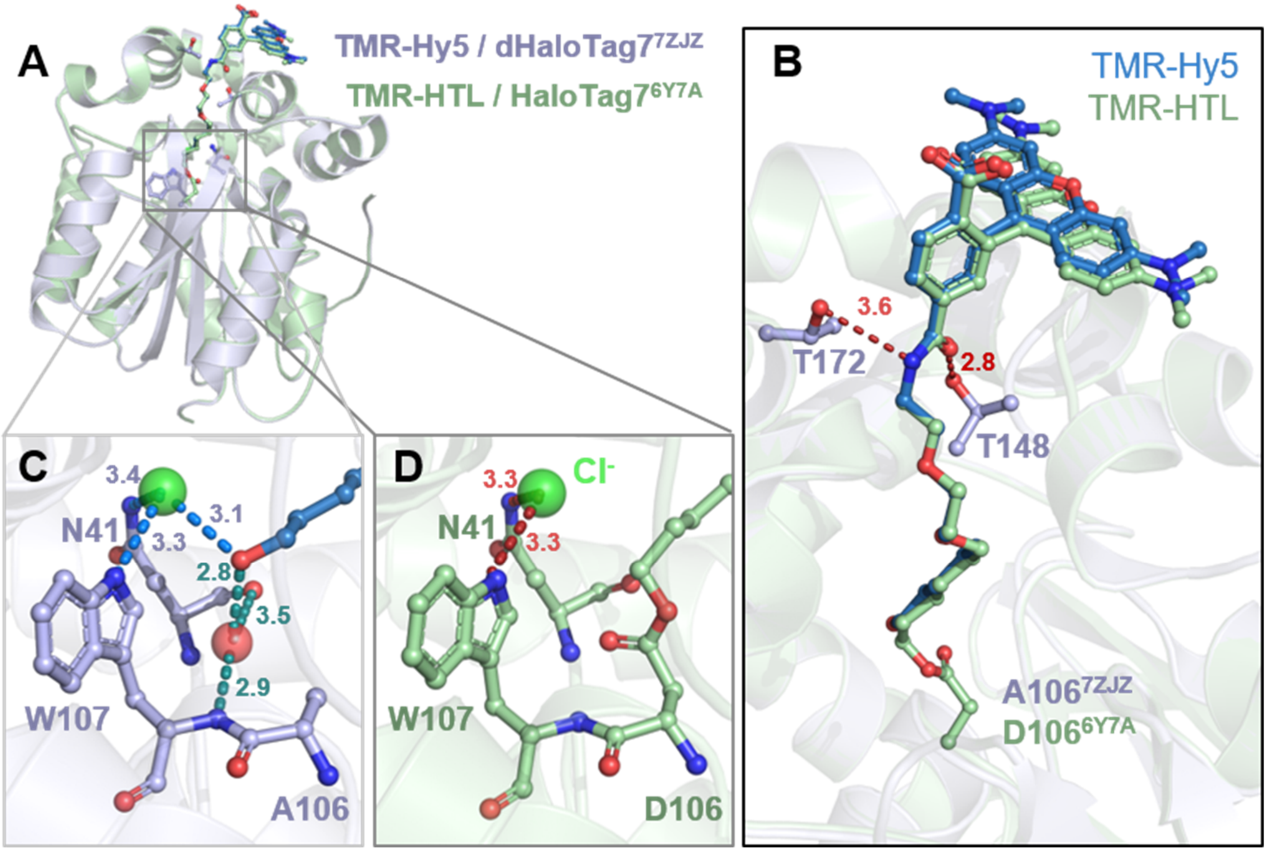

**Figure S. 12 | Structural analysis of the TMR-Hy5/dHaloTag7 complex.**
**A.** Structural comparison between the TMR-Hy5/dHaloTag7 complex (PDB-ID: 7ZJZ, 1.5 Å resolution) and TMR-HTL/HaloTag7 covalent complex (PDB-ID: 6Y7A, 1.4 Å resolution)[1]. Tertiary structures are represented as cartoons. The ligands and relevant residues are represented as sticks. **B.** Magnification on the TMR-ligands binding sites. Distances in Å. The TMR moieties are located at the protein's surface while the alkane chains are buried in the protein hydrophobic tunnel as previously described.[1] **C.** Magnification on the Hy5 binding site. Polar interactions between Hy5 and dHaloTag7 residues: TMR-Hy5 interacts with a structural water and chloride-ion molecule, presented as spheres. The chloride ions forms hydrogen bonds with W107 and N41 site chain and the water molecule occupies the space freed by the D106A mutation and interacts with the W107 and N41 main chain. **D.** Magnification on the covalent bond between TMR-HTL and the HaloTag7 protein. The TMR-labeled HaloTag7 features a chloride ion (green sphere) like found also in the TMR-Hy5/dHaloTag7 complex.

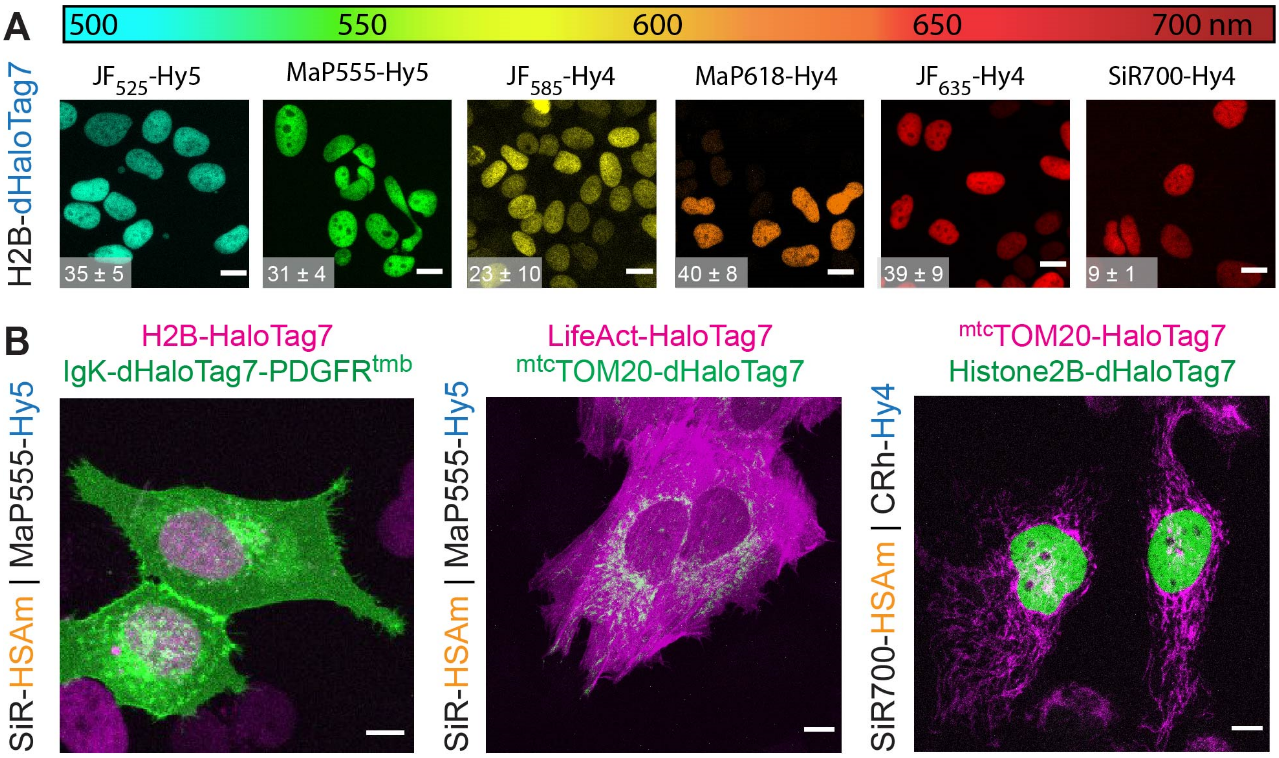

**Figure S. 13 | Dual-color confocal images using orthogonal xHTLs-(d)HaloTag7 pairs.**

**A.** Live-cell confocal images of different fluorescent xHTL probes covering the spectrum. H2B-dHaloTag7 expressing U2OS cells stained with 500 nM Hy5/Hy4 xHTLs probes and imaged via live-cell confocal microscopy. Sum projections. Scale: 10 µm. Signal-over-background ratios are given in bottom-left corner. **B.** Dual color images using orthogonal xHTLs-(d)HaloTag7 pairs at different subcellular localization of live U2OS cells. 500 nM of xHTLs. Localizations: Nucleus (H2B) and plasma membrane (IgK/PDGFR), actin (LifeAct) and mitochondria (TOM20). Scale: 10 µm. Images are all max. projection.

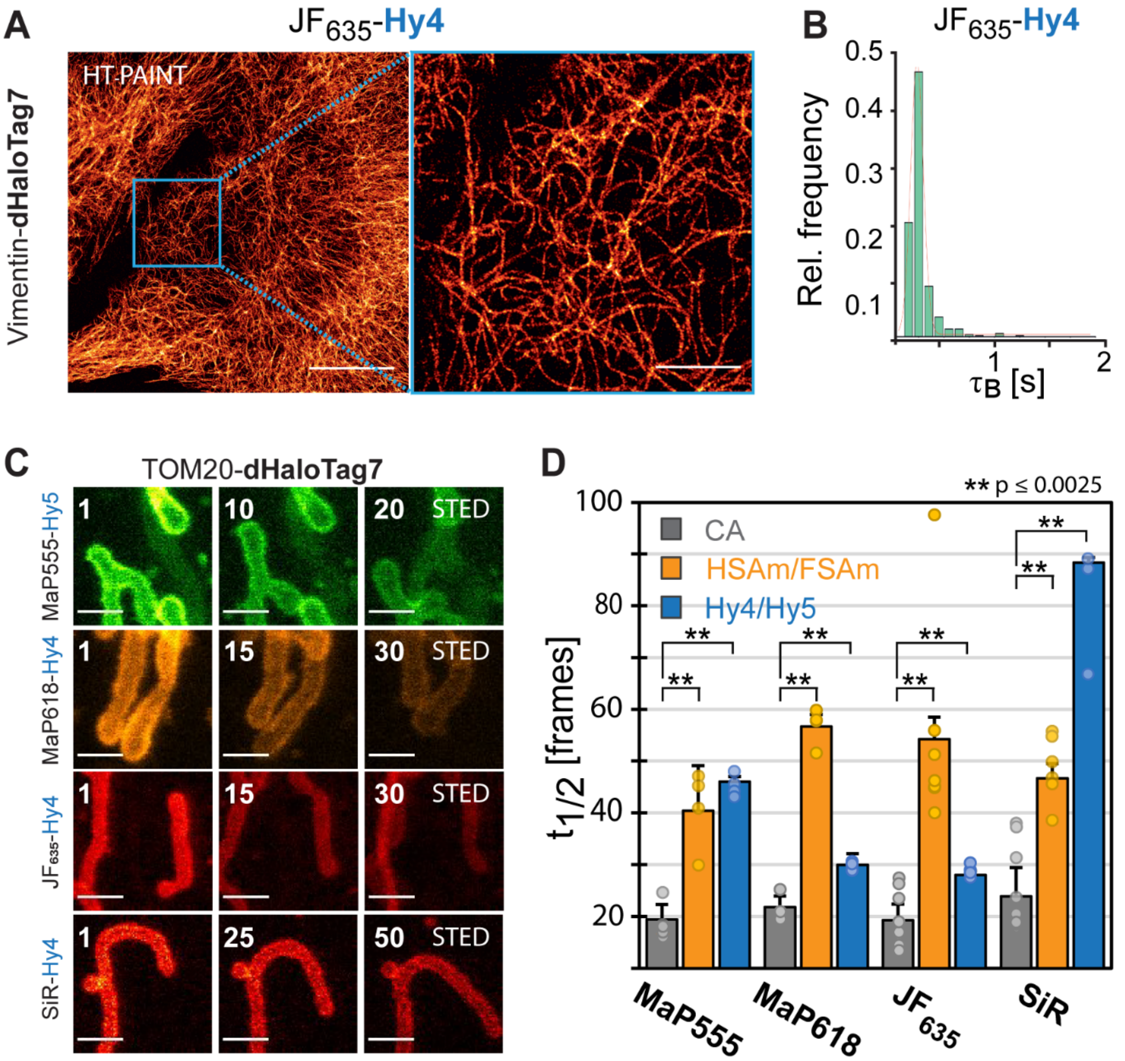

**Figure S. 14 | Super-resolution fluorescence microscopy enabled by Hy4/5 and dHaloTag7.**
**A.** Reconstructed super-resolution image of fixed U2OS cells' intermediate filaments obtained by HT-PAINT microscopy using xHTLs and dHaloTag7 (36 nm resolution). Cells overexpressing Vimentin-dHaloTag7 were stained with $JF_{635}$-Hy4 (3 nM). Scale bars: 10 µm (overview) or 2 µm (magnified region). **B.** Exemplary relative frequency distribution recorded for $JF_{635}$-Hy4. A Gaussian function was fitted to the data to determine the mean bright times ($\tau_B$) in experiment performed as described in A. Analogously, the $\tau_B$ [ms] parameter was obtained of additional xHTLs (mean ± S.D.): SiR-HSAm = 715 ± 15, $JF_{635}$-HSAm = 365 ± 5, SiR-FSAm = 1125 ± 15, SiR-Hy4: 282 ± 4 and $JF_{635}$-Hy4: 233 ± 2. The inverse to $\tau_B$ yields the kinetic unbinding constant $k_{off}$ [$s^{-1}$]: $JF_{635}$-Hy4: 4.35, SiR-Hy4: 3.4. **C.** Multi-frame STED images of U2OS mitochondria stained with xHTLs. Cells expressing TOM20-dHaloTag7 were stained with MaP555-Hy5 or MaP618-, $JF_{635}$-, SiR-Hy4 (500 nM). Frame numbers indicated in top-left corner. Scale bars: 10 µm. **D.** Comparison of STED bleaching experiments. Number of frames at which the intensity reaches half the initial one (half-life $\tau_{1/2}$) for various fluorophore-(x)HTLs. Normalized mean intensities were background corrected and fitted with a mono-exponential decay function. The half-life $\tau_{1/2}$ is represented. Individual data (dots), mean values (bars ± S.D., n≥3 experiments). Significance was calculated using two-sided t-tests including the Welch correction, not significant (n.s.): p > 0.05 (*), significant: p ≤ 0.0025 (**).

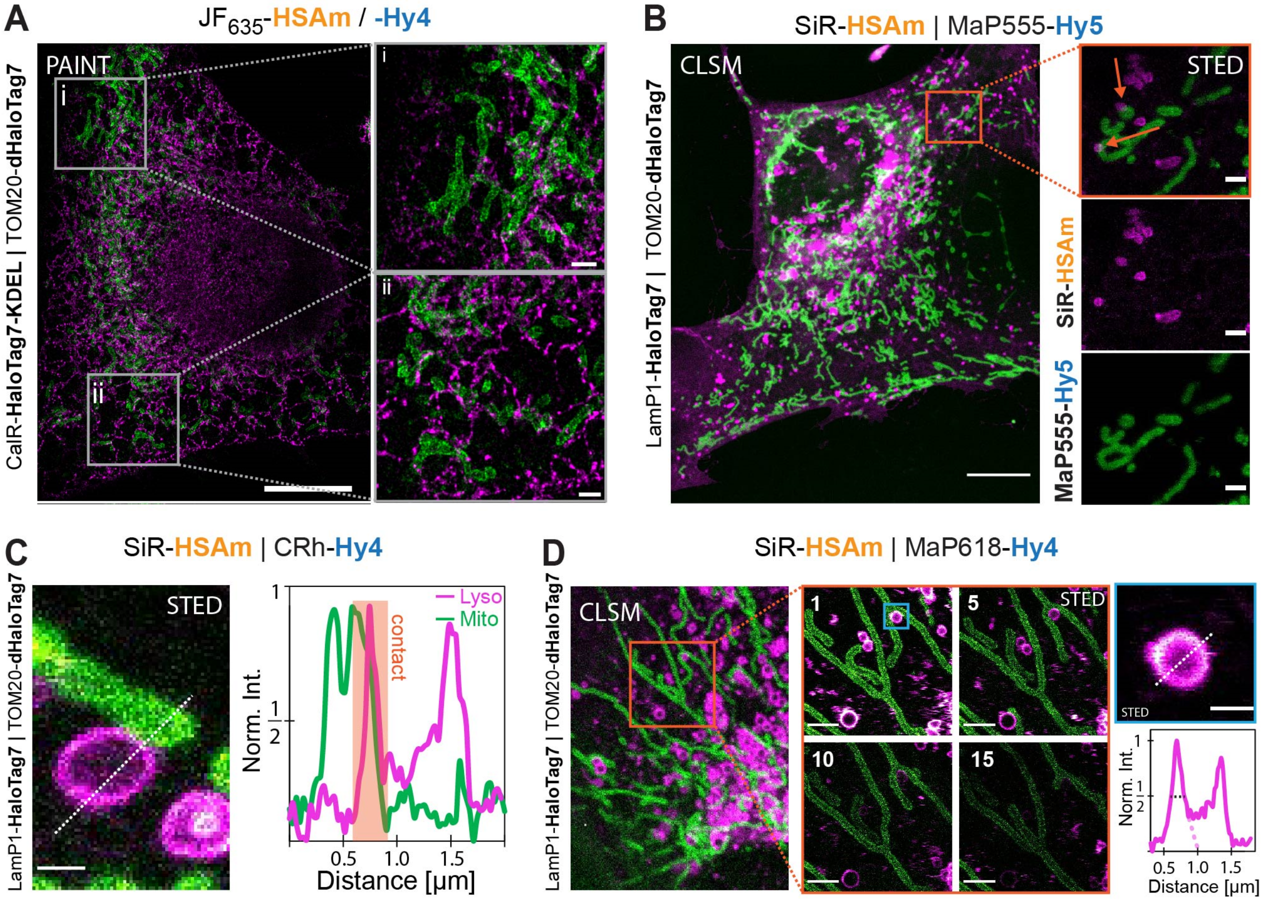

**Figure S. 15 | Dual-color super-resolution microscopy using xHTLs.**

**A.** Two-target super-resolution image of fixed U2OS cell's mitochondria and endoplasmic reticulum obtained by HT-PAINT microscopy using xHTLs. The cells were sequentially imaged with orthogonal label $JF_{635}$-HSAm (5 nM) and $JF_{635}$-Hy4 (3 nM) targeting the endoplasmic reticulum (CalR-HaloTag7-KDEL) and mitochondria (TOM20-dHaloTag7), respectively. White boxes indicate magnified regions. Scale bars: 10 µm (overview) and 2 µm (magnified region). **B.** Dual-color, live-cell confocal and STED image of U2OS cells' lysosome and mitochondria labeled with xHTLs (500 nM). Cells express TOM20-HaloTag7 and LamP1-dHaloTag7 through a T2A translational fusion and were labeled by SiR-HSAm (pink) and MaP555-Hy5 (green), respectively. Orange arrows indicate mitochondria-lysosome contact sites in the magnified region. Scale bars: 10 µm (overview) or 1 µm (magnified region). **C.** Dual-color STED image (magnification of image presented in Fig 2G) showing U2OS mitochondria-lysosome contacts (left panel). Cells were stained with SiR-HSAm (pink) and CRh-Hy4 (green, 500 nM each). Scale bar: 0.5 µm. Right panel: fluorescence intensity profile along the white dashed line along the mitochondria-lysosome contact site from the STED image. It highlights the hollow lysosome and its thin contact with mitochondria at sub-diffraction resolution. **D.** Multi-frame STED imaging of U2OS cells expressing LamP1-HaloTag7 and TOM20-dHaloTag7 stained with SiR-HSAm (pink) and MaP618-Hy4 (green, 500 nM each) respectively. Scale bar: 0.5 µm. The fluorescence profile along the white dashed line of the top right lysosome STED image highlights the hollow lysosome structure. The mean vesicular membrane diameter FWHM (black dashed line) under STED microscopy conditions was calculated to be 195 ± 85 (n = 15 vesicles).

### Supplementary Tables

| | Fluo-xHTL | | Spectral prop. | | | | Binding properties | | | | Applications | | | | | | | | | |
|---|---|---|---|---|---|---|---|---|---|---|---|---|---|---|---|---|---|---|---|---|
| | xHTL | Dye | $\lambda_{ext}$ | $\lambda_{em}$ | ΦHT | ± S.D. | $K_d$ | (CI 95%) | $k_1$ | ± S.D. | $F/F_0$ | ± S.D. | $R_{Kd}^{-1}$ | ± S.E. | Microscopy | | | | Best comb. w. | |
| | | | [nm] | [nm] | % | | [nM] | | $[10^6 M^{-1} s^{-1}]$ | | buff/HT | | HT7/dHT7 | | Conf.[live] | STED[live] | PAINT | MINFLUX | $\lambda_{ext}$ = 550 nm | $\lambda_{ext}$ = 610 - 650 nm |
| **HT7 ligands** | **HSAm** | TMR | 555 | 578 | 61.3 | ± 0.4 | 310 | (275 - 351) | 6.4 | ± 0.1 | 1.3 | ± 0.1 | 57 | ± 7 | ☐ | ☐ | ☐ | ☐ | | |
| | | CRh | 616 | 636 | 72.2 | ± 0.5 | 142 | (115 - 176) | 2.5 | ± 0.3 | 1.6 | ± 0.3 | N.D. | | ☑ | ☑ | ☐ | ☐ | | CRh-Hy4 |
| | | MaP618 | 616 | 636 | 67.4 | ± 0.1 | N.D. | | N.D. | | 29.9 | ± 0.3 | N.D. | | ☑ | ☑ | ☐ | ☐ | **MaP555-Hy5** | MaP618-Hy4 |
| | | $JF_{635}$ | 635 | 660 | 59.6 | ± 0.9 | 117 | (84 - 162) | 7.8 | ± 0.4 | 8.7 | ± 2.0 | N.D. | | ☑ | ☑ | ☑ | ☑ | | SiR-Hy4 |
| | | SiR | 646 | 672 | 61.3 | ± 0.4 | 109 | (91 - 130) | 4.6 | ± 0.1 | 2.5 | ± 0.7 | 56 | ± 7 | ☑ | ☑ | ☑ | ☑ | | **$JF_{635}$-Hy4** |
| | **FSAm** | TMR | 555 | 578 | 60.2 | ± 0.1 | 166 | (150 - 185) | 6.8 | ± 0.1 | 1.4 | ± 0.1 | 10 | ± 2 | ☐ | ☐ | ☐ | ☐ | | |
| | | MaP555 | 558 | 578 | 45.3 | ± 0.6 | 167 | (63 - 70) | 6.3 | ± 0.2 | 5.8 | ± 1.4 | N.D. | | ☐ | ☑ | ☐ | ☐ | / | / |
| | | CRh | 616 | 636 | 68.8 | ± 0.5 | 60 | (48 - 75) | 5.0 | ± 0.8 | 2.3 | ± 0.3 | N.D. | | ☑ | ☑ | ☐ | ☐ | | |
| | | SiR | 646 | 672 | 60.2 | ± 0.1 | 67 | (48 - 93) | 7.3 | ± 0.3 | 10 | ± 3.9 | 4 | ± 1 | ☑ | ☑ | ☑ | ☑ | | |
| | | | $\lambda_{ext}$ | $\lambda_{em}$ | ΦdHT | ± S.D. | $K_d$ | (C I95%) | $k_1$ | ± S.D. | $F/F_0$ | ± S.D. | $R_{Kd}$ | ± S.E. | Conf.[live] | STED[live] | PAINT | MINFLUX | Best comb. w. | |
| **dHT7 ligands** | **Hy4** | TMR | 555 | 578 | 57.9 | ± 0.4 | 1004 | (836 - 1204) | 11 | ± 0.3 | 1.1 | ± 0.2 | 113 | ± 21 | ☐ | ☐ | ☐ | ☐ | | |
| | | CRh | 618 | 636 | 71.1 | ± 0.5 | 219 | (150 - 382) | 15 | ± 0.4 | 1.5 | ± 0.3 | N.D. | | ☑ | ☑ | ☐ | ☐ | | |
| | | MaP618 | 618 | 636 | 68.3 | ± 0.2 | N.D. | | N.D. | | 37 | ± 6.5 | N.D. | | ☑ | ☑ | ☐ | ☐ | | CRh-HSAm |
| | | $JF_{635}$ | 635 | 660 | 68.2 | ± 0.3 | 272 | (170 - 436) | 26 | ± 1.8 | 19 | ± 2.7 | N.D. | | ☑ | ☑ | ☑ | ☐ | | MaP618-HSAm |
| | | SiR | 646 | 672 | 60.7 | ± 0.4 | 385 | (253 - 589) | 12 | ± 0.4 | 1.3 | ± 0.8 | 108 | ± 14 | ☑ | ☑ | ☑ | ☐ | / | SiR-HSAm |
| | **Hy5** | TMR | 555 | 578 | 60.9 | ± 0.4 | 125 | (101 - 154) | 10 | ± 0.2 | 1.4 | ± 0.2 | 65 | ±16 | ☑ | ☐ | ☐ | ☐ | | **$JF_{635}$-HSAm** |
| | | MaP555 | 558 | 578 | 53.8 | ± 1.2 | 186 | (123 - 189) | 14 | ± 0.4 | 1.4 | ± 0.2 | N.D. | | ☑ | ☐ | ☐ | ☐ | | |
| | | SiR | 646 | 672 | 60.9 | ± 0.4 | 86 | (74 - 100) | 10 | ± 0.3 | 1.9 | ± 0.2 | 51 | ± 7 | ☑ | ☑ | ☑ | ☐ | | |

ΦHT / ΦdHT – Quantum Yield bound to (d)HaloTag7, CI 95% - 95% confidence interval, $F/F_0$ – Fluorescence intensity increase upon (d)HaloTag7 binding, $R_{Kd}$ – Ratio $K_d$(dHaloTag7)/$K_d$(HaloTag7) S.D. – standard deviation, S.E. – standard error. N.D. – Not determined.

**Table S. 1 | Data collection and refinement statistics for the crystal structure.**

| **Data collection** | **HaloTag7 HSAm-TMR (7ZJ0)** | **HaloTag7 FSAm-TMR (7ZIY)** | **dHaloTag7 Hy5-TMR (7ZIZ)** |
|---|---|---|---|
| Space group | P1 | P1 | $P2_12_12$ |
| Unit-cell parameters | | | |
| a, b, c (Å) | 44.29, 49.93, 78.72 | 44.24, 46.06, 79.06 | 77.83, 88.71, 44.20 |
| $\alpha$, $\beta$, $\gamma$ (°) | 71.23, 89.91, 67.81 | 94.41, 90.00, 109.51 | 90.00, 90.00, 90.00 |
| Radiation source | PXII-X10SA, SLS | PXII-X10SA, SLS | PXII-X10SA, SLS |
| Wavelength (Å) | 0.99988 | 0.99996 | 1.00008 |
| Temperature (K) | 100 | 100 | 100 |
| Resolution range (Å) | 50-1.50 (1.60-1.50) | 50-1.70 (1.80-1.70) | 50-1.50 (1.60-1.50) |
| No. of observed reflections | 165811 (29286) | 108087 (15170) | 347669 (62060) |
| No. of unique reflections | 85890 (14710) | 59538 (8598) | 49719 (8612) |
| Multiplicity | 1.9 (2.0) | 1.8 (1.8) | 7.0 (7.2) |
| Completeness (%) | 91.4 (88.6) | 92.3 (84.7) | 99.8 (99.6) |
| $R_{merge}$ (%) | 3.9 (43.2) | 3.6 (23.4) | 6.2 (34.4) |
| $<I/\sigma(I)>$ | 11.8 (2.5) | 12.3 (3.0) | 18.7 (6.2) |
| $CC_{1/2}$ (%)[#] | 99.7 (77.0) | 99.8 (90.6) | 99.8 (97.1) |
| **Refinement** | | | |
| Molecules per a.u. | 2 | 2 | 1 |
| No. of reflections | 85886 | 59535 | 49714 |
| No. of reflections in test set | 4295 | 2977 | 2486 |
| Resolution range (Å) | 43.36-1.50 | 39.40-1.70 | 44.35-1.50 |
| No. of non-hydrogen atoms | | | |
| Protein | 4728 | 4714 | 2360 |
| Ligand/ion | 132 | 104 | 51 |
| Water | 417 | 271 | 233 |
| Total | 5277 | 5089 | 2644 |
| R (%) | 17.47 | 19.10 | 16.92 |
| $R_{free}$ (%) | 20.52 | 22.43 | 19.48 |
| RMS deviations from ideal | | | |
| bonds (Å) | 0.013 | 0.012 | 0.011 |
| angles (°) | 1.264 | 1.226 | 1.188 |
| B-factors (Å$^2$) | | | |
| Protein | 19.26 | 20.32 | 16.69 |
| Ligand/ion | 22.25 | 21.03 | 18.41 |
| Water | 26.61 | 26.23 | 25.79 |
| Average | 19.92 | 20.65 | 17.52 |
| Wilson B ( Å$^2$) | 16.69 | 19.22 | 15.44 |
| Ramachandran statistics (%) | | | |
| favored regions | 96.0 | 95.4 | 96.9 |
| allowed regions | 4.0 | 4.6 | 3.1 |
| disallowed regions | 0 | 0 | 0 |
| Clashscore | 1.78 | 1.70 | 1.06 |

Values in parentheses are for the highest resolution shell; [#]as implemented in XDS[21].

**Table S.2 | Fluorescence intensity increase of fluorogenic xHTL probes upon target binding.**

| Dye | HSAm | FSAm | Hy4 | Hy5 |
|---|---|---|---|---|
| $JF_{525}$ | 1.1 ± 1.6 | 5.2 ± 0.2 | N.D. | 1.4 ± 0.4 |
| TMR | 1.3 ± 0.1 | 1.4 ± 0.1 | 1.1 ± 0.2 | 1.4 ± 0.2 |
| MaP555 | 1.5 ± 0.4 | 5.8 ± 1.4 | N.D. | 1.4 ± 0.2 |
| CRh | 1.6 ± 0.3 | 2.3 ± 0.3 | N.D. | N.D. |
| MaP618 | 29.9 ± 0.3 | N.D. | 37.1 ± 6.5 | N.D. |
| $JF_{635}$ | 8.7 ± 2.0 | N.D. | 18.9 ± 2.7 | N.D. |
| SiR | 2.5 ± 0.7 | 10.2 ± 3.9 | 1.3 ± 0.8 | 1.9 ± 0.2 |
| SiR700 | 2.8 ± 0.6 | N.D. | 2.9 ± 0.3 | N.D. |

HSAm/FSAm binding to HaloTag7 and Hy4/Hy5 binding to dHaloTag7. N.D. – not determined.

**Table S.4 | Single-molecule binding kinetics and image resolution in HT-PAINT**

| Dye | Ligand | $\tau_b$ [ms] ± S.D. | $k_{off}$ [$s^{-1}$] | res [nm] |
|---|---|---|---|---|
| SiR | HSAm | 715 ± 15 | 1.7 | 34 |
| $JF_{635}$ | HSAm | 365 ± 5 | 2.7 | 31 |
| SiR | FSAm | 1125 ± 15 | 1.4 | 32 |
| SiR | Hy4 | 282 ± 4 | 3.4 | 37 |
| $JF_{635}$ | Hy4 | 233 ± 2 | 4.4 | 36 |
| Atto655 | P1 (DNA 9mer) | 620* | 1.6* | 32 |

* Taken from Jungmann *et al.* (2010)[43], S.D. – standard deviation

### Protein sequences

>His-TEV-HaloTag7 (and variant)

MHHHHHHHHHHENLYFQGIGTGFPFDPHYVEVLGERMHYVDVGPRDGTPVLFLHGNPTSSYVWR
NIIPHVAPTHRCIAPDLIGMGKSDKPDLGYFFDDHVRFMDAFIEALGLEEVVLVIHXWGSALGFHWAK
RNPERVKGIAFMEFIRPIPTWDEWPEFARETFQAFRTTDVGRKLIIDQNVFIEGTLPMGVVRPLTEVE
MDHYREPFLNPVDREPLWRFPNELPIAGEPANIVALVEEYMDWLHQSPVPKLLFWGTPGVLIPPAE
AARLAKSLPNCKAVDIGPGLNLLQEDNPDLIGSEIARWLSTLEI

Green: His-tag, Purple: TEV-cleavage site, Blue: HaloTag7, X = D for HaloTag7 & X = A for dHaloTag7.

#### NMR spectra

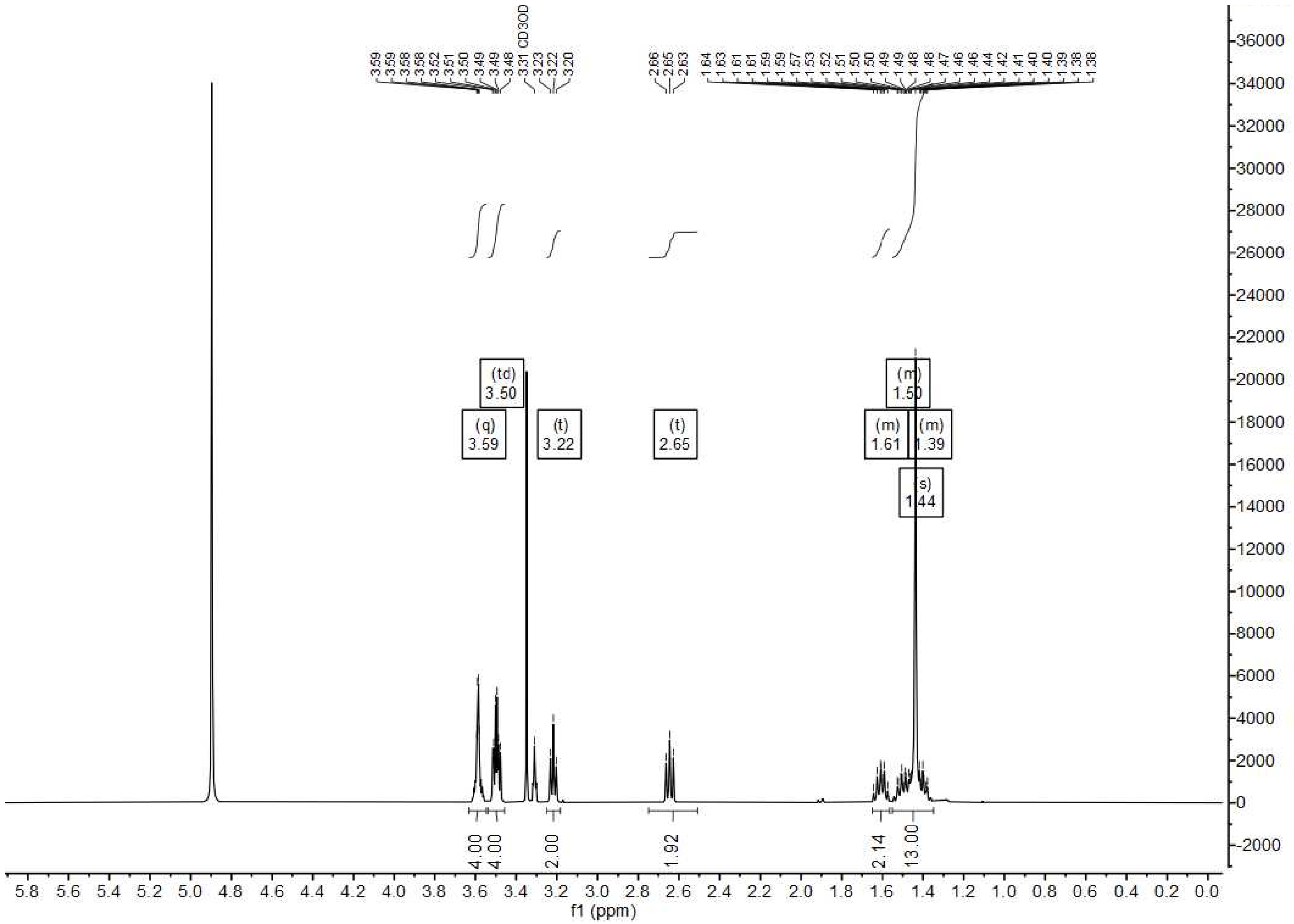

**Spectrum 1.** $^{1}$H-NMR (MeOD) of Boc-NH-PEG$_2$-C$_5$-NH$_2$ (**6**).

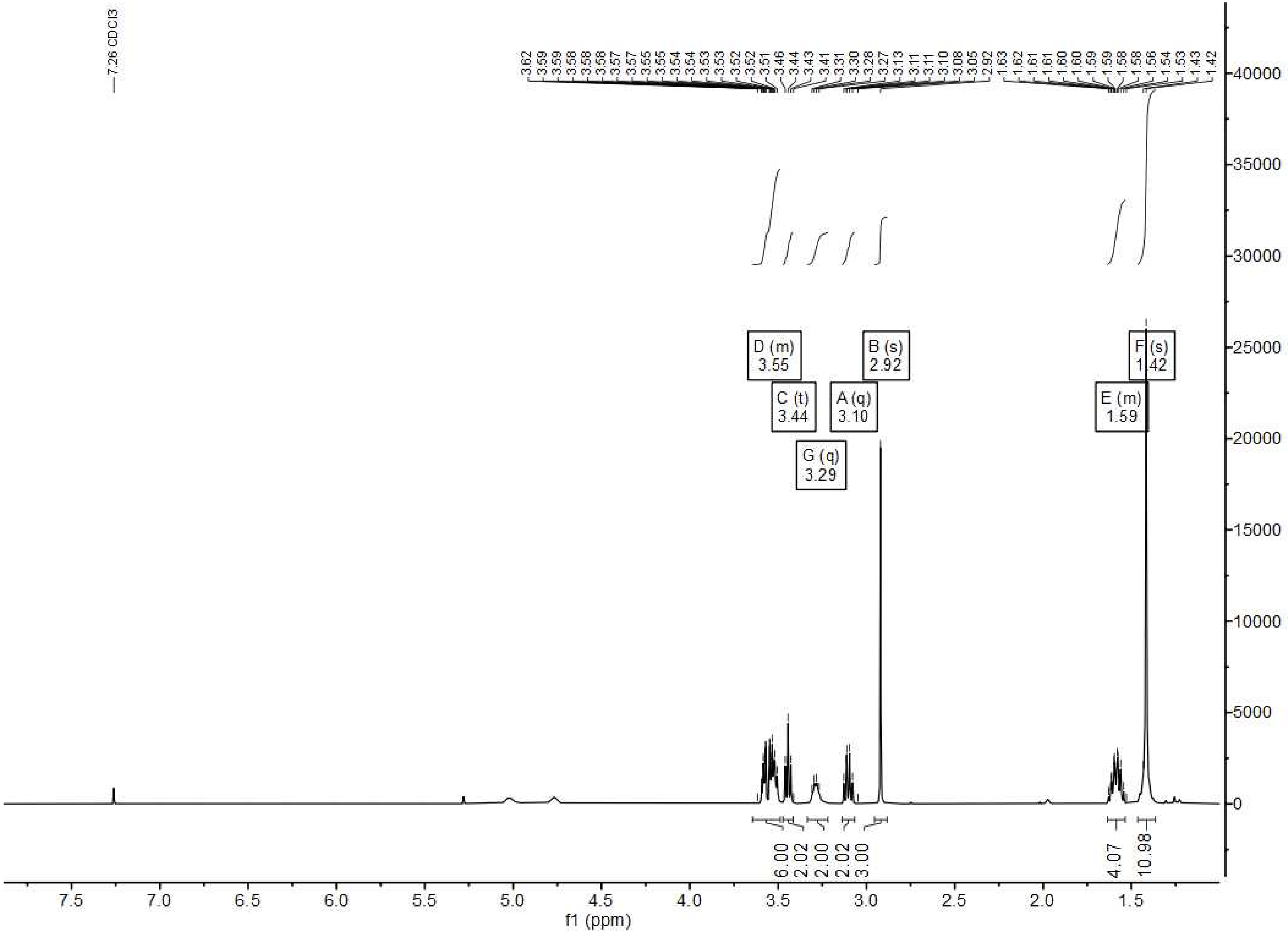

**Spectrum 2.** $^1$H-NMR ($CDCl_3$) of Boc-HSAm (**7**).

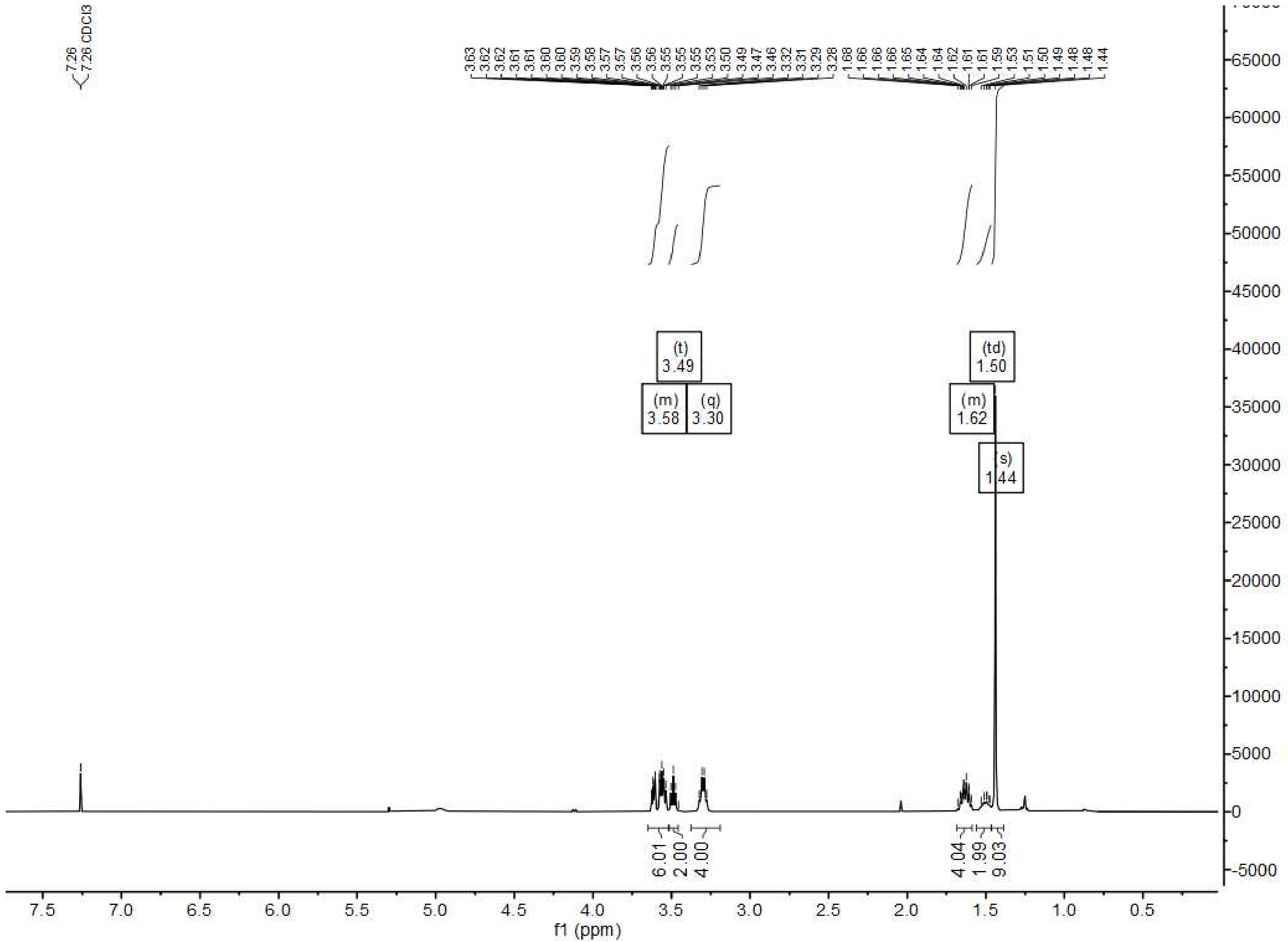

**Spectrum 3**. $^{1}$H-NMR ($CDCl_3$) of Boc-FSAm (**8**).

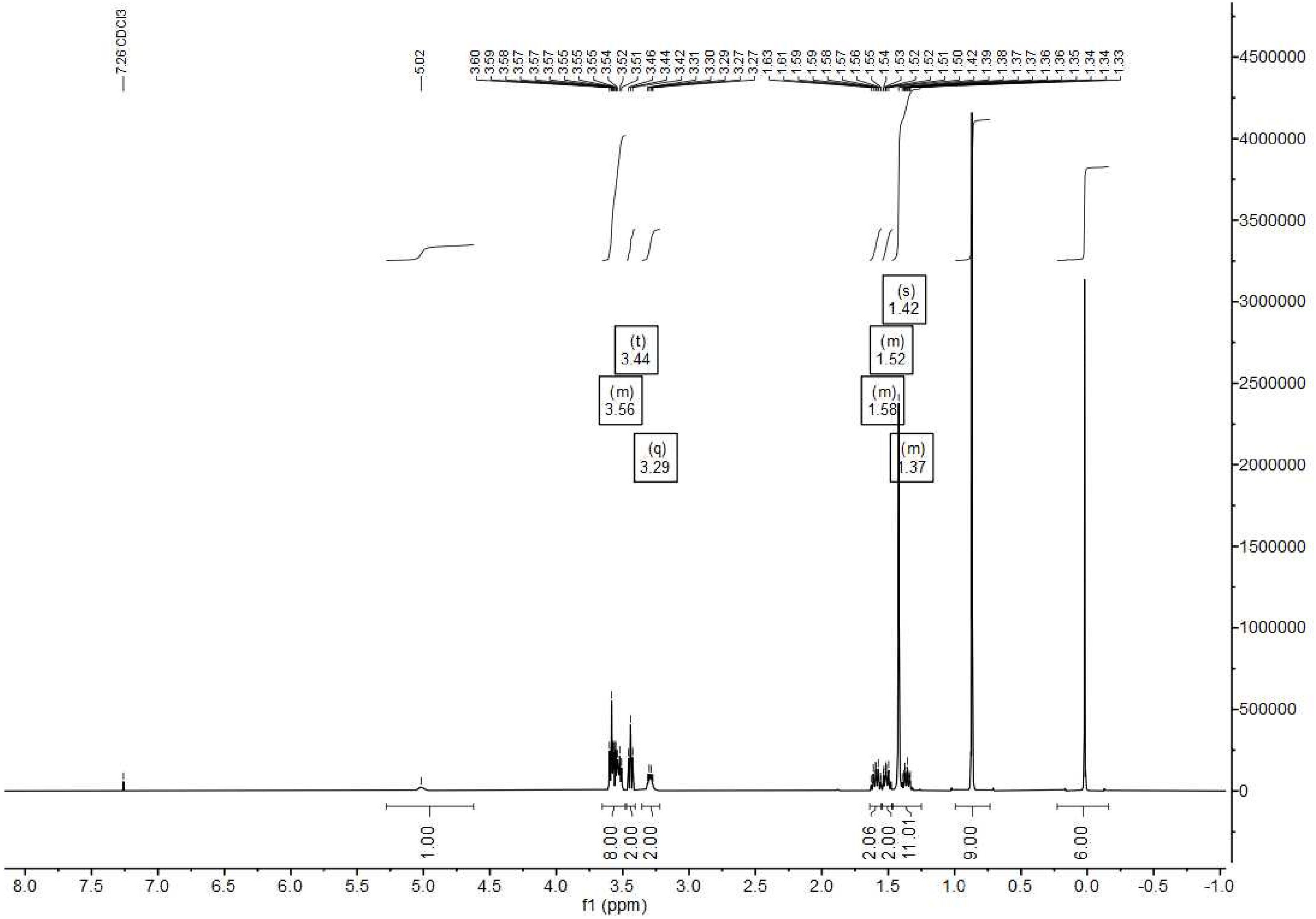

**Spectrum 4.** $^1$H-NMR ($CDCl_3$) of Boc-Hy5-TBS (**11**).

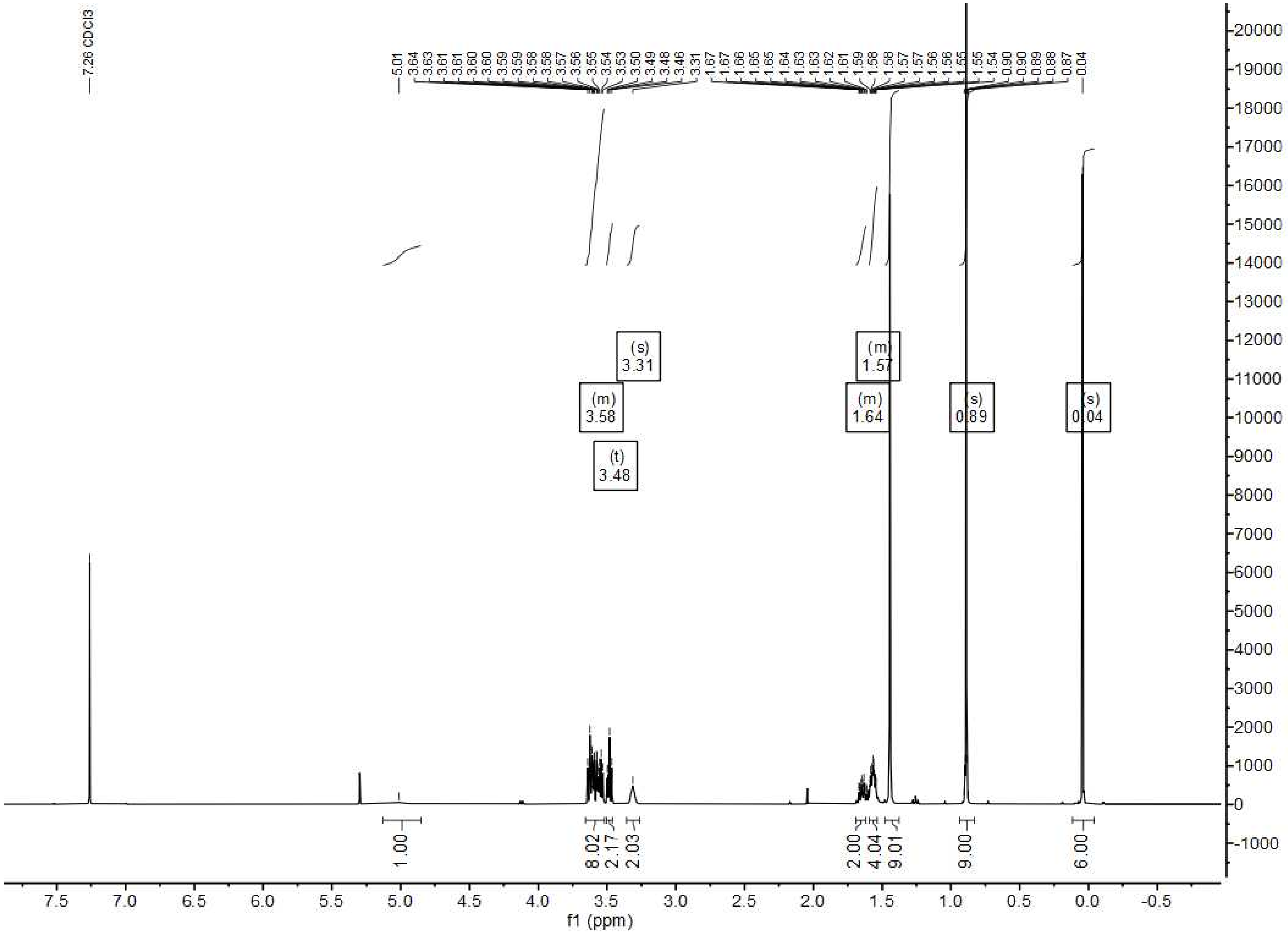

**Spectrum 5.** $^{1}$H-NMR ($CDCl_3$) of Boc-Hy4-TBS (**13**).

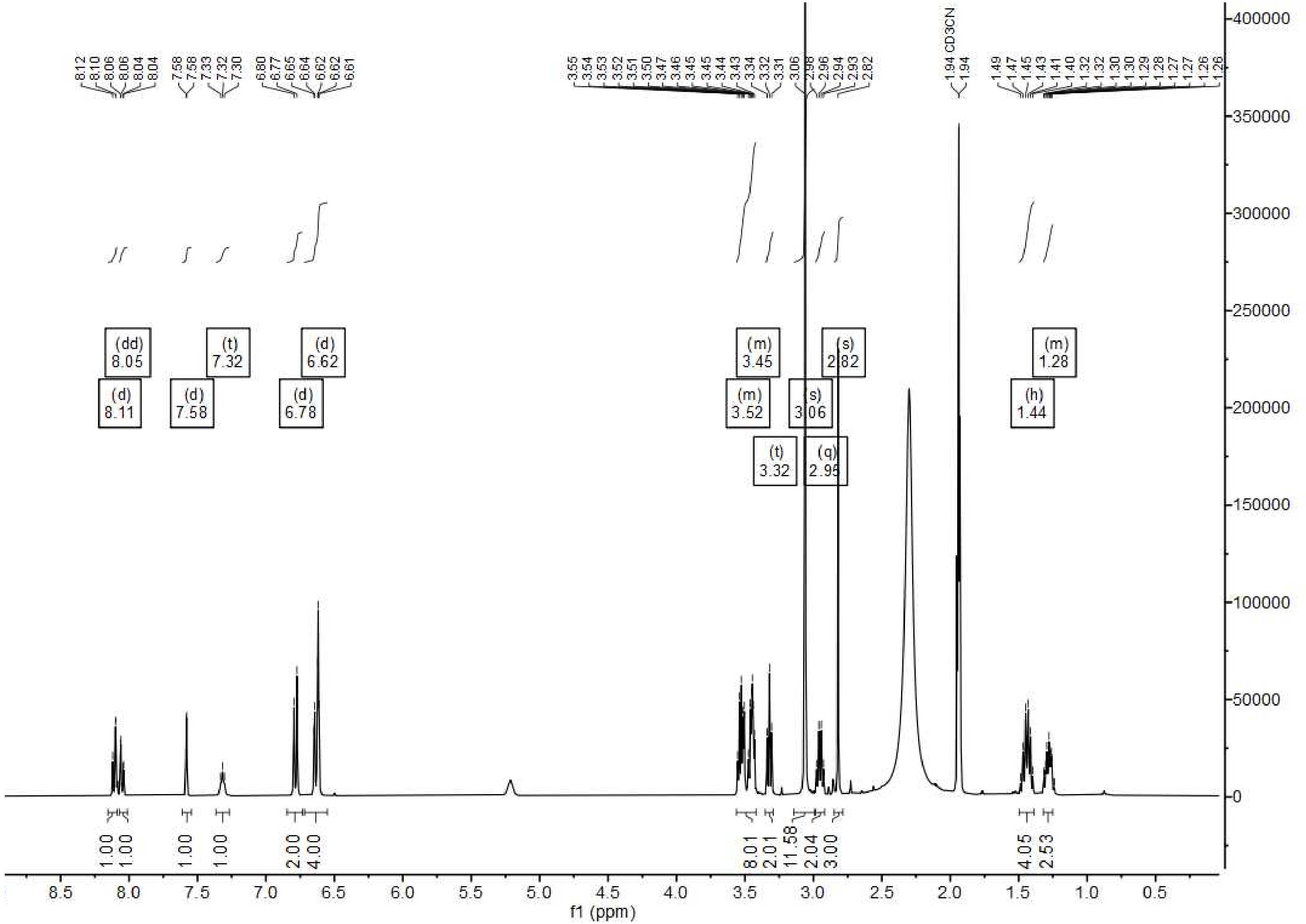

**Spectrum 6**. ${}^{1}$H-NMR ($CD_3CN$) of TMR-HSAm (**14**).

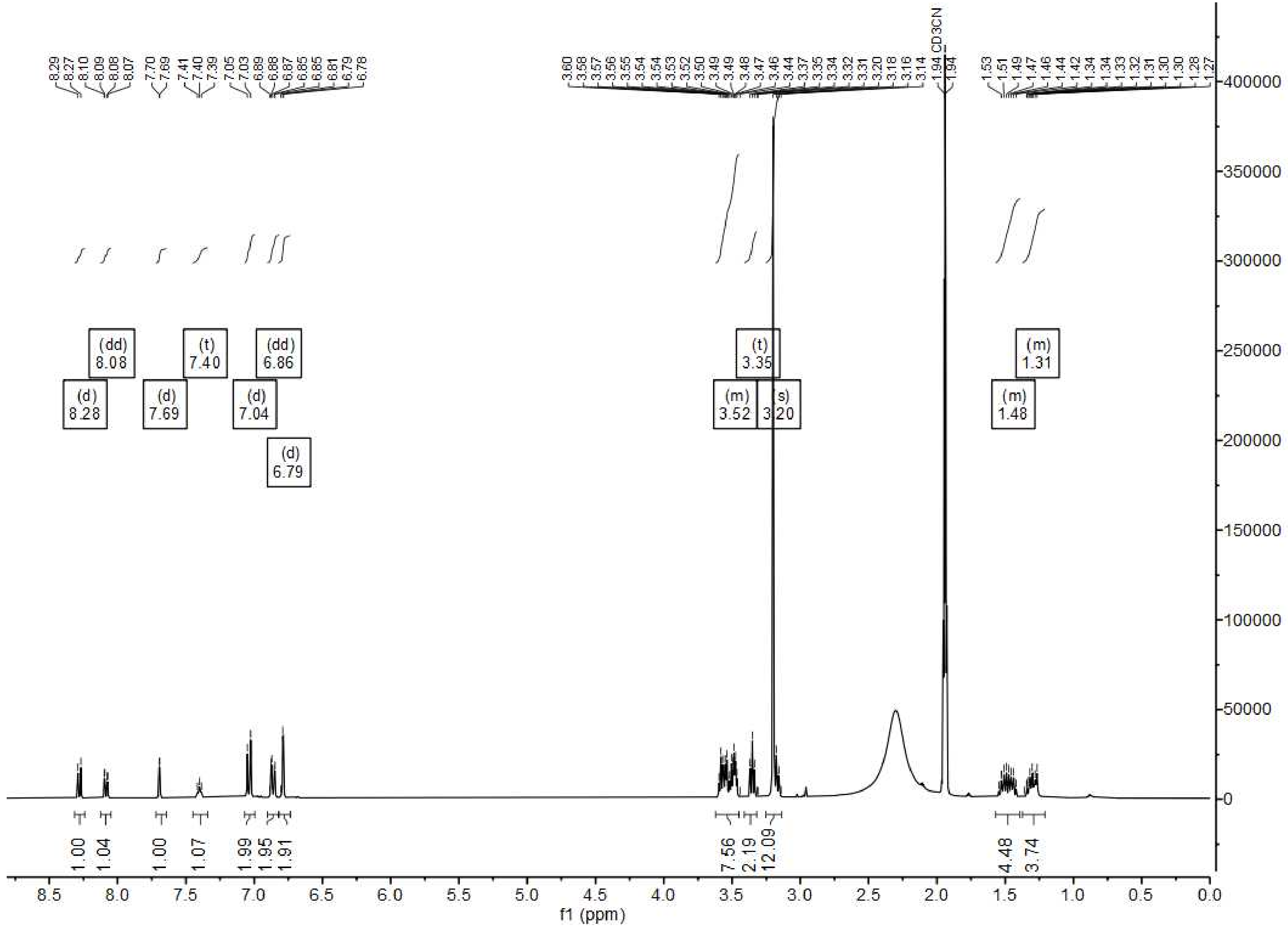

**Spectrum 7.** $^1$H-NMR ($CD_3CN$) of TMR-FSAm (**15**).

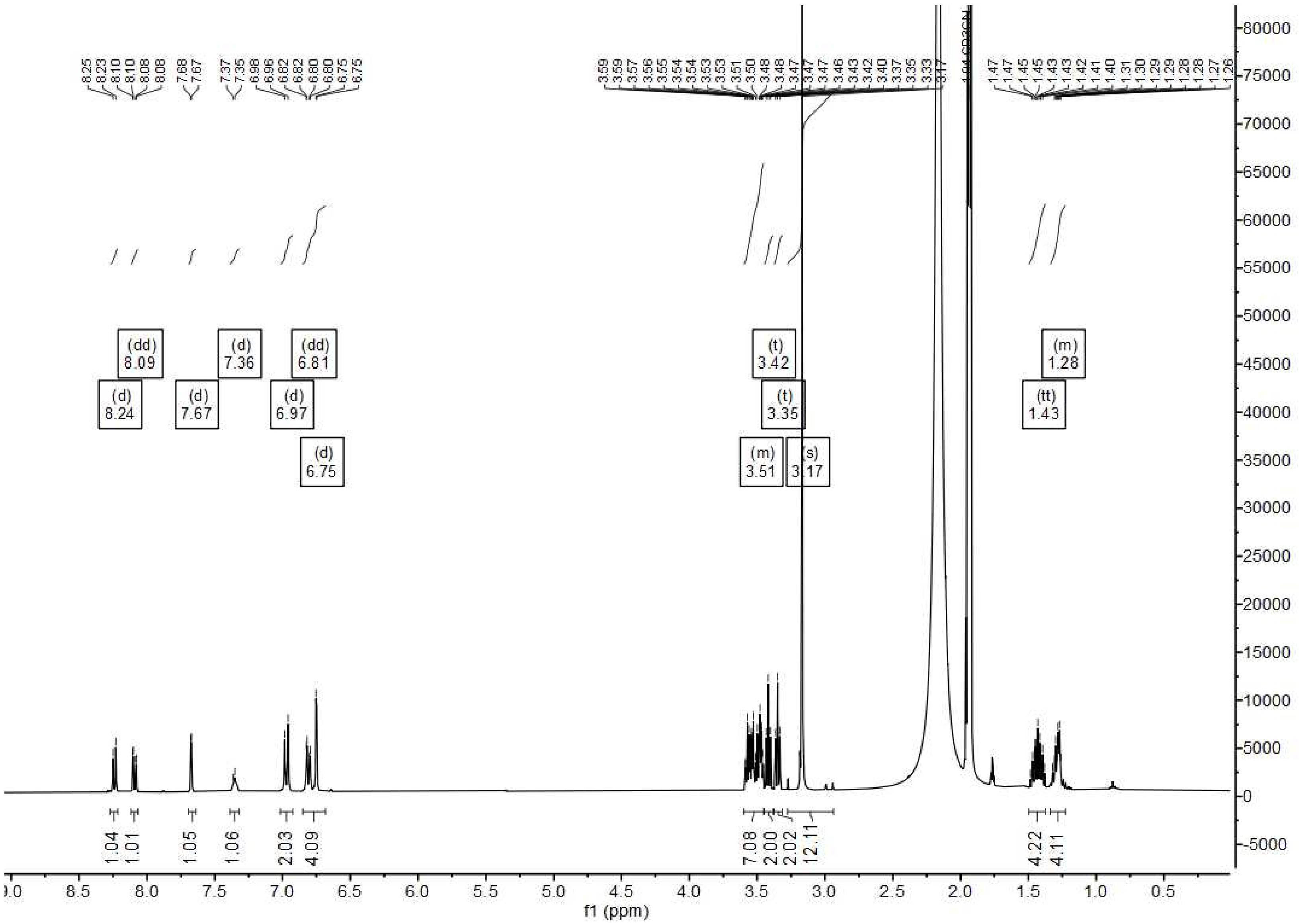

**Spectrum 8.** $^1$H-NMR ($CD_3CN$) of TMR-Hy5 (**16**).

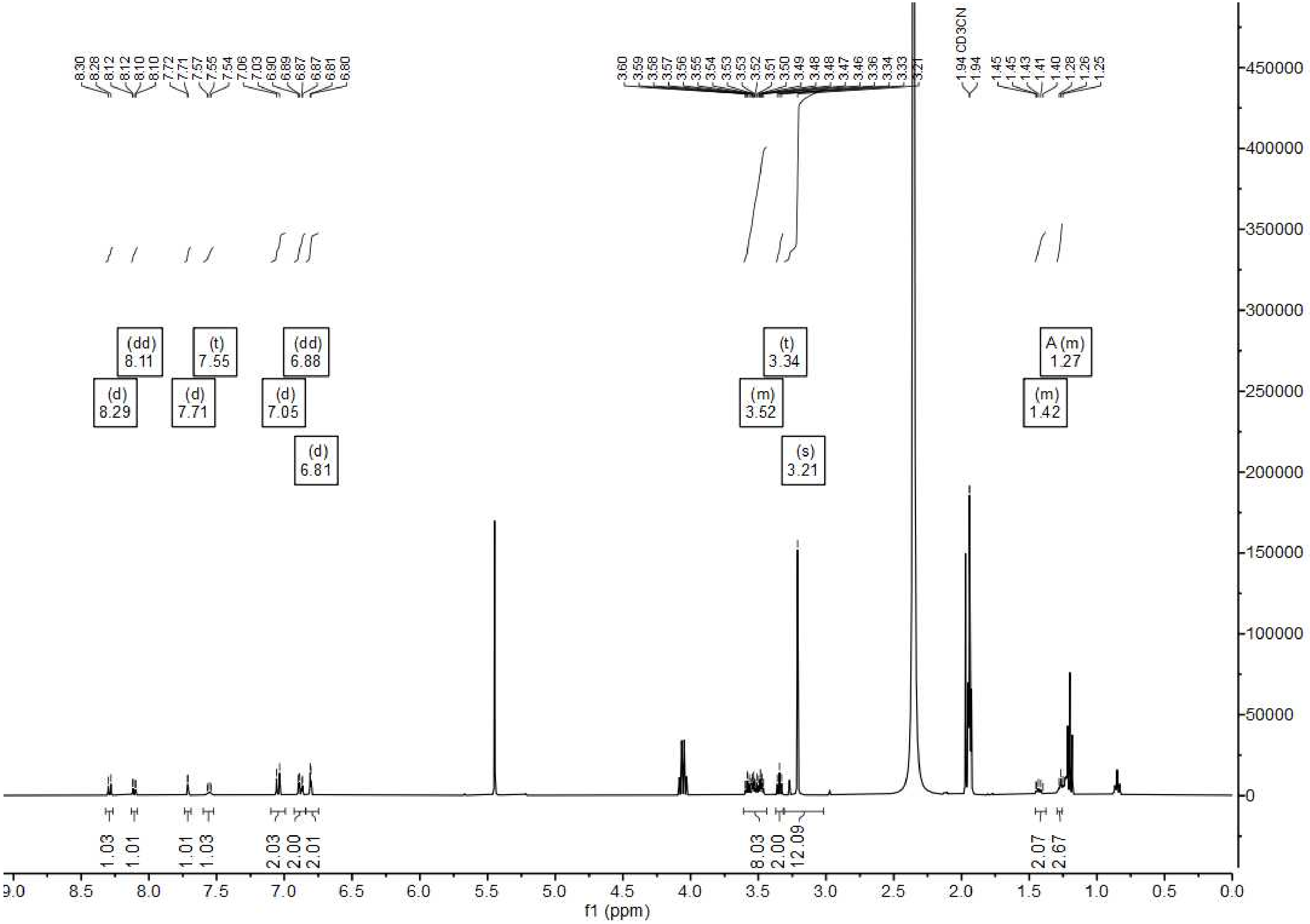

**Spectrum 9.** $^{1}$H-NMR ($CD_3CN$) of TMR-Hy4 (**17**).
